# Caveolar Compartmentalization of Pacemaker Signaling is Required for Stable Rhythmicity of Sinus Nodal Cells and is Disrupted in Heart Failure

**DOI:** 10.1101/2024.04.14.589457

**Authors:** Di Lang, Haibo Ni, Roman Y. Medvedev, Fang Liu, Claudia P. Alvarez-Baron, Leonid Tyan, Daniel G.P. Turner, Aleah Warden, Stefano Morotti, Thomas A. Schrauth, Baron Chanda, Timothy J. Kamp, Gail A. Robertson, Eleonora Grandi, Alexey V. Glukhov

## Abstract

**Background:** Heart rhythm relies on complex interactions between electrogenic membrane proteins and intracellular Ca^2+^ signaling in sinoatrial node (SAN) myocytes; however, mechanisms underlying the functional organization of proteins involved in SAN pacemaking and its structural foundation remain elusive. Caveolae are nanoscale, plasma membrane pits that compartmentalize various ion channels and transporters, including those involved in SAN pacemaking, via binding with the caveolin-3 scaffolding protein, but the precise role of caveolae in cardiac pacemaker function is unknown. Our objective was to determine the role of caveolae in SAN pacemaking and dysfunction (SND).

**Methods:** Biochemical co-purification, *in vivo* electrocardiogram monitoring, *ex vivo* optical mapping, *in vitro* confocal Ca^2+^ imaging, and immunofluorescent and electron microscopy analyses were performed in wild type, cardiac-specific caveolin-3 knockout, and 8-weeks post-myocardial infarction heart failure (HF) mice. SAN tissue samples from donor human hearts were used for biochemical studies. We utilized a novel 3-dimensional single SAN cell mathematical model to determine the functional outcomes of protein nanodomain-specific localization and redistribution in SAN pacemaking.

**Results:** In both mouse and human SANs, caveolae compartmentalized HCN4, Ca_v_1.2, Ca_v_1.3, Ca_v_3.1 and NCX1 proteins within discrete pacemaker signalosomes via direct association with caveolin-3. This compartmentalization positioned electrogenic sarcolemmal proteins near the subsarcolemmal sarcoplasmic reticulum (SR) membrane and ensured fast and robust activation of NCX1 by subsarcolemmal local SR Ca^2+^ release events (LCRs), which diffuse across ∼15-nm subsarcolemmal cleft. Disruption of caveolae led to the development of SND via suppression of pacemaker automaticity through a 50% decrease of the L-type Ca^2+^ current, a negative shift of the HCN current (*I*_f_) activation curve, and a 40% reduction of Na^+^/Ca^2+^-exchanger function, along with ∼2.3-times widening of the sarcolemma-SR distance. These changes significantly decreased the SAN depolarizing force, both during diastolic depolarization and upstroke phase, leading to bradycardia, sinus pauses, recurrent development of SAN quiescence, and significant increase in heart rate lability. Computational modeling, supported by biochemical studies, identified NCX1 redistribution to extra-caveolar membrane as the primary mechanism of SAN pauses and quiescence due to the impaired ability of NCX1 to be effectively activated by LCRs and trigger action potentials. HF remodeling mirrored caveolae disruption leading to NCX1-LCR uncoupling and SND.

**Conclusions:** SAN pacemaking is driven by complex protein interactions within a nanoscale caveolar pacemaker signalosome. Disruption of caveolae leads to SND, potentially demonstrating a new dimension of SAN remodeling and providing a newly recognized target for therapy.

## Introduction

Sinus node dysfunction (SND) is associated with abnormal impulse formation and propagation in the sinoatrial node (SAN), the primary pacemaker of the heart. On the electrocardiogram, SND usually manifests as sinus bradycardia, sinus arrest, or sinoatrial block, sometimes accompanied by supraventricular tachyarrhythmias (“tachy-brady” syndrome) and irregular prolonged pauses between heartbeats.^1^ However, the molecular and cellular mechanisms underlying SND are not completely understood, with one of the most critical limitations in treating SND being our incomplete understanding of structural and functional organization of cardiac pacemaking, including complex interactions between the components of the SAN pacemaker system. Recent studies^2,3^ have demonstrated a crucial role of rhythmic intracellular Ca^2+^ oscillations produced by the sarcoplasmic reticulum (SR) Ca^2+^ release channels, ryanodine receptors (RyR2s), in both the generation and autonomic regulation of pacemaker automaticity. These Ca^2+^ oscillations represent subsarcolemmal, local Ca^2+^ release (LCR) events that occur during the late phase of the spontaneous diastolic depolarization and activate Na^+^-Ca^2+^ exchanger (NCX1) inward current (*I*_NCX_). *I*_NCX_ significantly accelerates diastolic membrane depolarization, initiated by a hyperpolarization-activated *I*_f_ current that progressively declines over time, which triggers the activation of the L-type Ca^2+^ current (*I*_Ca,L_) and generates SAN action potential (AP). While it has been suggested that stable SAN pacemaking relies on the well-coordinated, Ca^2+^-mediated spatiotemporal synchronization between SR proteins (i.e., LCRs, referred to as an intracellular “Ca^2+^ clock”) and sarcolemmal proteins (including NCX1 and L-type Ca^2+^ channels, LTCC, envisioned as a “membrane clock”),^4-6^ the mechanisms underlying their functional organization and associated structural foundation remain elusive.

Whereas ventricular and atrial myocytes possess a transverse-axial tubular system that closely positions sarcolemmal and SR membranes within a ∼12-nm wide dyadic space and provides a structural foundation for excitation-contraction coupling, pacemaker cells lack these surface membrane extensions.^7^ Instead, SAN cells (SANCs) are enriched with caveolae structures, with rabbit SANCs exhibiting a caveolar density twice that of atrial myocytes and four to five times that of ventricular myocytes, as estimated from electron microscopy micrographs.^8^ Caveolae are small (50 – 100 nm) invaginations of the plasma membrane rich in cholesterol and sphingolipids, which are defined by their principal structural protein, caveolin. Caveolin-3 (Cav-3) is the specific and predominant caveolin isoform expressed in muscle^9^ and acts as a scaffolding protein by compartmentalizing and concentrating signaling molecules within caveolae.^4,10,11^ Moreover, through direct protein-protein interaction, Cav-3 can modulate the biophysical properties of associated ion channels and organize large macromolecular signaling complexes,^12-14^ contributing to nanodomain-specific regulation of cardiac physiology and pathophysiological remodeling and arrhythmogenesis.^11,15-17^ Disruption of caveolar nanodomains and/or downregulation or malfunction of Cav-3 were observed in various pathological conditions that are concomitantly associated with SND, including heart failure (HF),^18^ atrial fibrillation,^19^ and hypertension.^20^ Clinically, patients with *CAV3* genetic variants often exhibit symptoms such as sinus bradycardia, elevated heart rhythm lability, and cardiac arrest in sleep.^21^ Nevertheless, the role of caveolae in SAN pacemaking and SND remains to be elucidated.

Here, we performed a detailed multi-level electrophysiological, biochemical, and computational analysis of cardiac pacemaker organization, linking SAN ultrastructure, protein expression, and nanodomain-specific localization as well as protein-protein interaction, to stable rhythmic activity of the SAN *in vivo*, *ex vivo*, and *in vitro*. Specifically, our results demonstrate a key role of Cav-3 in organizing a macromolecular pacemaker complex and supporting functional coupling between NCX1, LTCC, and RyR2 molecules across a ∼15-nm caveolar subspace. This coupling is critically required for synchronization between the intracellular Ca^2+^ signaling and membrane excitability to generate pacemaker APs and support a stable, regular SAN rhythm. Further, we show that disruption of caveolae nanodomains in a cardiac-specific Cav-3 conditional knock-out (Cav-3KO) mouse model and in an 8-week post-myocardial infarction mouse HF model impairs the synchronization of the pacemaker components and leads to LCR- to-AP failure, sinus pauses, increased heart rate lability, tachycardia-bradycardia paroxysms, and enhanced atrial ectopy. The integrative experimental methods and simulation approaches presented here provide a comprehensive understanding of how disease-induced remodeling at the nanoscale level is manifested into dysfunction at the cellular and organ levels. Our findings indicate that caveolae represent a new dimension of SAN remodeling and could be considered a potential therapeutic target to restore SAN functioning in heart disease by either preventing the degradation or promoting the restoration of SAN cytoarchitecture.

## Methods

All experiments were conducted in accordance with the National Institutes of Health Guide for the Care and Use of Laboratory Animals (NIH Pub. No. 80-23). All methods and protocols used in these studies have been approved by the Animal Care and Use Committee of University of Wisconsin-Madison (U.S.A.) following the Guidelines for Care and Use of Laboratory Animals published by NIH (publication No. 85-23, revised 1996). All animals used in this study received humane care in compliance with the Guide for the Care and Use of Laboratory Animals. Human heart collection protocols were approved by the University of Wisconsin Institutional Review Board.

The data, methods used in the analysis (including program code for SAN modeling or script for caveolin binding motif screening), and materials used to conduct the research will be made available to any researcher for purposes of reproducing the results. For a detailed description of methods, see Expanded Methods in the Supplementary Materials.

## Results

### Caveolar macromolecular pacemaker complex

Using transmission electron microscopy (TEM) imaging, we found that mouse SANCs are highly abundant in caveolae (Fig. 1A and Figs. S1 and S2), with an average caveolae density twice as high than in atrial myocytes and four times higher than in ventricular myocytes (Fig. S3), consistent with previous findings in rabbit^8^ and pig SANCs.^22^ Caveolae are positioned adjacent to the subsarcolemmal SR, effectively reducing the distance between the SR membrane and the nearest cell surface membrane: 15 ± 1 nm to the nearest caveolar membrane, compared to 28 ± 3 nm to the closest non-caveolar membrane (n=76 cells from 5 mice, *P*<0.001; Fig. 1A). Although RyR2s exhibited a regular striated expression pattern throughout the cell, only subsarcolemmal RyR2s were associated with Cav-3, indicating the absence of transversal-axial tubular system in SANCs.

**Figure 1.**
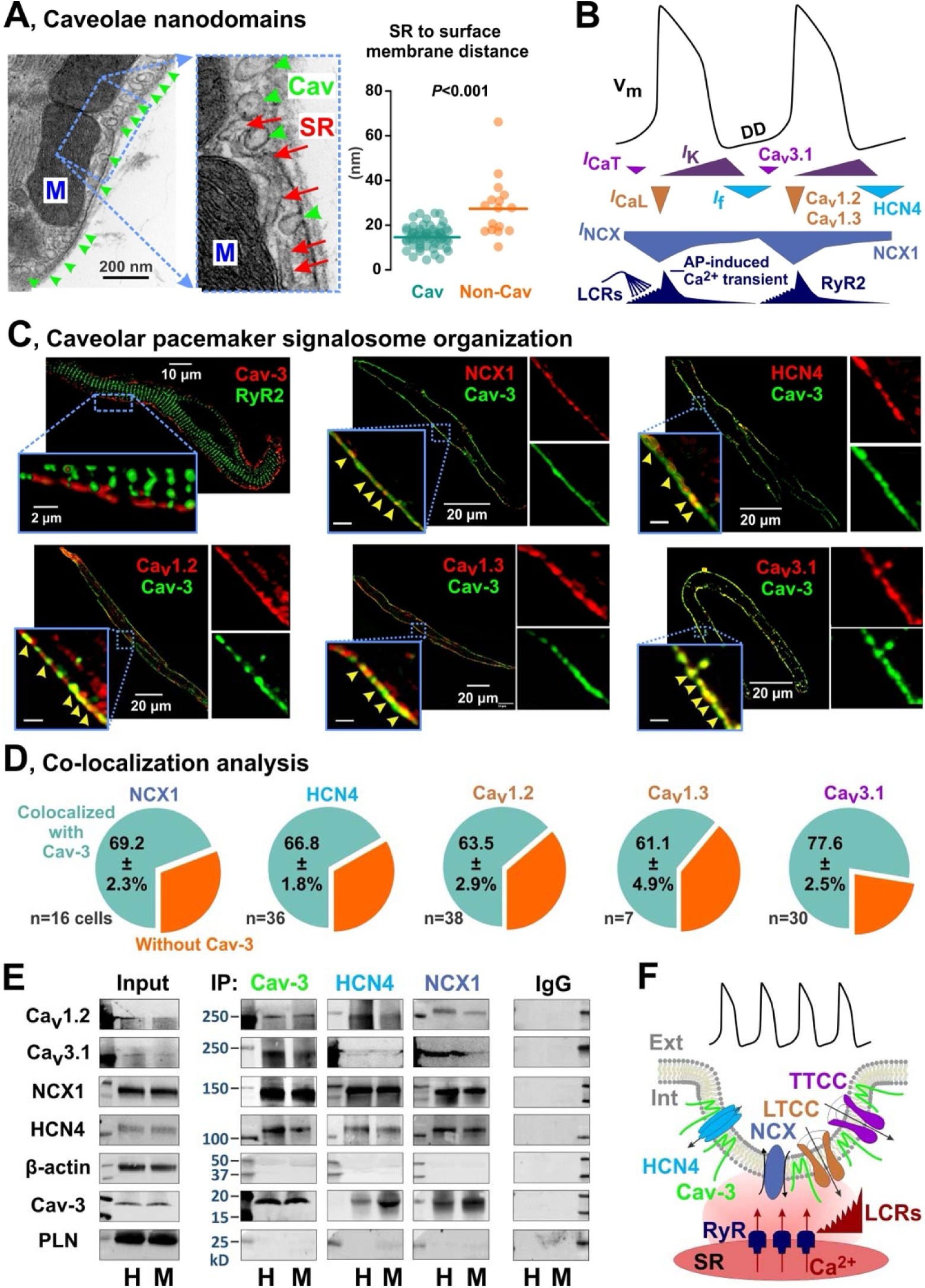
Caveolar macromolecular pacemaker complex in SANCs. **(A)** TEM photograph (*left*) of SANC demonstrates a highly dense distribution of caveolae structures (Cav, *green arrows*) that are located adjacent to subsarcolemmal SR (*orange arrows*). n=76 cells from 5 mice; each dot represents caveolae density from one TEM cell. Caveolae significantly shorten the distance between the SR membrane and the nearest sarcolemmal (i.e., caveolar (Cav, green) or non-caveolar (Non-Cav, orange)) membranes. *P*-value by Student’s t-test. **(B)** Typical SAN AP and the timing of membrane (V_m_) and Ca^2+^-clock components. DD – diastolic depolarization. Modified from Lakatta et al.^2^ with permission. **(C)** Immunofluorescent labeling of the membrane and Ca^2+^ clock components in mouse SANCs. Yellow arrowheads in inserts show colocalization clusters. **(D)** Colocalization analysis for sarcolemmal pacemaker proteins with regard to their association with Cav-3 in mouse SANCs. **(E)** Mouse (*M*) and human (*H*) SAN tissues were subjected to immunoprecipitation with anti-Cav-3, anti-NCX or anti-HCN4 antibodies, and the immunoprecipitates were analyzed by immunoblotting for sarcolemmal pacemaker proteins. Phospholamban (PLN) and β-actin were used as non-caveolar control proteins. Results are representative of four different experiments. Additional blots, antibodies testing and negative controls are shown in Supplemental Figs. S5 and S6. **(F)** Proposed model of caveolar pacemaker signalosome which concentrates components of the membrane clock to caveolae positioning them in a close proximity to subsarcolemmal RyRs and thus enabling their efficient spatial-temporal synchronization.

To elucidate the contribution of caveolae in the localization of electrogenic membrane proteins involved in SAN pacemaking (Fig. 1B), we performed co-immunolabeling for Cav-3 and HCN4 channels, NCX1, L-type Ca^2+^ channels Ca_v_1.2 and Ca_v_1.3, and T-type Ca^2+^ channels Ca_v_3.1 (Fig. 1C). Structural proximity of caveolar and SR membranes was confirmed by co-immunolabeling of caveolar scaffolding protein Cav-3 and subsarcolemmal RyR2 (Fig. 1C). Subsequent colocalization analysis revealed that 60-80% of pacemaker proteins are associated with Cav-3 (Fig. 1D). To determine whether Cav-3 is directly associated with the components of the membrane clock, we performed a series of immunoprecipitation experiments. In both mouse and human SAN tissue (Fig. S4) lysates, we found that Ca_v_1.2, Ca_v_3.1, NCX1 and HCN4 co-immunoprecipitated with Cav-3, but not with the contractile protein β-actin and the SR protein phospholamban, used as negative controls (Fig. 1E and Fig. S5 and S6). The presence of such a macromolecular pacemaker complex was further supported by subjecting mouse and human SAN lysates to immunoprecipitation with anti-NCX1 and anti-HCN4. Triple immunofluorescent staining for Cav-3, HCN4, and Ca_v_1.2 (Fig. S7) clearly demonstrated the close association of these proteins, with a distinct punctate expression pattern on the surface membrane, likely indicative of caveolae structures.^4,10^ These data were further corroborated by the presence of complementary caveolin binding motifs identified in the amino acid sequences of mouse and human HCN4, Ca_v_1.2, Ca_v_1.3, Ca_v_3.1 and NCX1 proteins (Fig. S8). These results provide the basis for our hypothesis that caveolae provide a structural foundation for the coupled-clock pacemaker system by concentrating membrane-clock components within the caveolar membrane, and positioning them near the SR membrane, thus providing the spatial-temporal coupling between the membrane- and Ca^2+^-clocks (Fig. 1F).

### Downregulation of caveolae leads to SND

To determine whether caveolae play a functional role in SAN pacemaking, we utilized Cav-3KO mice.^12,24^ As we have previously shown, cardiac-specific conditional deletion of *CAV3* results in significant downregulation of Cav-3 mRNA and protein expression levels as well as caveolae structures in ventricular^12^ and atrial^17^ myocardium. Here, we further show that deletion of *CAV3* significantly decreased Cav-3 mRNA and protein expression levels and caveolae density in the SAN (Fig. S9). *In vivo*, Cav-3KO mice exhibited slower heart rates, sinus pauses, and significantly increased heart rate variability when compared with their litter-mate controls (Fig. 2A-C). This was accompanied by paroxysms of alternating periods of tachycardia-bradycardia and enhanced atrial ectopy (Fig. 2A, *bottom panel*), a hallmark of SND. Enhanced atrial ectopic activity was characterized by profound beat-to-beat alterations of ECG P-wave morphology, including fractionated, premature, and inverted P-waves (red arrows in the insertion in Fig. 2A), along with dramatically increased beat-to-beat cycle length (CL) lability (Fig. 2C).

**Figure 2.**
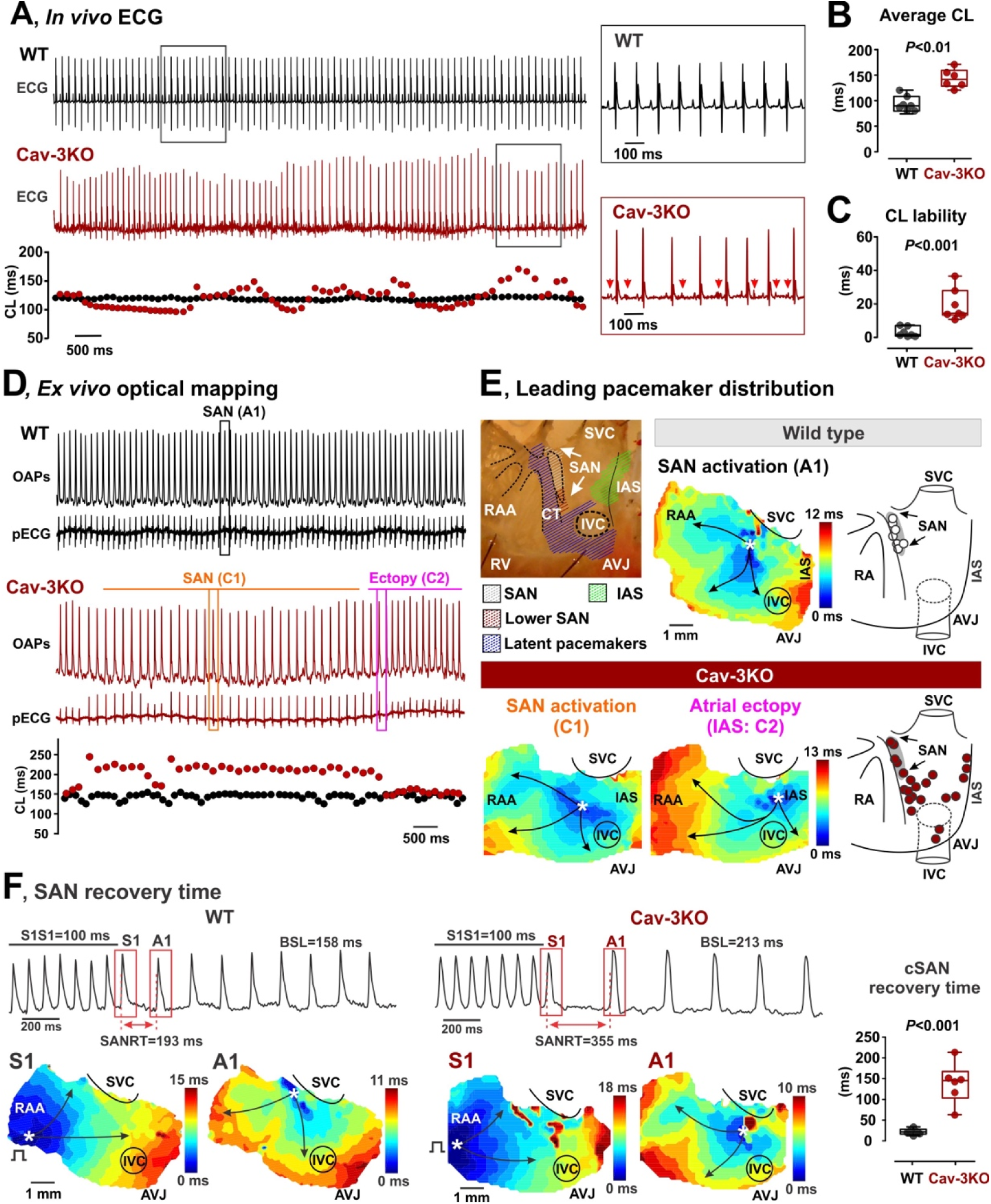
SND in cardiac-specific conditional Cav-3KO mice. **(A)** *In vivo* ECG telemetry demonstrates highly irregular heart rate in Cav-3KO versus WT mice. Note premature and fractionated P-waves highlighted by red arrows in the insertion. **(B and C)** Averaged cycle length (CL) (B) and CL lability (C) measured in WT (n=7) and Cav-3KO (n=6) mice. **(D)** *Ex vivo* optical mapping from isolated spontaneously beating SAN preparations demonstrates SND phenotype in Cav-3KO mice, as seen from representative optical APs (OAPs) and pseudo-ECG (pECG) traces. Below, corresponding beat-to-beat CLs are shown. SAN and ectopic beats highlighted by boxes were used for atrial activation reconstruction shown in panel E. **(E)** Representative atrial activation contour maps reconstructed for WT (SAN activation, A1, *top panel*) and Cav-3KO mouse (SAN activation, C1, and activation from an ectopic pacemaker, C2). On the right of the activation maps, distributions of the leading pacemaker locations summarized from all WT and Cav-3KO mice are shown. SVC and IVC, superior and inferior vena cava; RAA, right atrial appendage; RV, right ventricle; CT, crista terminalis; IAS, inter-atrial septum; AVJ, atrioventricular junction. **(F)** SAN recovery time in WT and Cav-3KO mice. Representative optical maps obtained at the last pacing stimulus (S1) and at the first spontaneous post-pacing atrial beat (A1) are shown. On the right, summarized data for SAN recovery time corrected to basic heart rate is shown. *P*-values by Student’s t-test.

*Ex vivo* fluorescent optical mapping of APs from isolated SAN preparations confirmed SND phenotype in Cav-3KO mice, manifesting in alternating periods of tachycardia and bradycardia with increased atrial ectopy (Fig. 2D). Furthermore, optical mapping enabled localization of the leading pacemaker, which was consistently identified within the SAN in WT mice, with its location exhibiting greater variability in preparations from Cav-3KO mice (Fig. 2E and Figs. S10 and S11). Importantly, in Cav-3KO SAN preparations, the periods of bradycardia were driven by leading pacemakers located within the anatomically defined SAN region, while periods of atrial tachycardia were associated with a shift of the leading pacemaker outside of the SAN to multiple ectopic foci located within “an extensive distributed system” of atrial pacemakers^25,26^ (Fig. 2E). In addition to the elevated lability of spontaneous beating rate, Cav-3KO mice demonstrated a significantly prolonged heart rate-corrected SAN recovery time (Fig. 2F), which further supports their SND phenotype.

### Caveolae disruption impairs spatial and temporal coupling between Ca^2+^ and V_m_ clocks resulting in pacemaker dysfunction

To determine the mechanisms by which the downregulation of Cav-3 and caveolae contribute to SND, we performed confocal Ca^2+^ imaging on isolated SANCs to evaluate properties of both “Ca^2+^ clock” and “membrane clock” components of the pacemaker system as well as their functional coupling. The line scan was oriented along the long axis of the SANCs just beneath the sarcolemma to record local Ca^2+^ release events (LCRs)^5,27,28^ that represent a crucial component of the intracellular Ca^2+^ clock.^2,5^ WT-SANCs exhibited stable regular spontaneous beating with negligible beat-to-beat CL variations in both AP-induced CaTs and LCRs (Fig. 3A, *upper panel*, and Fig. 3A1). In contrast, Cav-3KO*-*SANCs displayed the SND phenotype observed in both *in vivo* ECG telemetry and *ex vivo* optical mapping (Fig. 2), characterized by highly irregular AP-induced CaTs and prolonged SAN pauses (Fig. 3A, *low panel*, and Fig. 3A2). In Cav-3KO*-*SANCs, irregular CaTs (blue dots in Poincaré beat-to-beat CL plots on the right of Figs. 3A1-2) were accompanied by regular rhythmic LCRs (red dots in Poincaré plots) which were less efficient in triggering APs, thus resulting in an irregular rhythm. In contrast, WT-SANCs showed a 1:1 coupling between LCRs and AP-induced global CaTs enabling a regular rhythm.

**Figure 3.**
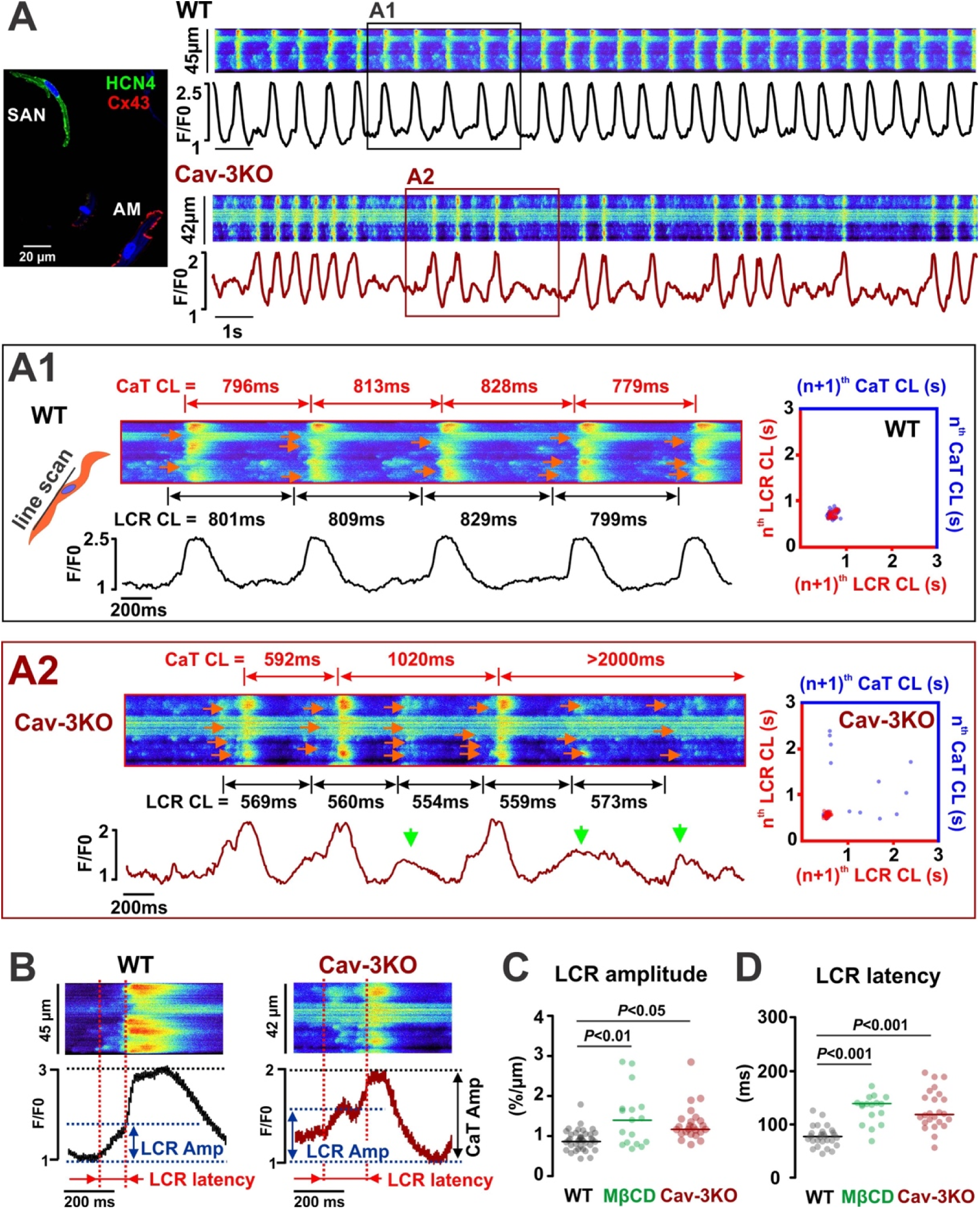
Functional uncoupling between V_m_ and Ca^2+^ clocks in Cav-3KO*-*SANCs. **(A)** *In vitro* confocal imaging of spontaneous [Ca^2+^]*_i_*in WT and Cav-3KO SANCs identified as spontaneously beating, HCN4-positive and Cx43-negative cells located within the SAN region (AM: atrial myocyte). Selected segments of representative traces are shown enlarged below for WT (A1) and Cav-3KO (A2) cells. Confocal line scan was positioned beneath the sarcolemma along the long cell axis to record local Ca^2+^ releases (LCRs, indicated by red arrows on the scan) together with CaT. LCR-V_m_ uncoupling (A2) was characterized by unsuccessful CaT (green arrows) accompanied by regular rhythmic LCRs. LCR-to-CaT dissociation is evident from the corresponding beat-to-beat LCR and CaT CL Poincaré scatter plots (*right*). **(B)** Enlarged representative LCR-to-CaT events showing LCR amplitude (the amount of Ca^2+^ released during LCR) and LCR latency (LCR-to-CaT time). Summarized data for LCR amplitude **(C)** and latency **(D)** is shown for WT (n=34 cells), MβCD-WT (n=17 cells) and Cav-3KO (n=24 cells) SANCs. *P*-values by one-way ANOVA with Bonferroni correction.

Importantly, we were able to induce a similar SND phenotype in WT-SANCs treated with 10 mM MβCD, a cholesterol binder that disrupts caveolae structures.^16^ Cav-3KO and MβCD-WT SANCs exhibited a significantly prolonged LCR latency (i.e., the time period from LCR to the beginning of CaT, Fig. 3B,D) and augmented amplitude (i.e., the amount of Ca^2+^ released during each event, Fig. 3B,C). The former indicates an increased distance between LCR generating RyR release sites on the SR membrane and NCX-mediated Ca^2+^ extrusion sites on the sarcolemma, which would weaken *I*_NCX_ activation and lead to LCR-to-AP failure and SAN pauses. The increased LCR amplitude suggests larger LCRs, potentially mediated by either higher SR Ca^2+^ content due to preceding LCR-to-AP failures, hypoactive RyRs, and/or a longer release time during the prolonged LCR-to-AP period observed in Cav-3KO and MβCD-WT SANCs. This was further accompanied by an increase in spontaneous Ca^2+^ spark activity in Cav-3KO-SANCs (Fig. S12).

We found that downregulation of caveolae led to a significantly increased distance between sarcolemmal and SR membranes from 18 ± 1 nm in WT to 41 ± 3 nm in Cav-3KO SANCs (Fig. 4A), supporting the notion of NCX1-RyR2 uncoupling. Immunogold labeling combined with TEM confirmed the localization of NCX1 within caveolae in WT-SANCs, with some NCX1 clusters located in close proximity to the SR membrane (highlighted by a pink shaded area, Fig. 4B), akin to the SR-caveolae membrane arrangements shown in Fig. 1A. On the other hand, while NCX1 clusters were still observable on the sarcolemma of Cav-3KO*-*SANCs, they were not associated with caveolae structures. Furthermore, immunofluorescent analysis performed for NCX1 and Cav-3 colocalization (Fig. 4C) as well as NCX1/Cav-3 proximity ligation assay measurements (PLA, Fig. 4D and Fig. S13) confirmed a dramatic decrease in NCX1-Cav-3 interaction in Cav-3KO*-*SANCs. Altogether, these findings suggest that loss of caveolae nanodomains disrupts the functional coupling between LCR (Ca^2+^-clock) and AP (membrane clock) by increasing the physical distance between NCX1 and RyR2s.

**Figure 4.**
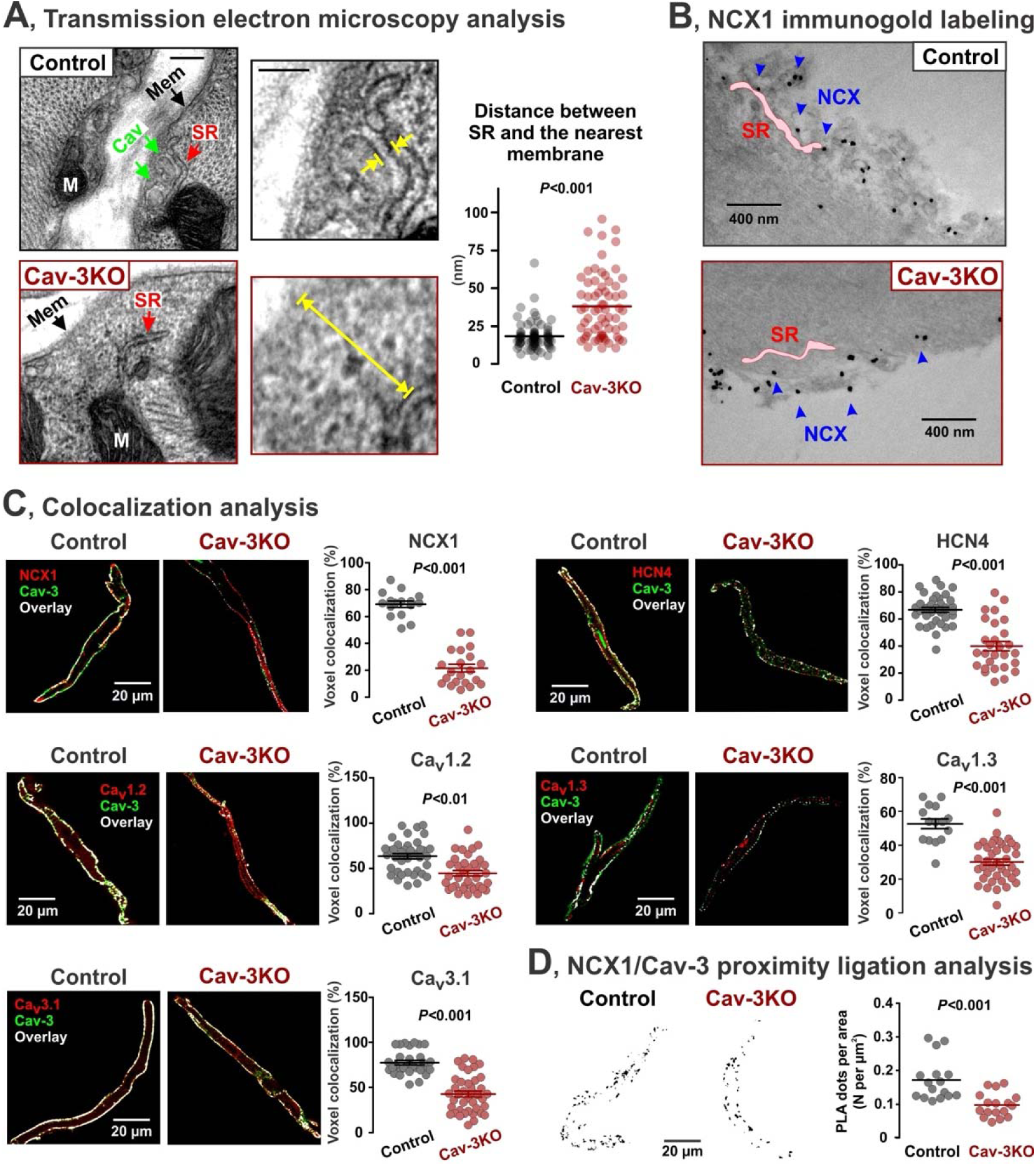
Disruption of caveolar macromolecular pacemaker complex in Cav-3KO mice. **(A)** Distance (indicated be yellow arrows) between the SR membrane (red arrow) and the nearest sarcolemmal membrane (green arrows for caveolar membrane, Cav, and black arrows for non-caveolar sarcolemmal membrane, Mem). n=76 images from 5 control mice and n=66 images from 4 Cav-3KO mice. **(B)** NCX1 immunogold staining confirms that NCX1 is released from caveolae and distributed in the membrane surface in Cav-3KO SANCs, where NCX1 molecules are located further from SR. **(C)** Colocalization analysis for Cav-3 and NCX1 (16/21 control/Cav-3KO cells, 3-5 mice/group), Cav-3 and HCN4 (36/28 control/Cav-3KO cells, 3-5 mice/group), Cav-3 and Ca_v_1.2 (38/36 control/Cav-3KO cells, 3-5 mice/group), Cav-3 and Ca_v_1.3 (15/40 control/Cav-3KO cells, 3-5 mice/group), and Cav-3 and Ca_v_3.1 (30/40 control/Cav-3KO cells, 3-5 mice/group). Representative confocal images show Cav-3 expression in green, specific proteins of interest in red, and overlayed signal in white color. **(D)** *Left:* Representative pictures of proximity ligation assay between NCX1 and Cav-3 in control and Cav-3KO cells. *Right:* Comparison of the puncta density formed by the association of NCX1 with Cav-3 in control (n=16 cells) and Cav-3KO (n=17 cells) SANCs, 3 mice/group. *P*-values by Student’s t-test.

### Disruption of caveolar macromolecular pacemaker complex in Cav-3KO mice leads to SND

To investigate the impact of caveolae disruption on the localization of other pacemaker proteins, we conducted co-immunolabeling for Cav-3 and key electrogenic proteins involved in cardiac pacemaking, including HCN4, Ca_v_1.2, Ca_v_1.3, and Ca_v_3.1 in WT and Cav-3KO SANCs (Fig. 4C). Colocalization analysis revealed a significant decrease in voxel colocalization for all the protein pairs, providing evidence of the redistribution of caveolar pacemaker proteins due to caveolae disruption. To assess the effects of the observed redistribution of pacemaker proteins on SAN pacemaking, we developed a novel mathematical model of the SANC that integrates mouse SAN electrophysiology with a detailed description of subcellular stochastic properties of individual SR Ca^2+^ release units (Fig. 5A, a detailed model description is available in the Supplementary Materials). The model consisted of 2,176 (34 × 8 × 8) Ca^2+^ release units (CRUs). Each CRU comprises 5 Ca^2+^ compartments: cytosolic, submembrane, cleft space, network SR, and junctional SR. The peripheral CRUs are coupled to external membranes, whereas central CRUs are not coupled to external membranes and have fewer RyRs than peripheral CRUs. In the peripheral CRUs, the caveolae-associated regions encompass the whole cleft space and the majority of the submembrane space. Our baseline model displayed stable and rhythmic spontaneous beating with rhythmic LCRs preceding AP-induced CaT (Fig. 5B, *baseline model 1*), in agreement with the phenotype of WT-SANCs (Fig. 3A).

**Figure 5.**
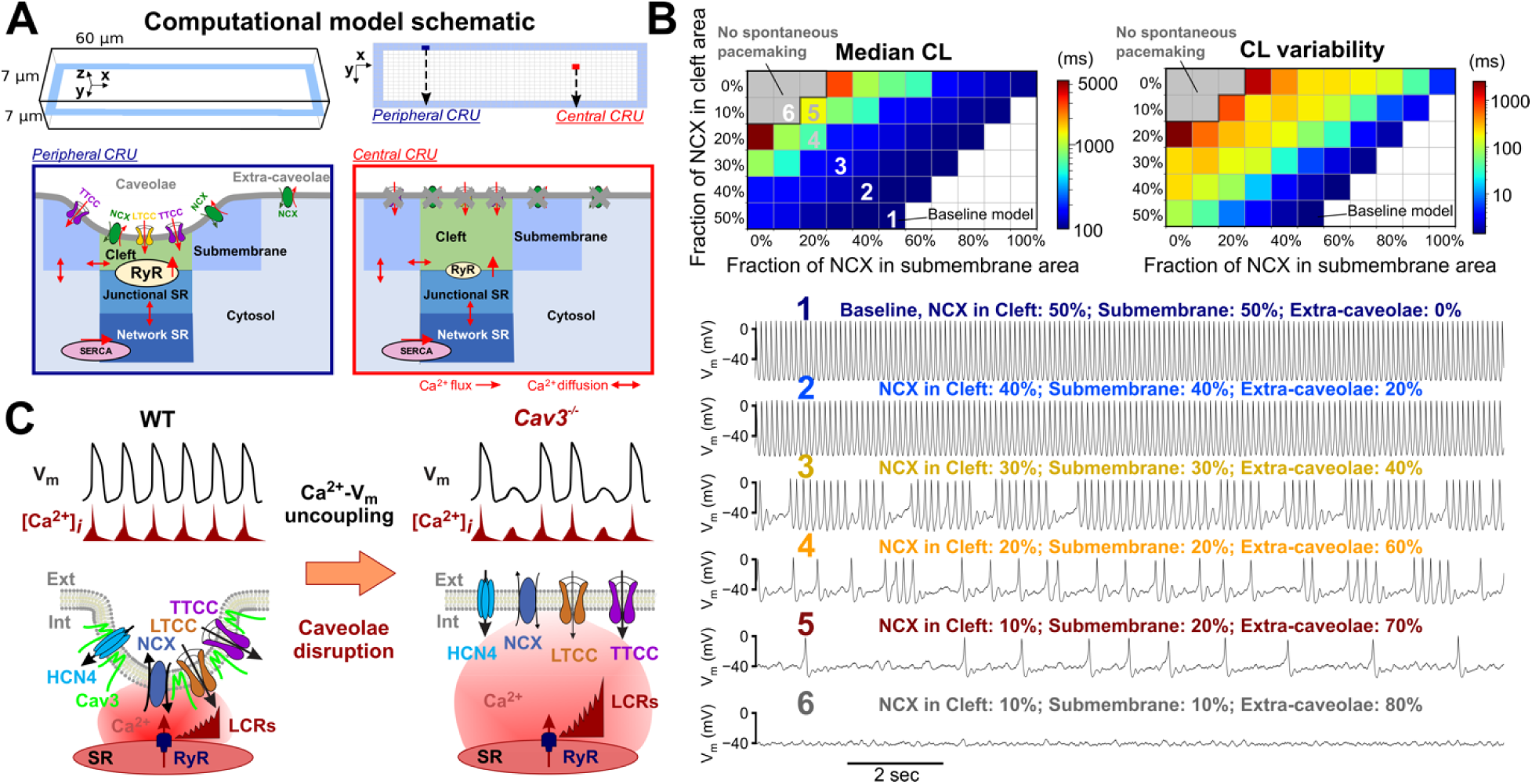
Computational modeling of compartmentalized mouse SAN myocyte electrophysiology. **(A)** Schematic of the 3D V_m_ and subcellular Ca^2+^ handling model for a mouse SANC. **(B)** Disruption of SAN pacemaking rhythm occurs over a wide range of NCX1 redistribution patterns away from portions of the sarcolemma facing the cleft and submembran Ca^2+^ compartments. *Top*: Median CL (*left*) and CL variability (*right*) with respect to varying fractions of NCX1 in the cleft space (Y-axis) and the submembrane area (X-axis); the remaining fraction of NCX1 was redistributed to face the cytosolic Ca^2+^ compartment (extra-caveolae). *Bottom*: Time courses of V_m_ for the baseline model and models with various degrees of NCX1 redistribution; numbers indicate the NCX1 distribution parameters marked in the top left panel. **(C)** Proposed spatial-temporal organization of caveolar macromolecular pacemaker complex and its disruption upon downregulation of caveolae nanodomains.

When simulating a gradually increased distance between Ca^2+^ release sites and NCX1-mediated Ca^2+^ removal sites (i.e., by partially relocating NCX1 proteins in the extra-caveolae membrane; Fig. 5B, *models 2-5*), we observed prolonged SAN pauses (i.e., an increase in a median CL, *left panel*) and highly irregular AP-induced CaTs (as characterized by significantly elevated CL variability compared to baseline model, *right panel*). These simulations recapitulate the experimentally observed SND phenotype *in* Cav-3KO mice (Fig. 3). A more significant redistribution of NCX1 from caveolar clefts to extra-caveolar membrane resulted in complete uncoupling between LCR and AP and led to loss of SAN excitability (Fig. 5B, *model 6*, absence of spontaneous activity as a gray area on a graph), similar to previous findings in NCX1 knock-out mice.^29^

Similar analysis performed for *I*_Ca,L_ (Fig. S14) and *I*_Ca,T_ (Fig. S15) currents did not reveal significant effects on SAN pauses and CL variability: while some occasional SAN exit blocks were observed, they did not lead to loss of SAN excitability or profound SAN pauses, as found with NCX1 redistribution. These results support the hypothesis that redistribution of NCX1 proteins from caveolae near Ca^2+^ release sites to distant membrane regions is crucial in the development of SND because it impairs the ability of NCX1 to be efficiently activated by LCRs and trigger APs. Notably, by simulating various degrees of NCX1 re-location from the release sites (Fig. 5B), we identified a parameter range that is permissive of this behavior, attesting to the robustness of our conclusions, within a spectrum of phenotypes ranging from normal AP generation to loss of SAN firing.

Besides protein redistribution, reduced expression of the pacemaker proteins and/or changes in associated ion currents could also induce SND. RT-qPCR profiling did not show any statistically significant changes in expression of Ca_v_1.2, Ca_v_1.3, Ca_v_3.1, NCX1, HCN1, HCN2, and HCN4 mRNA in Cav-3KO vs. WT mouse SAN tissues (Fig. 6A). Similarly, we did not observe changes in *I*_f_ current amplitude in Cav-3KO mice; however, we found a significant shift of the *I*_f_ activation curve to more negative voltages in Cav-3KO compared with WT-SANCs (Fig. 6B), which may indicate a reduced contribution of *I*_f_ to diastolic depolarization. Moreover, despite the unchanged expression of Ca_v_1.2 and Ca_v_1.3 mRNA, we revealed a significant decrease in atrial *I*_Ca,L_ current amplitude in Cav-3KO mice, while *I*_Ca,T_ amplitude was unchanged (Fig. 6C). Furthermore, we found a significant decrease in the NCX function in Cav-3KO*-*SANCs that was estimated from the caffeine-induced [Ca^2+^]*_i_* decay time (Fig. 6D) supporting a weakened coupling between V_m_ and Ca^2+^-clocks as it was found in *in vitro* (Fig. 3) and *in silico* (Fig. 5) experiments.

**Figure 6.**
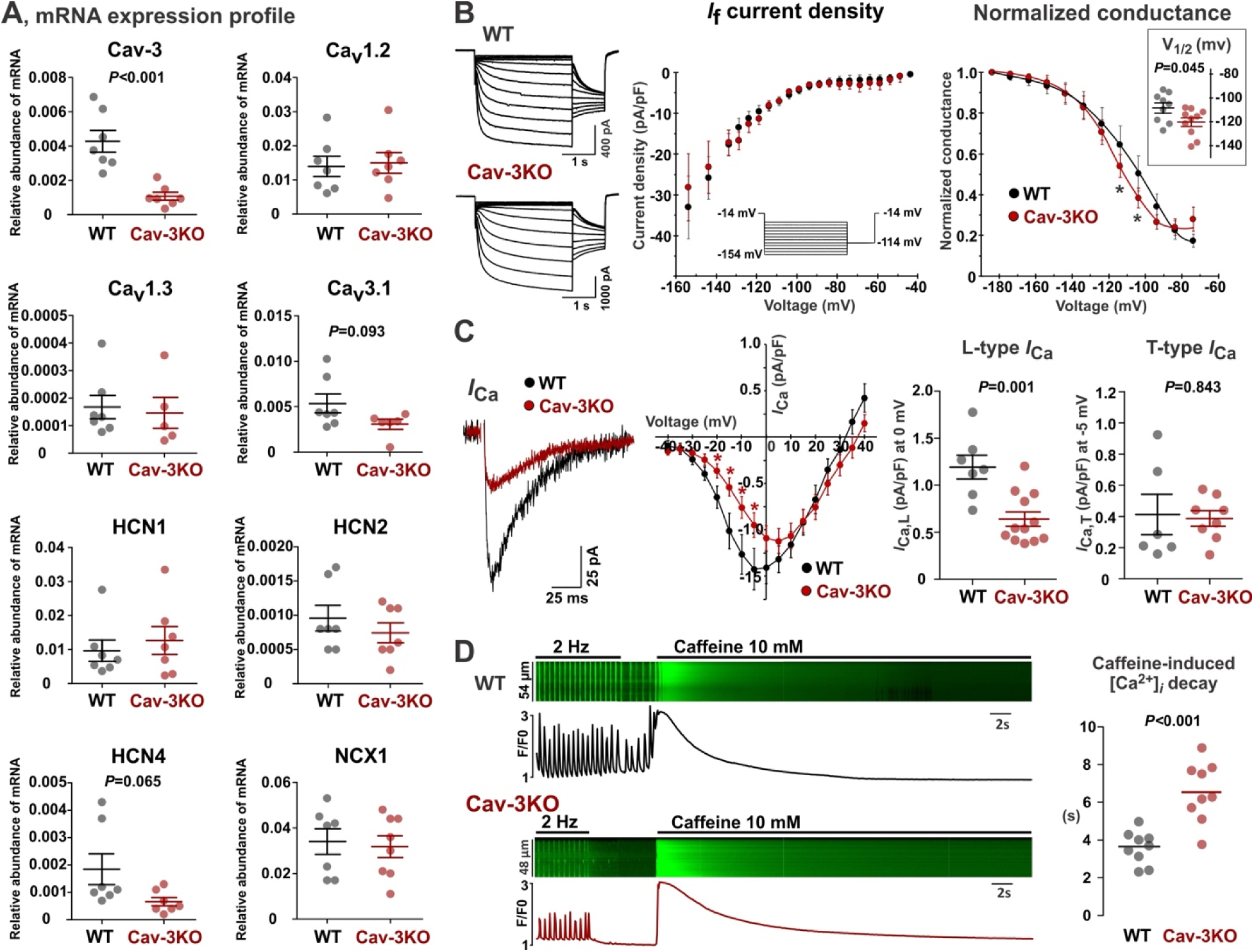
Molecular and functional remodeling in SAN pacemaker components in Cav-3KO mice. **(A)** mRNA expression profiling of the membrane clock proteins in WT (n=7) and Cav-3KO (n=8) SAN tissue samples. **(B)** Cav-3KO*-*SANCs display a similar *I*_f_ current density and a tendency for more hyperpolarized mean values of half-activation voltage (V_1/2_) compared to WT-SANCs (9/10 WT/Cav-3KO cells). **(C)** Atrial cardiomyocytes from Cav-3KO show a significant decrease in *I*_Ca,L_ density and no statistical change in *I*_Ca,T_ density (6/8 WT/Cav-3KO cells). **(D)** Cav-3KO*-*SANCs demonstrate a significant reduction in the NCX1 function estimated as a caffeine (10 mM) induced [Ca^2+^]*_i_* decay time (9 cells/group). *P*-values by Student’s t-test.

### SND: protein downregulation versus redistribution

Our biochemical and computational modeling studies point to the disruption of the macromolecular pacemaker complex and the redistribution of NCX1 and LTCC proteins from their caveolar location near SR Ca^2+^ release sites to distant membrane regions as key contributors to SND in Cav-3KO mice (Fig. 5B-C). At the same time, functional assessments revealed a significant downregulation of NCX1 and *I*_Ca,L_ function, but not expression (Fig. 6). Since both redistribution and diminished activities of NCX1 and LTCC can independently contribute to pacemaker abnormalities, to independently test redistribution vs. diminished activities we performed *in vitro* confocal Ca^2+^ imaging (Fig. 7A) and *in silico* simulations (Fig. 7B) on SANCs during selective inhibition of components of the membrane clock. Inhibition of *I*_f_, *I*_Ca,L_, and *I*_Ca,T_ currents by 60% resulted in significant slowing of SAN pacemaker activity in both experiments and simulations (Fig. 7C and 7E). However, these decreases in spontaneous beating rate were not accompanied by an increased beat-to-beat CL variability, sinus pauses and SAN quiescence (Fig. 7D and 7F), in contrast to what was observed during caveolae disruption in both Cav-3KO and MβCD-WT SANCs (Fig. 3). At the same time, weakening NCX1 function recapitulated both a decreased beating rate and a significantly increased CL variability in both *in vitro* and *in silico* experiments (Fig. 7C-F). Complete inhibition of NCX1 (3 μM SEA0400) blocked the SAN firing by uncoupling APs from regular rhythmic LCRs (Fig. S16). Importantly, despite the lack of pacemaker activity, LCRs were observed during complete NCX1 inhibition but they failed to trigger APs, as we also found in Cav-3KO and MβCD-WT SANCs. These experiments confirm the pivotal role of NCX1-RyR2 localization in LCR-to-AP coupling across the ∼15-nm caveolar subspace.

**Figure 7.**
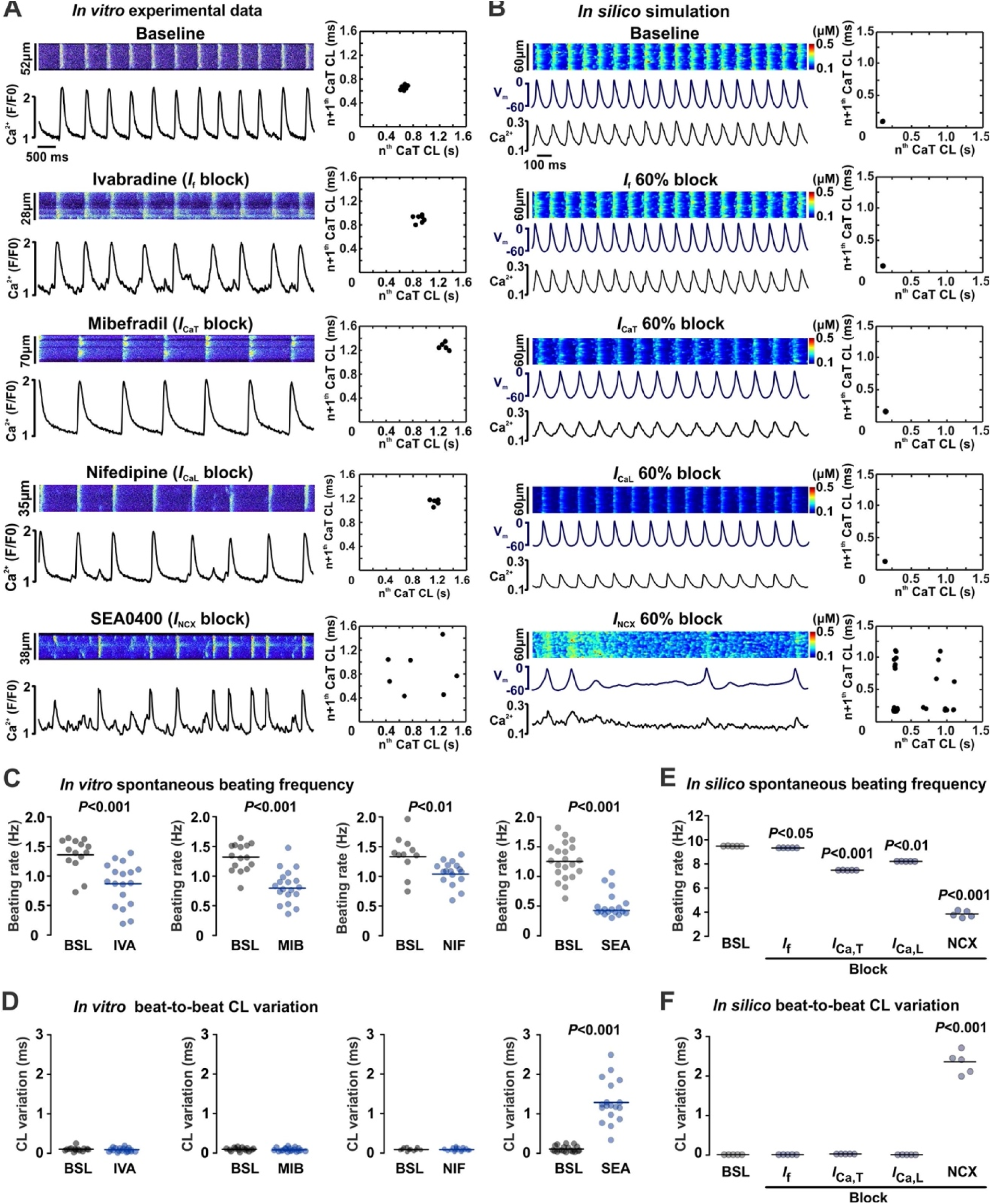
NCX1-mediated SND phenotype induced in WT-SANCs *in vitro* and *in silico*. Representative confocal (A) and simulated (B) CaTs for baseline condition and during selective inhibition of components of the membrane clock. For each condition, corresponding CaT CL Poincaré scatter plots are shown. For pharmacological inhibition, IC_50_ concentrations of corresponding blockers were used to partially block *I*_f_ (1 μM ivabradine, IVA, 14/18 baseline/treatment cells), *I*_Ca,T_ (1 μM mibefradil, MIB, 15/19 baseline/treatment cells), *I*_Ca,L_ (1 μM nifedipine, NIF, 11/17 baseline/treatment cells), or *I*_NCX_ currents (0.3 μM SEA0400, SEA, 21/18 baseline/treatment cells). N=5 for *in silico* simulations per condition. Summarized changes in spontaneous beating rate (panel C for *in vitro* and panel E for *in silico* experiments) and CL variation (panel D for *in vitro* and panel F for *in silico* experiments) are shown for each intervention. *P-*values by Student’s t-test for panels C and D, and one-way ANOVA with Bonferroni correction for panels E and F.

### SND in heart failure mice is associated with caveolae downregulation and uncoupling between membrane and Ca^2+^ clocks

SND is common in HF and linked to a reduction in functional SAN reserve.^30^ To address a possible role of caveolae/Cav-3 downregulation in SND in HF, we employed an 8-week post-myocardial infarction mouse model of HF and compared them with age-matched control mice (AMC; Fig. S17). Similar to Cav-3KO mice, HF mice displayed *in vivo* bradycardia (Fig. 8A), accompanied by a highly irregular heart rate, tachycardia-bradycardia-like arrhythmias, and enhanced atrial ectopy. This was further associated with a competitive shift of the leading pacemaker from the SAN to ectopic foci (Figs. 8B, S18-20) and an increased corrected SAN recovery time (Fig. 8B), similar to what we found in Cav-3KO mice (Fig. 2), confirming SND phenotype in HF mice.

**Figure 8.**
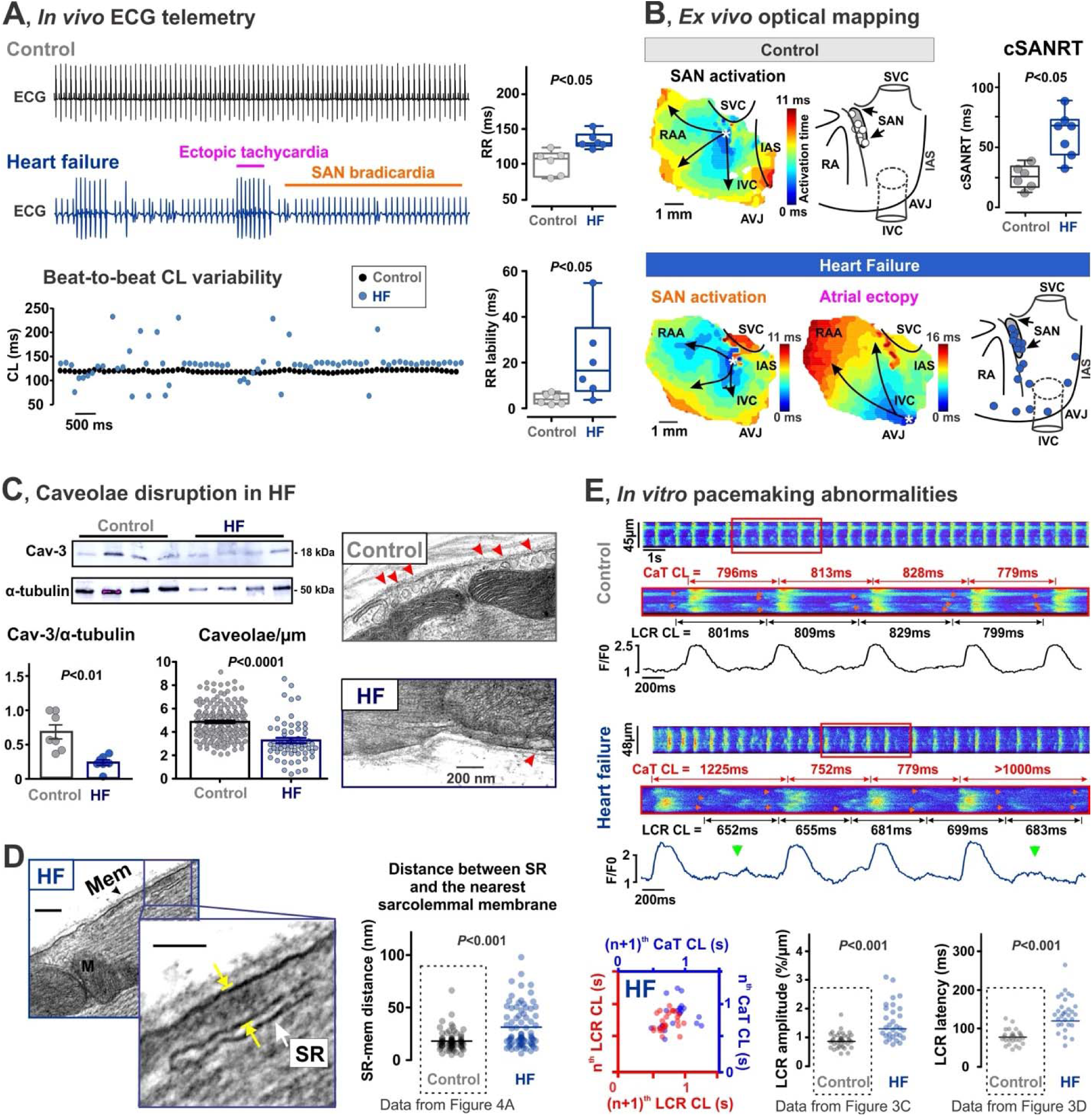
SND in HF mice is associated with caveolae downregulation and uncoupling between membrane (V_m_) and Ca^2+^ clocks. **(A)** *Left:* Representative *in vivo* ECG telemetry traces and corresponding beat-to-beat cycle length (CL) changes from control (age-matched wild type mice) and HF mice. *Right:* Average ECG RR interval and RR lability are shown for both groups. **(B)** Representative atrial activation maps are shown together with the distribution of leading pacemaker locations summarized from control (n=6) and HF (n=7) mice. On the right, SAN recovery time corrected to heart rate (cSANRT) is shown. Labels are same as in Figure 2. **(C)** Downregulation of Cav-3 protein (n=7/group) and caveolae during HF (n=183 cells from 5 control mice, and n=59 cells from 4 HF mice). **(D)** Increase in the distance (indicated be yellow arrows) between SR (white arrow) and the nearest sarcolemmal (Mem, black arrow) membranes in HF SANCs (n=77 cells from 4 mice). Cav – caveola, M – mitochondria. Control data is reproduced from Figure 4A for comparison. **(E)** *In vitro* confocal imaging of spontaneous [Ca^2+^]*_i_* in control and HF SANCs. Control CaT traces are reproduced from Figure 3A1. CaT CL periods are shown in blue, LCR (indicated by red arrows on confocal scans) CL periods are shown in red. Green arrows indicate LCRs that failed to trigger APs. LCR-to-CaT dissociation is indicated by beat-to-beat LCR and CaT CL Poincaré scatter plots. Control LCR amplitude data is reproduced from Figure 3C. Control LCR latency data is reproduced from Figure 3D. *P*-values by Student’s t-test.

Importantly, HF remodeling resulted in significant downregulation of Cav-3 protein expression and caveolae in SAN tissues (Fig. 8C). Similar to Cav-3KO*-*SANCs (Fig. 4A), this was associated with a significantly increased distance between sarcolemmal and SR membranes, from 18 ± 1 nm in AMC (N/n = 6/76 mice/cells) to 34 ± 2 nm in HF (*P*<0.001, Fig. 8D), and impaired spatial-temporal coupling between Ca^2+^ (LCRs) and V_m_ (AP-induced CaTs) clocks leading to highly irregular AP-induced CaTs and prolonged SAN pauses (Fig. 8E). Overall, the observed findings indicate that HF-induced disruption of caveolae structures is likely associated with similar mechanisms as those observed in Cav-3KO mice as well as MβCD-WT SANCs, and reveal caveolae and caveolar macromolecular pacemaker complex as new contributors to SAN remodeling and SND.

## Discussion

Despite extensive research into the cellular and molecular mechanisms of SAN pacemaking over the past several decades and increasing understanding of the contribution of individual components of the coupled-clock system to spontaneous AP firing, significant gaps remain in our knowledge of the structural organization of the SAN pacemaker system and the spatial-temporal interactions among its components. This gap is particularly crucial to address because simply altering the function of one or more pacemaker ion currents cannot fully explain the complex SND phenotype observed in both Cav-3-KO and HF mice, characterized by bradycardia, sinus pauses, recurrent development of SAN quiescence, and profound beat-to-beat CL variations. Our findings significantly contribute to the emerging evidence highlighting an additional layer of pacemaker organization,^4,6,29,32^ i.e., a tightly coupled, interacting network of molecules involved in pacemaker activity. In the present study, we demonstrated, in both mouse and human SANCs, the pivotal role of caveolae in organizing sarcolemmal electrogenic proteins within a specialized pacemaker signalosome, mediated by their association with Cav-3 scaffolding protein. This organization ensures the close proximity of these proteins to the subsarcolemmal SR membrane, facilitating the fast and robust activation of NCX1 by RyR2-mediated LCRs during diastolic depolarization that enables AP triggering and supports a stable, regular SAN rhythm.

### Disruption of caveolar pacemaker signalosome and SND

Our study revealed caveolae downregulation in Cav3-KO and HF mice, which disrupted the physical arrangement between the SR and sarcolemmal membranes. This led to a redistribution of NCX1 to non-caveolar areas, limiting its ability to be effectively activated by LCRs and to initiate SANC APs, and resulted in LCR-to-AP failure, sinus pauses, increased heart rate lability, tachycardia-bradycardia paroxysms, and enhanced atrial ectopy. Interestingly, increased ectopic activity coincided with periods of tachycardia, whereas the localization of the leading pacemaker within the SAN region was observed during bradycardic rhythm, in both Cav-3KO (Fig. 2E) and HF mice (Fig. 8A,B). Similar observations were recently reported by Landstrom et al. in tamoxifen-inducible, pacemaker tissue-specific (Hcn4-targeted), junctophilin 2 knockdown mice (Hcn4:shJph2).^33^ Junctophilin 2 provides a structural bridge between the sarcolemma and SR membranes in atrial and ventricular working myocytes, facilitating the functional coupling between LTCCs and RyR2 at dyadic junctions.^34^ However, it remains unknown whether junctophilin 2 serves a similar function in pacemaker cells. Notably, junctophilin 2 is closely associated with Cav-3 in ventricular myocytes,^35^ suggesting a potential contribution to caveolae-SR coupling in SANCs. This is consistent with early termination of spontaneous membrane depolarizations observed in Hcn4:shJph2 SANCs that failed to trigger AP,^34^ akin to our observations in Cav-3KO (Fig. 3A) and HF mice (Fig. 6E) during caveolae-mediated LCR-to-AP uncoupling. These findings further highlight the key role of structural clock coupling in the mechanism of SAN pacemaking.

As demonstrated in Cav-3KO mice (Fig. 6D) and observed in Hcn4:shJph2 mice,^34^ SND resulting from V_m_-Ca^2+^-clock uncoupling is associated with a substantial (∼50%) reduction in NCX1 function, despite unaltered NCX1 expression. Using our novel, nanodomain-specific SAN model, we demonstrated that the SND phenotype could be sourced from redistribution of 60% caveolar NCX1 to the extra-caveolar membrane (Fig. 5B), consistent with findings from proximity ligation assay experiments (Fig. 4D), leading to the inability of LCRs to effectively increase NCX current and thus hindering the generation of APs. This mechanism is further supported by a significant prolongation of LCR latency observed in Cav-3KO (Fig. 3D) and HF mice (Fig. 6E). Extreme scenarios of complete either pharmacological (Fig. S16) or genetic^29^ NCX1 inhibition or redistribution of 80% of caveolar NCX1 to extra-caveolar membrane (Fig. 5B, *model 6*) would lead to LCR-to-AP failure and loss of SAN excitability.

Our results also demonstrate a significant (∼50%) decrease in *I*_Ca,L_ density (Fig. 3C) and a leftward shift of *I*_f_ activation curve (Fig. 3B) in Cav-3KO mice which could further contribute to V_m_-Ca^2+^-clock uncoupling by decreasing the overall depolarizing force (both during diastolic depolarization and upstroke phase) in SANCs in the setting of significantly weakened NCX1 function. Similar *I*_Ca,L_ decrease has been previously shown in rat atrial myocytes during caveolae disruption by MβCD (by 30%),^16^ in Cav-3KO mouse ventricular myocytes (by 28%),^12^ and in rat ventricular myocytes (by 30%) during Cav-3 inhibition by a membrane-permeable TAT-tagged peptide C3SD^36^ (designed to disrupt binding of Cav-3 to its partner proteins). The observed decrease in *I*_Ca,L_ density could be associated with a reduction in single LTCC availability on the membrane^16^ and/or decrease in single channel open probability (P_o_) due to channel dephosphorylation,^36^ probably via disruption of Cav-3-mediated LTCC coupling with protein kinase A (PKA).^37^ The latter mechanism could be associated with the impairment of Cav-3 mediated cAMP signaling, as recently demonstrated in SAN,^4^ atrial,^17^ and ventricular myocytes^24,38^ during caveolae disruption, which could also explain the leftward shift of the *I*_f_ activation curve (Fig. 3B). At the same time, we did not observe caveolae-mediated changes in *I*_Ca,T_ current (Fig. 3C), which could be augmented via both PKA and protein kinase C (PKC) phosphorylation.^39^ In contrast, *I*_Ca,L_ is suppressed by PKC activation,^40^ and PKC activation has been linked to caveolae degradation.^41^ Therefore, a decrease in PKA activity, an increase in PKC activity, or their combination could contribute to the observed changes during caveolae disruption in Cav-3-KO mice. Importantly, our combined experimental/simulation studies indicate that neither inhibition of *I*_f_, *I*_Ca,L_ or *I*_Ca,T_ (Fig. 7), nor their redistribution from caveolae nanodomains (Figs. S14 and S15) could recapitulate the observed SND phenotype.

### Enhanced atrial ectopy

In both Cav-3KO (Fig. 3) and HF mice (Fig. 6), SND was accompanied by dramatically elevated atrial ectopic activity, which was also previously observed in various SND models^34,42,43^ and linked to hyperactive subsidiary atrial pacemakers. Characterized by both HCN4- and Cx43-positive expression,^26,42^ subsidiary atrial pacemakers are widely distributed throughout the entire region located between the superior and inferior vena cava and between the crista terminalis and intra-atrial septum.^25^ Though the subsidiary pacemakers can provide a relatively regular rhythm, they are characterized by slower resting and exertional rates, prolonged post-pacing recovery times, and increased beat-to-beat heart rate variability.^26,44^ The mechanism of subsidiary pacemaker automaticity is believed to rely on the expression of *I*_Na_^45^ and a greater contribution of LCR-mediated *I*_NCX_ to spontaneous diastolic depolarization than *I*_f_.^46^ Indeed, the crucial role of the Ca^2+^ clock in spontaneous automaticity has been highlighted in the subsidiary pacemakers, especially under sympathetic stimulation.^34,42^ Along with this mechanism, our findings indicate that Cav-3KO*-*SANCs have abnormal Ca^2+^ signaling characterized by elevated LCR amplitude (Fig. 3C) and augmentation of diastolic Ca^2+^ spark activity (Fig. S12). In a recent study, we demonstrated that caveolae disruption in mouse atrial myocytes by acute mechanical stretch, cholesterol depletion by MβCD, or *Cav-3* knock-out results in the elevation of cAMP level and cAMP-mediated augmentation of PKA-phosphorylation of RyR2 at Ser^2030^ and Ser^2808^ sites.^17^ However, enhanced atrial ectopy observed in Cav-3KO mice was not associated with sustained atrial flutter/fibrillation pattern, probably due to a lack of the structural atrial remodeling required for sustained atrial tachyarrhythmias.^42,47^

### Conclusions

Our results provide the first direct experimental evidence linking caveolae to SND and atrial arrhythmogenesis. These findings hold profound translational significance, as patients with *CAV3* variants exhibit sinus bradycardia, elevated heart rhythm lability, and cardiac arrest during sleep.^21^ Additionally, caveolae disruption and Cav-3 downregulation or malfunction are commonly observed in various pathological conditions, including HF,^18^ atrial fibrillation,^19^ and hypertension,^20^ all of which are associated with SND. Recent studies have also identified an age-dependent decrease in Cav-3 protein expression and caveolae density in both aged mouse and human atrial tissues,^48^ suggesting a potential role in the decline of SAN function and the development of SND in the elderly.

## Funding sources

This work was supported by NIH 2R01HL141214 (A.V.G., E.G., and T.J.K.), 1R01HL170521 (A.V.G. and E.G.), R01HL139738, R01HL146652 (A.V.G.), R01HL131517, P01 HL141084 (E.G.), R00HL138160 (S.M.), R01HL078878 (T.J.K.), R35NS116850 (B.C.) and 1R01HL131403-01A1 (G.A.R.); NIH Stimulating Peripheral Activity to Relieve Conditions grant 1OT2OD026580-01 (E.G.); AHA 17POST33370089, 846898, and 24SCEFIA1255230 (D.L.), AHA 20POST35120462 (H.N.), AHA 915335 (R.Y.M.), and AHA 903203 (D.G.P.T.); UC Davis School of Medicine Dean’s Fellow award (E.G.). D.T. would like to acknowledge NIH predoctoral training grant T32GM008688.

## Notes

### Competing Interest Statement

The authors have declared no competing interest.

### Summary of Updates

The revised version of the preprint contains an updated supplemental file, including corrected and new supplemental figures and methodological details, updated statistical analysis, revised main Figures 1, 4, and 8, as well as updated authorship list.

## References

1. Benditt D, Sakaguchi S, Goldstein M, Lurie K, Gornick C. Sinus node dysfunction: Pathophysiology, clinical features, evaluation, and treatment. *In:* Zipes DP, Jalife J, *eds* Cardiac Electrophysiology: From Cell to Bedside Philadelphia:WBSaunders Company. 1995:1215–1247.

2. Lakatta EG, Maltsev VA, Vinogradova TM. A coupled SYSTEM of intracellular Ca2+ clocks and surface membrane voltage clocks controls the timekeeping mechanism of the heart’s pacemaker. Circ Res. 2010;106:659–673. doi: 10.1161/CIRCRESAHA.109.206078

3. Lakatta EG, DiFrancesco D. What keeps us ticking: a funny current, a calcium clock, or both? J Mol Cell Cardiol. 2009;47:157–170. doi: 10.1016/j.yjmcc.2009.03.022

4. Ren L, Thai PN, Gopireddy RR, Timofeyev V, Ledford HA, Woltz RL, Park S, Puglisi JL, Moreno CM, Santana LF, et al. Adenylyl cyclase isoform 1 contributes to sinoatrial node automaticity via functional microdomains. JCI Insight. 2022;7. doi: 10.1172/jci.insight.162602

5. Lyashkov AE, Juhaszova M, Dobrzynski H, Vinogradova TM, Maltsev VA, Juhasz O, Spurgeon HA, Sollott SJ, Lakatta EG. Calcium cycling protein density and functional importance to automaticity of isolated sinoatrial nodal cells are independent of cell size. Circ Res. 2007;100:1723–1731. doi: 10.1161/CIRCRESAHA.107.153676

6. Lang D, Glukhov AV. Functional Microdomains in Heart’s Pacemaker: A Step Beyond Classical Electrophysiology and Remodeling. Front Physiol. 2018;9:1686. doi: 10.3389/fphys.2018.01686

7. Ayettey AS, Navaratnam V. The T-tubule system in the specialized and general myocardium of the rat. J Anat. 1978;127:125–140.

8. Masson-Pevet M, Gros D, Besselsen E. The caveolae in rabbit sinus node and atrium. Cell and tissue research. 1980;208:183–196.

9. Song KS, Scherer PE, Tang Z, Okamoto T, Li S, Chafel M, Chu C, Kohtz DS, Lisanti MP. Expression of caveolin-3 in skeletal, cardiac, and smooth muscle cells. Caveolin-3 is a component of the sarcolemma and co-fractionates with dystrophin and dystrophin-associated glycoproteins. J Biol Chem. 1996;271:15160–15165.

10. Balijepalli RC, Foell JD, Hall DD, Hell JW, Kamp TJ. Localization of cardiac L-type Ca(2+) channels to a caveolar macromolecular signaling complex is required for beta(2)-adrenergic regulation. Proc Natl Acad Sci U S A. 2006;103:7500–7505. doi: 10.1073/pnas.0503465103

11. Balijepalli RC, Kamp TJ. Caveolae, ion channels and cardiac arrhythmias. Prog Biophys Mol Biol. 2008;98:149–160. doi: 10.1016/j.pbiomolbio.2009.01.012

12. Markandeya YS, Gregorich ZR, Feng L, Ramchandran V, O’ Hara T, Vaidyanathan R, Mansfield C, Keefe AM, Beglinger CJ, Best JM, et al. Caveolin-3 and Caveolae regulate ventricular repolarization. J Mol Cell Cardiol. 2023;177:38–49. doi: 10.1016/j.yjmcc.2023.02.005

13. Tyan L, Foell JD, Vincent KP, Woon MT, Mesquitta WT, Lang D, Best JM, Ackerman MJ, McCulloch AD, Glukhov AV, et al. Long QT syndrome caveolin-3 mutations differentially modulate K v 4 and Ca v 1.2 channels to contribute to action potential prolongation. J Physiol. 2019;597:1531–1551. doi: 10.1113/JP276014

14. Tyan L, Turner D, Komp KR, Medvedev RY, Lim E, Glukhov AV. Caveolin-3 is required for regulation of transient outward potassium current by angiotensin II in mouse atrial myocytes. Am J Physiol Heart Circ Physiol. 2021;320:H787–H797. doi: 10.1152/ajpheart.00569.2020

15. Zaza A, Grandi E. Mechanisms of Cav3-associated arrhythmia: Protein or microdomain dysfunction? Int J Cardiol. 2020;320:97–99. doi: 10.1016/j.ijcard.2020.06.051

16. Glukhov AV, Balycheva M, Sanchez-Alonso JL, Ilkan Z, Alvarez-Laviada A, Bhogal N, Diakonov I, Schobesberger S, Sikkel MB, Bhargava A, et al. Direct Evidence for Microdomain-Specific Localization and Remodeling of Functional L-Type Calcium Channels in Rat and Human Atrial Myocytes. Circulation. 2015;132:2372–2384. doi: 10.1161/CIRCULATIONAHA.115.018131

17. Medvedev RY, Turner DGP, DeGuire FC, Leonov V, Lang D, Gorelik J, Alvarado FJ, Bondarenko VE, Glukhov AV. Caveolae-associated cAMP/Ca2^+^-mediated mechano-chemical signal transduction in mouse atrial myocytes. J Mol Cell Cardiol. 2023;184:75–87. doi: 10.1016/j.yjmcc.2023.10.004

18. Ratajczak P, Damy T, Heymes C, Oliviero P, Marotte F, Robidel E, Sercombe R, Boczkowski J, Rappaport L, Samuel JL. Caveolin-1 and -3 dissociations from caveolae to cytosol in the heart during aging and after myocardial infarction in rat. Cardiovasc Res. 2003;57:358–369.

19. Reilly SN, Liu X, Carnicer R, Recalde A, Muszkiewicz A, Jayaram R, Carena MC, Wijesurendra R, Stefanini M, Surdo NC, et al. Up-regulation of miR-31 in human atrial fibrillation begets the arrhythmia by depleting dystrophin and neuronal nitric oxide synthase. Science translational medicine. 2016;8:340ra374. doi: 10.1126/scitranslmed.aac4296

20. Egorov YV, Lang D, Tyan L, Turner D, Lim E, Piro ZD, Hernandez JJ, Lodin R, Wang R, Schmuck EG, et al. Caveolae-Mediated Activation of Mechanosensitive Chloride Channels in Pulmonary Veins Triggers Atrial Arrhythmogenesis. Journal of the American Heart Association. 2019;8:e012748. doi: 10.1161/JAHA.119.012748

21. Vatta M, Ackerman MJ, Ye B, Makielski JC, Ughanze EE, Taylor EW, Tester DJ, Balijepalli RC, Foell JD, Li Z, et al. Mutant caveolin-3 induces persistent late sodium current and is associated with long-QT syndrome. Circulation. 2006;114:2104–2112. doi: 10.1161/CIRCULATIONAHA.106.635268

22. Opthof T, de Jonge B, Jongsma HJ, Bouman LN. Functional morphology of the pig sinoatrial node. J Mol Cell Cardiol. 1987;19:1221–1236.

23. Couet J, Li S, Okamoto T, Ikezu T, Lisanti MP. Identification of peptide and protein ligands for the caveolin-scaffolding domain. Implications for the interaction of caveolin with caveolae-associated proteins. J Biol Chem. 1997;272:6525–6533. doi: 10.1074/jbc.272.10.6525

24. Wright PT, Diakonov I, Pannell L, Perera RK, Bork NI, Alvarez-Laviada A, Schobesberger S, Lucarelli C, Faggian G, Bhogal NK, et al. Cardiomyocyte membrane structure and cAMP compartmentation produce anatomical variation in β2AR-cAMP responsiveness in murine hearts. Cell Reports. 2018;23:459–469.

25. Boineau JP, Canavan TE, Schuessler RB, Cain ME, Corr PB, Cox JL. Demonstration of a widely distributed atrial pacemaker complex in the human heart. Circulation. 1988;77:1221–1237.

26. Morris GM, D’Souza A, Dobrzynski H, Lei M, Choudhury M, Billeter R, Kryukova Y, Robinson RB, Kingston PA, Boyett MR. Characterization of a right atrial subsidiary pacemaker and acceleration of the pacing rate by HCN over-expression. Cardiovasc Res. 2013;100:160–169. doi: 10.1093/cvr/cvt164

27. Le Scouarnec S, Bhasin N, Vieyres C, Hund TJ, Cunha SR, Koval O, Marionneau C, Chen B, Wu Y, Demolombe S, et al. Dysfunction in ankyrin-B-dependent ion channel and transporter targeting causes human sinus node disease. Proc Natl Acad Sci U S A. 2008;105:15617–15622. doi: 0805500105 [pii] 10.1073/pnas.0805500105

28. Musa H, Lei M, Honjo H, Jones SA, Dobrzynski H, Lancaster MK, Takagishi Y, Henderson Z, Kodama I, Boyett MR. Heterogeneous expression of Ca(2+) handling proteins in rabbit sinoatrial node. J Histochem Cytochem. 2002;50:311–324. doi: 10.1177/002215540205000303

29. Torrente AG, Zhang R, Zaini A, Giani JF, Kang J, Lamp ST, Philipson KD, Goldhaber JI. Burst pacemaker activity of the sinoatrial node in sodium-calcium exchanger knockout mice. Proc Natl Acad Sci U S A. 2015;112:9769–9774. doi: 10.1073/pnas.1505670112

30. Sanders P, Kistler PM, Morton JB, Spence SJ, Kalman JM. Remodeling of sinus node function in patients with congestive heart failure: reduction in sinus node reserve. Circulation. 2004;110:897–903. doi: 10.1161/01.CIR.0000139336.69955.AB

31. Luu M, Stevenson WG, Stevenson LW, Baron K, Walden J. Diverse mechanisms of unexpected cardiac arrest in advanced heart failure. Circulation. 1989;80:1675–1680.

32. Maltsev AV, Yaniv Y, Stern MD, Lakatta EG, Maltsev VA. RyR-NCX-SERCA local cross-talk ensures pacemaker cell function at rest and during the fight-or-flight reflex. Circ Res. 2013;113:e94–e100. doi: 10.1161/CIRCRESAHA.113.302465

33. Landstrom AP, Yang Q, Sun B, Perelli RM, Bidzimou MT, Zhang Z, Aguilar-Sanchez Y, Alsina KM, Cao S, Reynolds JO, et al. Reduction in Junctophilin 2 Expression in Cardiac Nodal Tissue Results in Intracellular Calcium-Driven Increase in Nodal Cell Automaticity. Circ Arrhythm Electrophysiol. 2023;16:e010858. doi: 10.1161/CIRCEP.122.010858

34. van Oort RJ, Garbino A, Wang W, Dixit SS, Landstrom AP, Gaur N, De Almeida AC, Skapura DG, Rudy Y, Burns AR, et al. Disrupted junctional membrane complexes and hyperactive ryanodine receptors after acute junctophilin knockdown in mice. Circulation. 2011;123:979–988. doi: 10.1161/CIRCULATIONAHA.110.006437

35. Poulet C, Sanchez-Alonso J, Swiatlowska P, Mouy F, Lucarelli C, Alvarez-Laviada A, Gross P, Terracciano C, Houser S, Gorelik J. Junctophilin-2 tethers T-tubules and recruits functional L-type calcium channels to lipid rafts in adult cardiomyocytes. Cardiovasc Res. 2021;117:149–161. doi: 10.1093/cvr/cvaa033

36. Bryant S, Kimura TE, Kong CH, Watson JJ, Chase A, Suleiman MS, James AF, Orchard CH. Stimulation of ICa by basal PKA activity is facilitated by caveolin-3 in cardiac ventricular myocytes. J Mol Cell Cardiol. 2014;68:47–55. doi: 10.1016/j.yjmcc.2013.12.026

37. Macdougall DA, Agarwal SR, Stopford EA, Chu H, Collins JA, Longster AL, Colyer J, Harvey RD, Calaghan S. Caveolae compartmentalise beta2-adrenoceptor signals by curtailing cAMP production and maintaining phosphatase activity in the sarcoplasmic reticulum of the adult ventricular myocyte. J Mol Cell Cardiol. 2012;52:388–400. doi: 10.1016/j.yjmcc.2011.06.014

38. Wright PT, Nikolaev VO, O’Hara T, Diakonov I, Bhargava A, Tokar S, Schobesberger S, Shevchuk AI, Sikkel MB, Wilkinson R, et al. Caveolin-3 regulates compartmentation of cardiomyocyte beta2-adrenergic receptor-mediated cAMP signaling. J Mol Cell Cardiol. 2014;67:38–48. doi: 10.1016/j.yjmcc.2013.12.003

39. Chemin J, Mezghrani A, Bidaud I, Dupasquier S, Marger F, Barrère C, Nargeot J, Lory P. Temperature-dependent modulation of CaV3 T-type calcium channels by protein kinases C and A in mammalian cells. J Biol Chem. 2007;282:32710–32718. doi: 10.1074/jbc.M702746200

40. Luo AT, Luo HY, Hu XW, Gao LL, Liang HM, Tang M, Hescheler J. Hyposmotic challenge modulates function of L-type calcium channel in rat ventricular myocytes through protein kinase C. Acta Pharmacol Sin. 2010;31:1438–1446. doi: 10.1038/aps.2010.112

41. Markandeya YS, Phelan LJ, Woon MT, Keefe AM, Reynolds CR, August BK, Hacker TA, Roth DM, Patel HH, Balijepalli RC. Caveolin-3 Overexpression Attenuates Cardiac Hypertrophy via Inhibition of T-type Ca2+ Current Modulated by Protein Kinase Calpha in Cardiomyocytes. J Biol Chem. 2015;290:22085–22100. doi: 10.1074/jbc.M115.674945

42. Glukhov AV, Kalyanasundaram A, Lou Q, Hage LT, Hansen BJ, Belevych AE, Mohler PJ, Knollmann BC, Periasamy M, Gyorke S, et al. Calsequestrin 2 deletion causes sinoatrial node dysfunction and atrial arrhythmias associated with altered sarcoplasmic reticulum calcium cycling and degenerative fibrosis within the mouse atrial pacemaker complex1. Eur Heart J. 2015;36:686–697. doi: 10.1093/eurheartj/eht452

43. Glukhov AV, Fedorov VV, Anderson ME, Mohler PJ, Efimov IR. Functional anatomy of the murine sinus node: high-resolution optical mapping of ankyrin-B heterozygous mice. Am J Physiol Heart Circ Physiol. 2010;299:H482–491. doi: ajpheart.00756.2009 [pii] 10.1152/ajpheart.00756.2009

44. Choudhury M, Black N, Alghamdi A, D’Souza A, Wang R, Yanni J, Dobrzynski H, Kingston PA, Zhang H, Boyett MR, et al. TBX18 overexpression enhances pacemaker function in a rat subsidiary atrial pacemaker model of sick sinus syndrome. J Physiol. 2018;596:6141–6155. doi: 10.1113/JP276508

45. Lei M, Goddard C, Liu J, Leoni AL, Royer A, Fung SS, Xiao G, Ma A, Zhang H, Charpentier F, et al. Sinus node dysfunction following targeted disruption of the murine cardiac sodium channel gene Scn5a. J Physiol. 2005;567:387–400. doi: jphysiol.2005.083188 [pii] 10.1113/jphysiol.2005.083188

46. Lang D, Glukhov AV. Cellular and Molecular Mechanisms of Functional Hierarchy of Pacemaker Clusters in the Sinoatrial Node: New Insights into Sick Sinus Syndrome. J Cardiovasc Dev Dis. 2021;8. doi: 10.3390/jcdd8040043

47. Zhang Y, Qi Y, Li JJ, He WJ, Gao XH, Zhang Y, Sun X, Tong J, Zhang J, Deng XL, et al. Stretch-induced sarcoplasmic reticulum calcium leak is causatively associated with atrial fibrillation in pressure-overloaded hearts. Cardiovasc Res. 2020. doi: 10.1093/cvr/cvaa163

48. Peart JN, Pepe S, Reichelt ME, Beckett N, See Hoe L, Ozberk V, Niesman IR, Patel HH, Headrick JP. Dysfunctional survival-signaling and stress-intolerance in aged murine and human myocardium. Exp Gerontol. 2014;50:72–81. doi: 10.1016/j.exger.2013.11.015

