## Supplementary Materials for "Caveolar Compartmentalization of Pacemaker Signaling is Required for Stable Rhythmicity of Sinus Nodal Cells and is Disrupted in Heart Failure"

### Methods

#### *Ethical approval*

All experiments were conducted in accordance with the National Institutes of Health Guide for the Care and Use of Laboratory Animals (NIH Pub. No. 80-23). All methods and protocols used in these studies have been approved by the Animal Care and Use Committee of University of Wisconsin-Madison (U.S.A.) following the Guidelines for Care and Use of Laboratory Animals published by NIH (publication No. 85-23, revised 1996). All animals used in this study received humane care in compliance with the Guide for the Care and Use of Laboratory Animals.

#### *Human tissue collection*

Human heart collection protocols were approved by the University of Wisconsin Institutional Review Board. Non-failing human hearts that went unused for organ transplant were obtained from the University of Wisconsin Organ Procurement Organization, Madison, WI. At the time of harvest, hearts were aseptically excised, perfused and stored in cold cardioplegia solution, and transported on ice as described previously in details (1-3). Human sinoatrial node (SAN) was identified as a condensed area of tissue located at the superior cavoatrial junction, spread adjacent to the crista terminalis about 2 cm towards inferior vena cava and usually being arranged around a prominent nodal as it was demonstrated previously both structurally and functionally (4-8). The location of the SAN was confirmed by a positive expression of HCN4 channels (responsible for pacemaker $I_f$ current) and a negative expression of atrial-specific connexin43 protein (**Data Supplement Figure S4**). Human SAN tissue was used for co-immunoprecipitation experiments (**Fig. 1E and Supplementary Figures S5 and S6**).

#### *Animals*

Adult (5-6-month-old) male and female cardiac-specific caveolin-3 knock-out mice (Cav-3KO) and wild type littermate controls (WT; flox+/Cre-) as well as 8-weeks post myocardial infarction heart failure male mice (HF) and age-matched control male mice (AMC) were used in the study. Because of significant sex differences in HF development (9-11), in the present study, we specifically used only male mice for HF model, including AMC male mice as control. In addition, because we did not observe any significant difference between Cav-3-WT littermate control mice and AMC mice, both *in vivo* and *ex vivo* (Figs. 2 and 8), for biochemical and single cell electrophysiological experiments AMC wild type mice were used as controls for both Cav-3KO and HF groups.

Cav-3 conditional knock-out mice were generated by crossing a mouse line with loxP-flanked exon 2 of Cav-3 with an α-myosin heavy chain proto-oncogene tyrosine-protein kinase MER Cre recombinase proto-oncogene tyro- sine-protein kinase MER (aMHC-MERCreMER) (12) mouse producing a tamoxifen-inducible, cardiac-specific Cav-3KO mouse (13-16). Mice were given an intraperitoneal injection of 1 mg tamoxifen/day for 3 consecutive days and sacrificed 21 days post-tamoxifen treatment.

Heart failure mouse model was done in the cardiovascular physiology core facility at the University of Wisconsin Madison. HF was induced by 8-weeks of myocardial infarction (MI) in C57BL/6J male mice by ligating the left anterior descending coronary artery as previously described (17). Specifically, mice were anesthetized and ventilated with 2% isoflurane and oxygen. Followed by a left thoracotomy, the left anterior descending coronary artery (LAD) was ligated with 8-0 nylon suture. Successful acute myocardial ischemia was confirmed when left ventricular wall became pale. Mice then were kept on heating pad until awake.

HF progression was verified by echocardiographic analysis at 8 weeks post-MI. Parameters including ejection fraction (EF) and left ventricular (LV) mass are shown in **Data Supplement Figure S17**. Specifically, systolic and diastolic function was assessed by echocardiographic examination in lightly anesthetized (1.5% isoflurane) mice using a 17.5-MHz transducer (RMV 707B; Visual Sonics, Toronto, ON, Canada), as previously described (18). Two-dimensional M-Mode images were recorded both in the short and long axes. Average values were obtained from the measurement of 3-5 consecutive cardiac cycles.

#### *Caveolin-binding motif (CBM) analysis*

Proteins' CBMs were identified based on the following motifs: ΦXXXXΦXXΦ or ΦXΦXXXXΦ where Φ is an aromatic amino acid and X is a nonaromatic amino acid (**Data Supplement Figure S8**). In brief, Python was used to convert protein amino acid sequences into binary and scanned for the aforementioned motifs. The python code is:

```
# first string
firstString = "FWHYARNDCQEGILKMPSTV"
secondString = "11112222222222222222"
string = "input amino acid sequence here"
print("Original string:", string)
translation = string.maketrans(firstString, secondString)
# translate string
print("Translated string:", string.translate(translation))
```

#### *In vivo ECG Recording*

Surface ECG recordings were done as previously described (19). Briefly, mice were lightly anesthetized with isoflurane vapor titrated to maintain the lightest anesthesia possible. On average, 1.5% volume isoflurane vapor was required to maintain adequate anesthesia. Loss of toe pinch reflex and respiration rate was used to monitor levels of anesthesia. Average respiration rate was not different between mouse groups. ECG was recorded for 5 minutes, and heart rate was measured as the average over a 30-second interval.

***Optical mapping***

The isolated heart preparations were performed as previously described (19-21). Briefly, mice were anesthetized with isoflurane (induced at 3-5% and maintained at 1-3%) to assure the appropriate level of anesthesia by checking the loss of pain reflex. After midsternal incision, the heart was removed and placed in oxygenated (95% $O_2$, 5% $CO_2$), mid-hypothermia (33 ± 1°C), modified Tyrode solution of the following composition: 128.2 mM NaCl, 4.7 mM KCl, 1.19 $NaH_2PO_4$, 1.05 mM $MgCl_2$, 1.3 mM $CaCl_2$, 20.0 mM $NaHCO_3$, and 11.1 mM glucose (pH = 7.35 ± 0.05). While bathed in the same solution, lung, thymus, and fat tissue were removed. A short section of aorta was attached to a custom made 21-gauge cannula. After cannulation, the heart was superfused and retrogradely perfused with Tyrode solution passed through the 5-µm filter (Millipore, Billerica) and warmed (37°C) using a water jacket and circulator (ThermoNESLAB EX7, Newtown). Perfusion was performed using a peristaltic pump (Peri-Star, WPI, Sarasota) under constant aortic pressure of 60 – 80 mmHg measured by pressure-amplifier (TBM4M, WPI, Sarasota).

For isolated SAN preparations, after cannulation ventricles were dissected away, and the atria were flattened and then pinned to the bottom of a Sylgard-coated chamber and superfused with Tyrode solution at a constant rate of ~15 ml/min. The medial limb of the crista terminalis (CT) was cut in order to open the right atria appendage (RAA), and the pacing electrode was placed on the edge of the RAA. Two Ag/AgCl electrodes were immersed into the superfusion solution and placed on the sides of preparation to document pseudo-ECG. The inter-atrial septum (IAS) tissue was partially removed to reduce scattering of the optical signal from tissue that was not in focus. A rim of ventricular tissue was preserved for pinning the preparation in order to prevent damage to the atria. Both the left and right atria (LA and RA) as well as the atrio-ventricular junction (AVJ) were accessible in the preparation.

***Imaging system***

After isolation, the SAN preparations were then immobilized by infusion of Blebbistatin (10 µM, Tocris Bioscience, USA) in the perfusion media in order to suppress motion artifacts in optical recordings. Then, the preparations were co-stained with voltage-sensitive dye RH-237 (ThermoFisher Scientific, USA) (1.25 mg/ml in dimethyl sulfoxide) by direct application of the dye on the tissue. Excitation light (520/44 nm) was generated by a 150-W halogen lamp with a constant-current, low-noise, power supply (MHAB-150W, Moritex USA Inc., CA, USA). A flexible light guide directed the band-pass-filtered light onto the preparation and a shutter was used to ensure that the preparation was exposed to light only during image acquisition. The fluorescent light emitted from the preparation was long-pass (>715 nm) filtered using an edge pass filter (Thorlabs, NJ) before reaching the camera. Emitted light was directed towards a MiCAM Ultima-L CMOS camera (SciMedia, CA, USA) with high spatial (100 x 100 pixels, 60 ± 10 µm per pixel) and temporal (1,000 - 3,000 frames/sec) resolution. The acquired fluorescent signal was digitized, amplified, and visualized using custom software (SciMedia, CA, USA).

***Experimental protocol***

After isolation, motion suppression and staining, preparations were equilibrated for 20 min before imaging. Next, control maps of atrial activation during spontaneous rhythm were made as previously described (21). Location of the leading pacemaker was visualized and attributed to one of the four anatomical regions as previously described based on the location and the presence of HCN4 and connexin43 protein expression (19): (1) primary SAN (as characterized by HCN4-positive and Cx43-negative expression pattern), (2) lower SAN (as characterized by both HCN4- and Cx43-positive expression pattern and located along the CR between the SVC and IVC), (3) intra-atrial septum (IAS), and (4) latent pacemakers (as characterized by both HCN4- and Cx43-positive expression pattern and located outside of SAN region and ICR). One mouse could have several sites of the leading pacemaker location because of the presence of multiple competing pacemakers during the optical recording.

Heart rhythm stability was calculated as shown in **Figure 2** – average CL and CL lability. Average CL was calculated for 150 continuous beats per mouse at baseline condition after a 30-min of stabilization period. CL lability is calculated as the standard deviation of 150 continuous beats.

To estimate SAN recovery time (SANRT), the preparation was paced (S1S1 = 100 ms) for at least 30 secs through a pacing electrode located at the RA appendage (22). SANRT was measured as the time interval between the last pacing beat and the first spontaneous beat. Corrected SANRT (SANRTc) was calculated as the difference between the SANRT and the resting cycle length (CL) measured before the SANRT pacing protocol.

***Optical mapping data processing***

A customized Matlab-based computer program was used to analyze the optical signals (20,21,23). Signals were filtered using the low-pass Butterworth algorithm at 256 Hz. Maximum upstroke derivative ($dV/dt_{max}$) was calculated for each action potential using the normalized optical signal and its derivatives. Activation maps were constructed from activation times, which were determined from the $dV/dt_{max}$.

***SAN Cell Isolation***

Single SAN cells were isolated from mice using a modified method based on previously published protocols (19,24-26). Cells exclusively from the region which corresponds to the primary pacemaker area characterized previously in the mouse heart by histology and immunolabeling of connexin43, connexin45, and HCN (19,27,28) as well as functionally by microelectrodes (29,30) and optical mapping (19-21,31). Briefly, after heart isolation and cannulation, the isolated SAN preparations were prepared as described above. Subsequently, the SAN region, bordered by the crista terminalis, atrial septum, and orifice of superior vena cava, was dissected from the heart. The SAN area was cut into strips which then was washed three times with 2 mins each time in the 'low $Ca^{2+}$, $Mg^{2+}$ free' solution (in mmol/L: 140 NaCl, 5.4 KCL, 1.2 $KH_2PO_4$, 0.066 $CaCl_2$, 50 taurine, 18.5 D-glucose, 5 HEPES and 1mg/ml bovine serum albumin (BSA), with pH adjusted to 6.9 with NaOH). The SAN tissue pieces were digested for 30 mins at 35°C in a cocktail of 1mg/ml Collagenase (Sigma) and 0.2mg/ml Elastase (Worthington) with gentle shaking every 5 min. The digested tissue was carefully rinsed with Kraft-Brühe solution (in mmol/L: 100 potassium glutamate, 10 potassium aspartate, 25 KCL, 10 $KH_2PO_4$, 2 $MgSO_4$, 20 taurine, 5 creatine, 0.5 EGTA, 20 glucose, 5 HEPES, and 0.1% BSA, with pH adjusted to 7.4 with KOH) and gently triturated with a pair of fire polished, wide-bore Pasteur pipettes to release SAN cells. Isolated cells were then plated on Laminin (Sigma) coated coverslips for 30 mins, and readapted to normal extracellular $Ca^{2+}$ concentration of 1.8 mmol/L.

SAN myocytes were identified by their small spindle shape and ability to beat spontaneously in the recording chamber when superfused with normal Tyrode's solution. We confirmed pacemaker phenotype in isolated cells by HCN4-positive and connexin43-negative immunofluorescence staining (in contract to HCN4-negative and connexin43-positive atrial cells) as shown in Figure 2A. Finally, HCN4-positive expression in studied SAN pacemaker cells is further confirmed by IF staining Figures 1C and 4D, Western blot (Figure 1E), RT-qPCR (Figure 6A), patch clamp recording of If current (Figure 6B) and sensitivity to a selective $I_f$ current inhibitor Ivabradine (Figure 7A) as well as unique morphological organization on TEM photographs, including tack of transversal tubules, disorganized sarcomere organization and a specific pattern of mitochondria distribution (Figure 1A and Figure 4A, B vs. TEM photographs of atrial and ventricular myocytes shown in Supplementary Figure 1).

#### *Intracellular confocal $Ca^{2+}$ recordings*

SAN cells were loaded with the $Ca^{2+}$ indicator dye Fluo4-AM (Invitrogen) for 15 mins and then were washed with Tyrode's solution (in mmol/L: 140 NaCl, 5.4 KCL, 1.2 $KH_2PO_4$, 1 $MgCl_2$, 5.55 D-glucose, 5 HEPES) for 15 mins at room temperature (22 ± 2°C). SAN cells with characteristic spindle and/or spider-like morphologies showing sustained spontaneous activity were selected for confocal $Ca^{2+}$ imaging using Leica SP5 confocal microscopy system following previously published methods (18,32). Confocal line scans were applied along the longest straight membrane of the cells to capture the local $Ca^{2+}$ release events with 2,000 Hz scanning speed under 63×/1.40 NA oil immersion objective, pinhole of 1 airy unit.

#### *Patch clamp studies*

**$I_{Ca,L}$ and $I_{Ca,T}$:** For calcium current recordings, electrophysiological recordings were carried out in the whole cell configuration of the patch-clamp technique at room temperature using the Axopatch 200B amplifier (Axon Instruments, Foster City, CA) with pCLAMP 10.7 software. Recording pipettes were pulled from thin-walled borosilicate glass capillaries (World Precision Instruments, Inc., Sarasota,FL) with pipette resistance of 3-5 MΩ. The recordings were filtered at 2 kHz and digitized at 20 kHz. $I_{Ca}$ currents were recorded in a voltage clamp mode of the patch-clamp technique. All solutions and buffers are indicated in mM/l. The external bath solution contained 120 Tetraethylammonium-Chloride, 10 CsCl, 10 Glucose, 10 HEPES, 1.5 $MgCl_2$, 1 $CaCl_2$, and pH 7.4 (CsOH). Intracellular pipette solution contained 100 Cs-methansulfonate, 30 CsCl, 10 HEPES, 5 EGTA, 5 Mg-ATP, 2 $MgCl_2$, and pH 7.2 (CsOH). Whole cell currents were recorded from a holding potential -80 mV with 300-ms test pulses from -40 to 50 mV, in 10 mV increments for $I_{Ca}$. For $I_{Ca,L}$ a holding potential was -40mV. We calculated $I_{Ca,T}$ by subtracting $I_{Ca,L}$ from $I_{Ca}$. Data were analyzed using Microcal Origin software (Origin Lab Corporation Northampton, MA).

**$I_f$:** $I_f$ current was recorded from spontaneously beating single cells superfused with a modified Tyrode solution containing (mM): NaCl, 115; KCl, 25; $CaCl_2$, 1.8; $MgCl_2$, 1; $MnCl_2$, 2; $BaCl_2$, 2; Hepes, 5; and glucose, 10 (pH 7.4, adjusted with NaOH). Isoproterenol was not added. Recordings were acquired using an Axopatch 200A amplifier (Molecular Devices), attached to a Digidata

1440A data acquisition system (Molecular Devices), and acquired using Clampex 10.0 at 20 KHz with a low-pass filter of 5 KHz. Electrodes were fabricated with patch glass (Warner Instruments) using a Sutter Instrument micropipette puller P-97 and were filled with an internal solution containing (mM): Potassium aspartate, 130; NaCl, 10; $CaCl_2$, 2; Hepes, 10; EGTA-KOH, 5; $MgCl_2$, 2; NaATP, 2, and NaGTP, 0.1 (pH 7.2, adjusted with KOH). The liquid junction potential resulting from the external and internal solutions used was 14 mV and was corrected after the recordings. Pipettes had resistances of 3–7 MΩ, before series resistance compensation. Steady-state current amplitudes were calculated at the end of a series of 3 s steps from -44 mV to -154 mV in 5 or 10 mV increments from a holding potential of -14 mV. Voltage steps were followed by a 1 s step to -114 mV to elicit tail currents and assess the voltage dependence of the activation. Steps were given at 10 s intervals. Some cells received 100 µM ZD 7288 to block HCN currents which eliminated most of the currents. No leak or blocker subtraction were used. The mean amplitudes of the tail currents at the end of the -114 mV step were plotted against the activation potential. Curves were fitted to a Boltzmann function, $I = A2 + [(A1-A2)/(1+e^{(V-V1/2)/z})]$, where A2 is the maximum tail current amplitude, A1 is the current offset, $V_{1/2}$ is the midpoint of activation, and z is the slope, using Origin 9.0 (OriginLab Corporation).

Fully activated $I_f$ current–voltage (*I/V*) relationships were obtained according to a previously published protocol (33).

#### *Isolated cardiomyocytes immunofluorescence labeling*

Immunolabeling was performed as previously described (19,34) on mouse SAN cells using the following primary antibodies (**Data Supplement Table I**). Briefly, isolated cardiomyocytes were fixed with 4% buffered paraformaldehyde or ice-cold methanol for 10 min. Cells fixed with paraformaldehyde were permeabilized with Triton X-100 (0.1%) for 10 min. After washing with PBS containing 0.05 % Tween-80 (PBS-T) (three 5-min washes), cells were incubated with 1 ml blocking solution (2% BSA and 2% goat serum in PBS-T) for 1.5 h at room temperature to block nonspecific binding. Subsequently, cells were incubated overnight with respective primary antibodies in blocking solution at 4°C. Excess primary antibody was washed off with the use of blocking solution (three 5-min washes). The cells were then incubated overnight with Alexa-conjugated secondary antibodies (Molecular Probes, Eugene, OR; 2 mg/ml) diluted 1:800 in blocking solution. Highly cross-absorbed goat anti-mouse Alexa Fluor 488, goat anti-rabbit Alexa Fluor 568 and goat anti rat Alexa Fluor 647 were used. The cells were then washed with PBS-T (three 5-min washes), and mounted on a coverslip. To determine nonspecific binding, control experiments with secondary antibody alone were also performed. Imaging was performed with a Leica SP5 Confocal microscope under 63×/1.40 NA oil immersion objective, pinhole of 1 airy unit. A sequential imaging pattern were applied during double staining fluorescent imaging.

To further confirm the close location of $Ca_v1.2$, Cav-3 and HCN4 in one complex triple staining experiments were performed. Labeled cells were incubated with highly cross-absorbed goat anti-mouse Alexa Fluor 488, goat anti-rabbit Alexa Fluor 568 and goat anti-rat Alexa Fluor 647 were used. Imaging was performed similarly as for double stained cells.

#### *Co-localization analysis*

Colocalization plots were used to determine the amount of different proteins that colocalized with cav-3 in the SAN cell membrane. In a 2-channel confocal image, each voxel has two intensity values (ranging from 0 to 1 after normalization in a 12-bit image), one for each red and green fluorescent signals. Voxels with normalized intensities <0.3 were considered background fluorescence and were excluded from colocalization analysis. A colocalization plot, generated with a custom-made Matlab-based computer program, displays these intensity values as a function of each other channel. By definition, two proteins are highly colocalized in a particular volume when fluorescence intensities corresponding to these two proteins are high in the voxel corresponding to this volume (1,18). Therefore, if two proteins are colocalized in many voxels, the colocalization plot will contain a significant diagonal distribution. Voxels with the highest degree of colocalization will be displayed in the upper right quadrant. In contrast, if the two proteins are not colocalized, the colocalization plot shows voxel values near each axis, with no diagonal elements present. The analysis was complemented with determination of percentage of colocalized voxels using BlobProb plugin in Fiji (http://www.fiji.sc) (35). Thresholds for each channel were determined as described above.

***Proximity ligation labeling***

Proximity ligation assay (PLA) was performed on methanol-fixed cardiomyocytes with the Duolink (inSitu) kit (Sigma-Aldrich). Cells were incubated overnight with primary antibodies against NCX and Cav-3. Secondary antibodies were PLA anti-Rabbit-plus and PLA anti-mouse-minus. PLA ligation and amplification steps were performed with far-red Duolink PLA kit according to manufacturer's instructions (Sigma-Aldrich, United States). Cell imaging was performed with 60x oil immersion Leica SP5 confocal microscope using 63x/140 NA oil immersion objective. Cells were illuminated by 633 helium-neon laser and detection was performed within 640-700 nm range. PLA analysis was performed by automatically thresholding cell images in FIJI (http://www.fiji.sc) and applying build-in "Analyze particles" plugin in FIJI to count the number of particles per area.

***Co-immunoprecipitation***

Mouse and human SAN tissue (pre-validated by immunostaining of HCN4 positive and Cx43 negative area as shown in **Data Supplement Figure S2** (36,37)) lysates as well as HEK293 cell line were used, and immunoprecipitations were carried out using anti-Cav-3 or anti-NCX, or anti-HCN4 antibodies as previously described (38); negative control of β-actin and phospholamban. Briefly, tissue samples from mice and human and HEK293 cells were flash frozen in liquid nitrogen, homogenized and lysed using buffer containing 150 mM NaCl, 25 mM Tris·HCl, 10 mM NaEGTA, 20 mM NaEDTA, and supplemented with 0.5% CHAPS and 1x protease inhibitor cocktail (B14001, bimake). The lysate (500 μg) was sonicated, rotated at 4ºC for 2 hours and centrifuged at 16,000 × g for 15 min at 4ºC. The soluble supernatant was incubated with aforementioned antibodies (20 μg) at 4ºC for 2 hours and then incubated with 25 μL (Protein A Mag Sepharose/Protein G Mag Sepharose, 28944006 /28944008, GE Life Sciences; 1:1 Protein A/G) at 4ºC for 2 hours, followed by incubation with anti-Cav-3, anti-NCX and anti-HCN4 (antibody details see **Data Supplement Table I**) in a total of 300-400 μL (depending on protein concentration) of lysate. After flowthrough collection, samples were eluted with 2.5% Acetic Acid. Laemmli Buffer (161-0747, Bio-Rad) and DTT were added and all samples were incubated at 65ºC for 10 min. Immune complexes were analyzed by SDS-PAGE (4-20% gradient gels, Bio-Rad) and Western blot by probing with appropriate antibodies.

***Transmission Electron Microscopy***

SAN tissue was isolated as previously described in SAN cell isolation section and fixed in the following fixative: 2.5% glutaraldehyde, 2.0% paraformaldehyde, and 0.2 mol/l cacodylate buffer for 24 - 48 hours. The samples were rinsed in the same buffer, post-fixed in 1% osmium tetroxide, dehydrated in a graded ethanol series, rinsed in propylene oxide, and embedded in Epon 812 substitute. After resin polymerization, the samples were then sliced into 70-nm sections with a Leica EM UC6 ultramicrotome and placed on 200 mesh transmission electron microscopy grids. The samples were post-stained in 8% uranyl acetate in 50% EtOH and Reynold's lead citrate, viewed on a Philips CM120 transmission electron microscope, and documented with a SIS MegaView III digital camera.

Electron microscopy images were analyzed by using the NIH ImageJ software. A threshold size for individual caveolae was set between 50 and 100 nm. The number of caveolae was counted as per unit length (μm) of myocyte sarcolemmal membranes from a series of random electron microscopy micrographs. Images with recognized sarcolemmal reticulum (SR) from WT, HF and Cav-3KO SAN tissue were selected and the distance between SR membrane to the closest sarcolemmal membrane was calculated.

***Statistical Analysis***

Student *t* test was used in 2-group comparisons. Multiple groups of normally distributed data of similar variance were compared by 1 - or 2 - way ANOVA. For multiple comparisons, the Bonferroni corrected *P* value is shown. All statistical analyses were performed using GraphPad Prism 5 (GraphPad Software). $P<0.05$ was considered statistically significant. Values were presented as mean ± SEM.

***Mathematical modeling and simulation***

<u>*Subcellular $Ca^{2+}$ signaling model*</u>. We used an established 3D model of rabbit ventricular $Ca^{2+}$ signaling (39,40) as the basis for developing an analogous model of the murine SAN cell that describes subcellular stochastic properties of individual SR $Ca^{2+}$ release units (CRUs). **Figure 5A** depicts the model schematic: in each peripheral CRU, i.e., a CRU coupled to external membrane, there are 5 $Ca^{2+}$ compartments (cytosolic, submembrane, cleft space, network SR, and junctional SR), whereas central CRUs are not coupled to external membrane and have less RyRs vs. the peripheral CRUs (**Data Supplement Table II**). We assigned the first two CRUs close to the surface membrane as peripheral CRUs and the rest as central CRUs. CRUs are coupled by $Ca^{2+}$ diffusions in submembrane, cytosol and the network SR. We modified this model to describe the structural data, global cellular dimensions, and electrical capacitance of typical mouse sinoatrial node cells based on (41,42). The new 3D SAN cell model has a dimension of 60 μm × 7 μm × 7 μm with a capacitance of 25 pF and comprises 2176 (34 × 8 × 8) CRUs. Detailed model updates and parameters are in the ***Appendix*** and **Data Supplement Table II**. Briefly, we updated $Ca^{2+}$ current models in the original 3D rabbit $Ca^{2+}$ signaling model to account for the mouse SAN cell-specific properties (**Data Supplement Table II**). Specifically, we replaced the model of L-type

$Ca^{2+}$ current ($I_{CaL}$) in the original 3D $Ca^{2+}$ signaling model with a Markov-equivalent implementation of the $I_{CaL}$ formulation from (41) using the approach employed in (43). The $Na^+$/$Ca^{2+}$ exchanger current ($I_{NCX}$) and T-type $Ca^{2+}$ current ($I_{CaT}$) of the new model were simulated using the formula in (41). Furthermore, we updated the RyR release model to reflect enhanced SAN RyR $Ca^{2+}$ sensitivity compared to the original model for rabbit ventricle. Our model does not explicitly account for functional caveolar or extra-caveolar $Ca^{2+}$ compartments, which would be intermediate (in terms of size and $Ca^{2+}$ concentration levels) between the cleft and subsarcolemmal spaces. Localizations of L-type $Ca^{2+}$ channels, T-type $Ca^{2+}$ channels and NCX were based on colocalization data collected in **Figure 1**.

*Coupled $Ca^{2+}$ and $V_m$ model*. To build a 3D mouse SAN myocyte model incorporating detailed local $Ca^{2+}$ signaling and electrophysiology, we coupled the new 3D subcellular $Ca^{2+}$ signaling model with a mouse SAN AP model (0D AP model). The 0D AP model was updated from (41) as previously described in (42) (details in the Appendix). For each time step, all $Ca^{2+}$-dependent transmembrane currents ($I_{CaL}$, $I_{CaT}$, $I_{NCX}$, Sarcolemmal membrane background $Ca^{2+}$ current and $Ca^{2+}$ pump - $I_{CaBk}$ and $I_{CaP}$) and intracellular $Ca^{2+}$ handling were calculated using the 3D $Ca^{2+}$ signaling model, and the remaining transmembrane currents ($Na^+$ and $K^+$ currents) were computed using the 0D AP model. The membrane voltage ($V_m$) was then computed from the total transmembrane current using the electrical circuit model as in (41,42). The intracellular $Na^+$ concentration was fixed to 10 mM as in (39). Of note, spatial models of rabbit SAN myocyte incorporating stochastic descriptions of $Ca^{2+}$ handling have been previously constructed, reveal key contributions of subcellular $Ca^{2+}$ release units and their coupling with sarcolemmal ionic proteins to maintaining normal and robust pacemaking activity (44-47).

*Simulation protocols*. Our new 3D SAN cell model was implemented in C++. The ordinary differential equations are solved using the forward Euler method except that $I_{CaL}$ and RyR flux were solved stochastically as in (39,40,43). The time step was 0.01 ms as in (39). A period of 30 sec was simulated and the data from the last 20 sec was analyzed.

*Source code*. The source code of our new 3D SAN cell model can be accessed from http://elegrandi.wixsite.com/grandilab/downloads and https://github.com/drgrandilab.

### *Appendix*

**0D mouse SAN AP model** We used an updated version of the Kharche et al. model (41) model of mouse sinoatrial node myocyte (SAM), as described in (42) based on experimental data obtained in the Proenza lab in 2-3 months old male C58BL/6J mice at physiological temperature. In the updated model, the parameters for $I_{CaL}$, $I_f$, $I_{to}$ and $I_{sus}$ were initially manually tuned to match our voltage-clamp experiments; then, an optimization based on reverse multivariable regression (non-linear iterative partial least squares method, as in (48)) was performed to identify model parameters (**Data Supplement Table II**) that best matched various recorded AP waveform parameters (including cycle length, $APD_{90}$ and $APD_{50}$, AP amplitude, diastolic depolarization rate and duration, upstroke velocity, maximum diastolic potential).

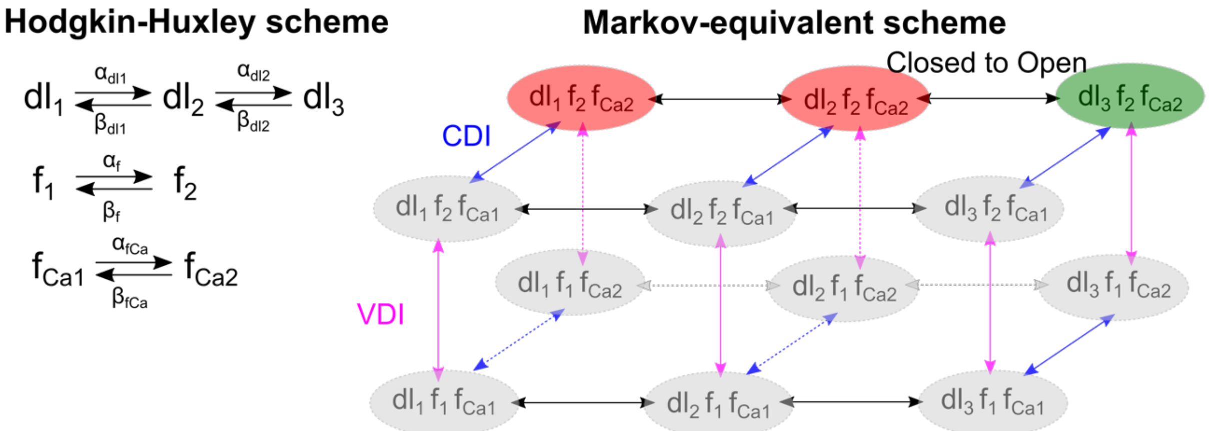


***Modeling scheme of L-type Ca$^{2+}$ current.*** Left panel: a Hodgkin-Huxley scheme of $I_{CaL}$ modified from (41) with the addition of *dl₃* state; Right panel: the Markov-equivalent scheme for the Hodgkin-Huxley $I_{CaL}$ model. CDI – Ca$^{2+}$-dependent inactivation. VDI – voltage-dependent inactivation. The conceptualization of conversion between the two schemes is detailed in (43).

**L-type Ca$^{2+}$ current:** The L-type Ca$^{2+}$ channels are assumed to be located in the cleft area in the peripheral CRUs and closely coupled with RyRs. We modified the Hodgkin-Huxley type model of $I_{CaL}$ in (41) to a Markov-equivalent scheme using the approach described in (43). To reflect the difference in maximum open probability between the Hodgkin-Huxley type model (close to 100%) and experimental measurements (~5%), we introduced an additional state ($dl_3$) in the Markov-equivalent scheme above as detailed in (43). The model is then solved stochastically as in (43). The $I_{CaL}$ current is described by

- Activation

$$dl_{inf} = \frac{1}{1 + \exp\left(-\frac{V + 13.5}{6.0}\right)}$$

$$a = -28.39 \cdot \frac{\mathrm{V} + 35.0}{\exp\left(-\frac{V + 35.0}{2.5}\right) - 1.0} - 84.9 \cdot \frac{V}{\exp(-0.208 \cdot V) - 1.0}$$

$$b = 11.43 \cdot \frac{V - 5}{\exp\big(0.4(V - 5)\big) - 1.0}$$

$$\tau = \frac{2000.0}{a + b}$$

$$\alpha_{dl} = \frac{dl_{inf}}{\tau}$$

$$\beta_{dl} = \frac{1 - dl_{inf}}{\tau}$$

$$\alpha_{dl2} = 0.3$$

$$\beta_{dl2} = 6.0$$

- Voltage-dependent inactivation

$$f_{inf} = \frac{1}{1 + \exp\left(\frac{V + 35.0}{7.3}\right)}$$

$$\tau_f = 7.4 + 45.77 \cdot \exp\left(-0.5\left(\frac{V + 28.1}{11.0}\right)^2\right)$$
$$\alpha_f = \frac{f_{inf}}{\tau_f}$$
$$\beta_f = \frac{1 - f_{inf}}{\tau_f}$$

- $Ca^{2+}$-dependent inactivation

$$fCa_{inf} = \frac{1}{\left(1 + \frac{C_p}{3.0}\right)^2}$$
$$\alpha_{fCa} = \frac{fCa_{inf}}{\tau_{fCa}}$$
$$\beta_{fCa} = \frac{1 - fCa_{inf}}{\tau_{fCa}}$$

- Local $I_{CaL}$

$$P_O = dl_3 f_2 f_{Ca2}$$
$$N_{L,O} = \sum P_O$$
$$I_{CaL} = N_{L,O} \cdot ica$$

where $P_O$ is a binary variable describing whether a LTCC channel is open or closed, $N_{L,O}$ is the total number of open LTCC, and $ica$ is the unitary current and computed as

$$ica = 4P_{Ca}zF \cdot \frac{\gamma_i C_p \cdot 0.001 \cdot \exp(2 \cdot z) - \gamma_O [Ca]_O}{\exp(2 \cdot z) - 1.0}$$

where

$$z = \frac{VF}{RT}$$

**T-type $Ca^{2+}$ current (*$I_{CaT}$*):** The $I_{CaT}$ formula from the Kharche et al. model (41) was coupled with the 3D $Ca^{2+}$ signaling model. We evenly distributed the $Ca^{2+}$ flux carried by $I_{CaT}$ to the peripheral CRUs, with 50% of the proteins in the cleft and 50% in the submembrane compartment for each CRU (**Data Supplement Table II**).

**$Na^+/Ca^{2+}$ exchanger (*$I_{NCX}$*):** The NCX model of the 3D $Ca^{2+}$ signaling model was replaced with that from (41). In the new baseline 3D $Ca^{2+}$ model of SAN, NCX was equally distributed among the peripheral CRUs, with 50% of the proteins in the cleft and 50% in the submembrane compartment for each CRU (**Data Supplement Table II**).

**SR $Ca^{2+}$ release and leak:** To simulate the $Ca^{2+}$ release from SR via RyR, we developed a new model of RyR $Ca^{2+}$ release (*schematic at right*) based on the RyR models from (39,49). The new model adds $[Ca]_{SR}$-dependence to closed-to-open transitions rates and higher sensitivity to cleft $Ca^{2+}$ compared with the Sato-Bers model (39). Also, in the new model, the SR $Ca^{2+}$ leak was redirected to the cleft space instead of the cytosolic space in (40). The new RyR model is described by

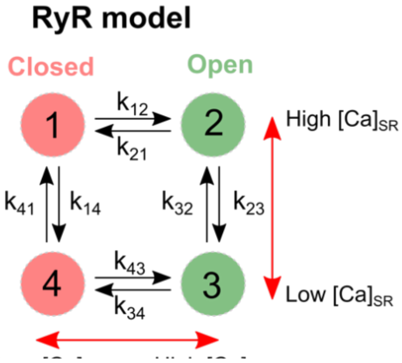

***RyR gating model schematic***

$$k_{CaSR} = Max_{SR} - \frac{Max_{SR} - Min_{SR}}{1 + \left(\frac{EC_{50SR}}{[Ca]_{SRj}}\right)^{2.5}}$$

$$k_{OCaSR} = \frac{1}{k_{CaSR}}$$

$$k_{12} = k_{OCaSR} K_u \frac{C_p^2}{C_p^2 + K_{cp}^2} + 0.00001$$

$$k_{43} = k_{OCaSR} K_b \frac{C_p^2}{C_p^2 + K_{cp}^2} + 0.00001$$

$$k_{14} = \frac{\widehat{M}(C_p) B_{CSQN}}{\tau_b B_{CSQN0}}$$

$$k_{23} = \frac{\widehat{M}(C_p) B_{CSQN}}{\tau_b B_{CSQN0}}$$

where $\widehat{M}$, $B_{CSQN}$ and $B_{CSQN0}$ are kept the same as in (39)

$$k_{21} = \frac{1}{\tau_{c1} k_{OCaSR}}$$

$$k_{34} = \frac{1}{\tau_{c2} k_{OCaSR}}$$

$$k_{41} = \frac{1}{\tau_u} \cdot k_{OCaSR}$$

$$k_{32} = \frac{k_{41} \cdot k_{12}}{k_{43}}$$

**$Ca^{2+}$ buffering:** Similar to the previous studies (39,40), the buffering of $Ca^{2+}$ in each compartment was modeled using an instantaneous buffering function, with an exception that troponin-$Ca^{2+}$ was modelled dynamically. In our 3D model of SAN, we removed buffering of SR, Myosin ($Ca^{2+}$), and Myosin (Mg) in the cleft and submembrane compartments as these buffers were absent in these compartments in the previous models (41,49).

**Background sarcolemmal membrane $Ca^{2+}$ flux ($I_{CaBk}$):** Similar to Sato-Bers model (39), $I_{CaBk}$ was modeled as

$$I_{CaBk} = G_{CaBk}(V - E_{Ca})$$

$$E_{Ca} = \frac{RT}{F} ln \frac{[Ca]_O}{c_x}$$

where $C_x$ is $Ca^{2+}$ concentration in the cleft or submembrane space. The compartmentalization of $I_{CaBk}$ was kept the same as in (49) (**Data Supplement Table II**).

**Sarcolemmal membrane $Ca^{2+}$ pump ($I_{CaP}$):** We added $I_{CaP}$ as did in Sato-Bers model (39) and compartmentalized the current as in (49). $I_{CaP}$ is computed by

$$I_{CaP} = \frac{Q_{CaP} V_{max}}{1 + \left(\frac{K_{mCaP}}{C_x}\right)^H}$$

where $C_x$ is the $Ca^{2+}$ concentration in the cleft or submembrane space. The compartmentalization of $I_{CaP}$ was kept the same as in (49). Parameters for both $I_{CaBK}$ and $I_{CaP}$ was tuned so that the two whole-cell currents from the 3D $Ca^{2+}$ signaling model were comparable to those from the updated 0D electrophysiological model (**Data Supplement Table II**).

### References


1. Glukhov AV, Fedorov VV, Kalish PW et al. Conduction remodeling in human end-stage nonischemic left ventricular cardiomyopathy. Circulation 2012;125:1835-47.
2. Lou Q, Fedorov VV, Glukhov AV, Moazami N, Fast VG, Efimov IR. Transmural heterogeneity and remodeling of ventricular excitation-contraction coupling in human heart failure. Circulation 2011;123:1881-90.
3. Glukhov AV, Fedorov VV, Lou Q et al. Transmural dispersion of repolarization in failing and nonfailing human ventricle. Circ Res 2010;106:981-91.
4. Fedorov VV, Glukhov AV, Chang R et al. Optical mapping of the isolated coronary-perfused human sinus node. J Am Coll Cardiol 2010;56:1386-94.
5. Li N, Csepe TA, Hansen BJ et al. Molecular Mapping of Sinoatrial Node HCN Channel Expression in the Human Heart. Circ Arrhythm Electrophysiol 2015;8:1219-27.
6. Li N, Hansen BJ, Csepe TA et al. Redundant and diverse intranodal pacemakers and conduction pathways protect the human sinoatrial node from failure. Science translational medicine 2017;9.
7. Keith A, Flack M. The form and nature of the muscular connection between the primary divisions of the vertebrate heart. J Anat Physiol 1906;41:172-189.
8. James TN. Anatomy of the human sinus node. Anat Rec 1961;141:109-39.
9. Miller WL, Grill DE, Mullan BP. Comparison of Blood Volume Profiles in Heart Failure With Preserved and Reduced Ejection Fractions: Sex Makes a Difference. Circ Heart Fail 2024;17:e010906.

10. Farooqui N, Killian JM, Smith J, Redfield MM, Dunlay SM. Advanced Heart Failure Characteristics and Outcomes in Women and Men. J Am Heart Assoc 2024;13:e033374.
11. Lindsey ML, Bolli R, Canty JM et al. Guidelines for experimental models of myocardial ischemia and infarction. Am J Physiol Heart Circ Physiol 2018;314:H812-H838.
12. Sohal DS, Nghiem M, Crackower MA et al. Temporally regulated and tissue-specific gene manipulations in the adult and embryonic heart using a tamoxifen-inducible Cre protein. Circ Res 2001;89:20-5.
13. Markandeya YS, Feng L, Vaidyanathan R et al. Caveolin-3 Regulates Cardiac Repolarization by Integrated Regulation of Multiple Ionic Currents. Circulation 2013;128:A15009.
14. Wright PT, Diakonov I, Pannell L et al. Cardiomyocyte membrane structure and cAMP compartmentation produce anatomical variation in β2AR-cAMP responsiveness in murine hearts. Cell Reports 2018;23:459-469.
15. Medvedev RY, Turner DGP, DeGuire FC et al. Caveolae-associated cAMP/$Ca2^{+}$-mediated mechano-chemical signal transduction in mouse atrial myocytes. J Mol Cell Cardiol 2023;184:75-87.
16. Tyan L, Turner D, Komp KR, Medvedev RY, Lim E, Glukhov AV. Caveolin-3 is required for regulation of transient outward potassium current by angiotensin II in mouse atrial myocytes. Am J Physiol Heart Circ Physiol 2021;320:H787-H797.
17. Liu YH, Wang D, Rhaleb NE et al. Inhibition of p38 mitogen-activated protein kinase protects the heart against cardiac remodeling in mice with heart failure resulting from myocardial infarction. J Card Fail 2005;11:74-81.
18. Egorov YV, Lang D, Tyan L et al. Caveolae-Mediated Activation of Mechanosensitive Chloride Channels in Pulmonary Veins Triggers Atrial Arrhythmogenesis. Journal of the American Heart Association 2019;8:e012748.
19. Glukhov AV, Kalyanasundaram A, Lou Q et al. Calsequestrin 2 deletion causes sinoatrial node dysfunction and atrial arrhythmias associated with altered sarcoplasmic reticulum calcium cycling and degenerative fibrosis within the mouse atrial pacemaker complex1. Eur Heart J 2015;36:686-97.
20. Glukhov AV, Fedorov VV, Anderson ME, Mohler PJ, Efimov IR. Functional anatomy of the murine sinus node: high-resolution optical mapping of ankyrin-B heterozygous mice. Am J Physiol Heart Circ Physiol 2010;299:H482-91.
21. Lang D, Glukhov AV. High-resolution Optical Mapping of the Mouse Sino-atrial Node. J Vis Exp 2016.
22. Ripplinger CM, Glukhov AV, Kay MW et al. Guidelines for assessment of cardiac electrophysiology and arrhythmias in small animals. Am J Physiol Heart Circ Physiol 2022;323:H1137-H1166.
23. Lou Q, Glukhov AV, Hansen B et al. Tachy-brady arrhythmias: the critical role of adenosine-induced sinoatrial conduction block in post-tachycardia pauses. Heart Rhythm 2013;10:110-8.
24. Egom EE, Vella K, Hua R et al. Impaired sinoatrial node function and increased susceptibility to atrial fibrillation in mice lacking natriuretic peptide receptor C. J Physiol 2015;593:1127-46.
25. Krishnaswamy PS, Egom EE, Moghtadaei M et al. Altered parasympathetic nervous system regulation of the sinoatrial node in Akita diabetic mice. J Mol Cell Cardiol 2015;82:125-35.

26. Sharpe EJ, St Clair JR, Proenza C. Methods for the Isolation, Culture, and Functional Characterization of Sinoatrial Node Myocytes from Adult Mice. J Vis Exp 2016.
27. Liu J, Dobrzynski H, Yanni J, Boyett MR, Lei M. Organisation of the mouse sinoatrial node: structure and expression of HCN channels. Cardiovasc Res 2007;73:729-38.
28. Swaminathan PD, Purohit A, Soni S et al. Oxidized CaMKII causes cardiac sinus node dysfunction in mice. J Clin Invest 2011;121:3277-88.
29. Verheijck EE, van Kempen MJ, Veereschild M, Lurvink J, Jongsma HJ, Bouman LN. Electrophysiological features of the mouse sinoatrial node in relation to connexin distribution. Cardiovasc Res 2001;52:40-50.
30. Golovko V, Gonotkov M, Lebedeva E. Effects of 4-aminopyridine on action potentials generation in mouse sinoauricular node strips. Physiol Rep 2015;3.
31. Ding Y, Lang D, Yan J et al. A phenotype-based forward genetic screen identifies Dnajb6 as a sick sinus syndrome gene. Elife 2022;11.
32. Lang D, Sato D, Jiang Y, Ginsburg KS, Ripplinger CM, Bers DM. Calcium-Dependent Arrhythmogenic Foci Created by Weakly Coupled Myocytes in the Failing Heart. Circulation research 2017.
33. Alvarez-Baron CP, Klenchin VA, Chanda B. Minimal molecular determinants of isoform-specific differences in efficacy in the HCN channel family. J Gen Physiol 2018;150:1203-1213.
34. Le Scouarnec S, Bhasin N, Vieyres C et al. Dysfunction in ankyrin-B-dependent ion channel and transporter targeting causes human sinus node disease. Proc Natl Acad Sci U S A 2008;105:15617-22.
35. Fletcher PA, Scriven DR, Schulson MN, Moore ED. Multi-image colocalization and its statistical significance. Biophys J 2010;99:1996-2005.
36. Glukhov AV, Hage LT, Hansen BJ et al. Sinoatrial node reentry in a canine chronic left ventricular infarct model: role of intranodal fibrosis and heterogeneity of refractoriness. Circ Arrhythm Electrophysiol 2013;6:984-94.
37. Lou Q, Hansen BJ, Fedorenko O et al. Upregulation of adenosine A1 receptors facilitates sinoatrial node dysfunction in chronic canine heart failure by exacerbating nodal conduction abnormalities revealed by novel dual-sided intramural optical mapping. Circulation 2014;130:315-24.
38. Balijepalli RC, Foell JD, Hall DD, Hell JW, Kamp TJ. Localization of cardiac L-type Ca(2+) channels to a caveolar macromolecular signaling complex is required for beta(2)-adrenergic regulation. Proc Natl Acad Sci U S A 2006;103:7500-5.
39. Sato D, Bers DM. How does stochastic ryanodine receptor-mediated Ca leak fail to initiate a Ca spark? Biophys J 2011;101:2370-9.
40. Restrepo JG, Weiss JN, Karma A. Calsequestrin-mediated mechanism for cellular calcium transient alternans. Biophys J 2008;95:3767-89.
41. Kharche S, Yu J, Lei M, Zhang H. A mathematical model of action potentials of mouse sinoatrial node cells with molecular bases. Am J Physiol Heart Circ Physiol 2011;301:H945-63.
42. Morotti S, Ni H, Peters CH et al. Intracellular Na(+) Modulates Pacemaking Activity in Murine Sinoatrial Node Myocytes: An In Silico Analysis. Int J Mol Sci 2021;22.
43. Song Z, Ko CY, Nivala M, Weiss JN, Qu Z. Calcium-voltage coupling in the genesis of early and delayed afterdepolarizations in cardiac myocytes. Biophys J 2015;108:1908-21.

44. Maltsev AV, Stern MD, Maltsev VA. Disorder in Ca2+ release unit locations confers robustness but cuts flexibility of heart pacemaking. J Gen Physiol 2022;154.
45. Maltsev AV, Maltsev VA, Mikheev M et al. Synchronization of stochastic Ca²(+) release units creates a rhythmic Ca²(+) clock in cardiac pacemaker cells. Biophys J 2011;100:271-83.
46. Stern MD, Maltseva LA, Juhaszova M, Sollott SJ, Lakatta EG, Maltsev VA. Hierarchical clustering of ryanodine receptors enables emergence of a calcium clock in sinoatrial node cells. J Gen Physiol 2014;143:577-604.
47. Maltsev AV, Yaniv Y, Stern MD, Lakatta EG, Maltsev VA. RyR-NCX-SERCA local cross-talk ensures pacemaker cell function at rest and during the fight-or-flight reflex. Circ Res 2013;113:e94-e100.
48. Sarkar AX, Sobie EA. Regression analysis for constraining free parameters in electrophysiological models of cardiac cells. PLoS Comput Biol 2010;6:e1000914.
49. Shannon TR, Wang F, Puglisi J, Weber C, Bers DM. A mathematical treatment of integrated Ca dynamics within the ventricular myocyte. Biophys J 2004;87:3351-71.
50. Markandeya YS, Gregorich ZR, Feng L et al. Caveolin-3 and Caveolae regulate ventricular repolarization. J Mol Cell Cardiol 2023;177:38-49.
51. Busija AR, Patel HH, Insel PA. Caveolins and cavins in the trafficking, maturation, and degradation of caveolae: implications for cell physiology. American journal of physiology Cell physiology 2017;312:C459-C477.
52. Balijepalli RC, Delisle BP, Balijepalli SY et al. Kv11.1 (ERG1) K+ channels localize in cholesterol and sphingolipid enriched membranes and are modulated by membrane cholesterol. Channels (Austin) 2007;1:263-72.

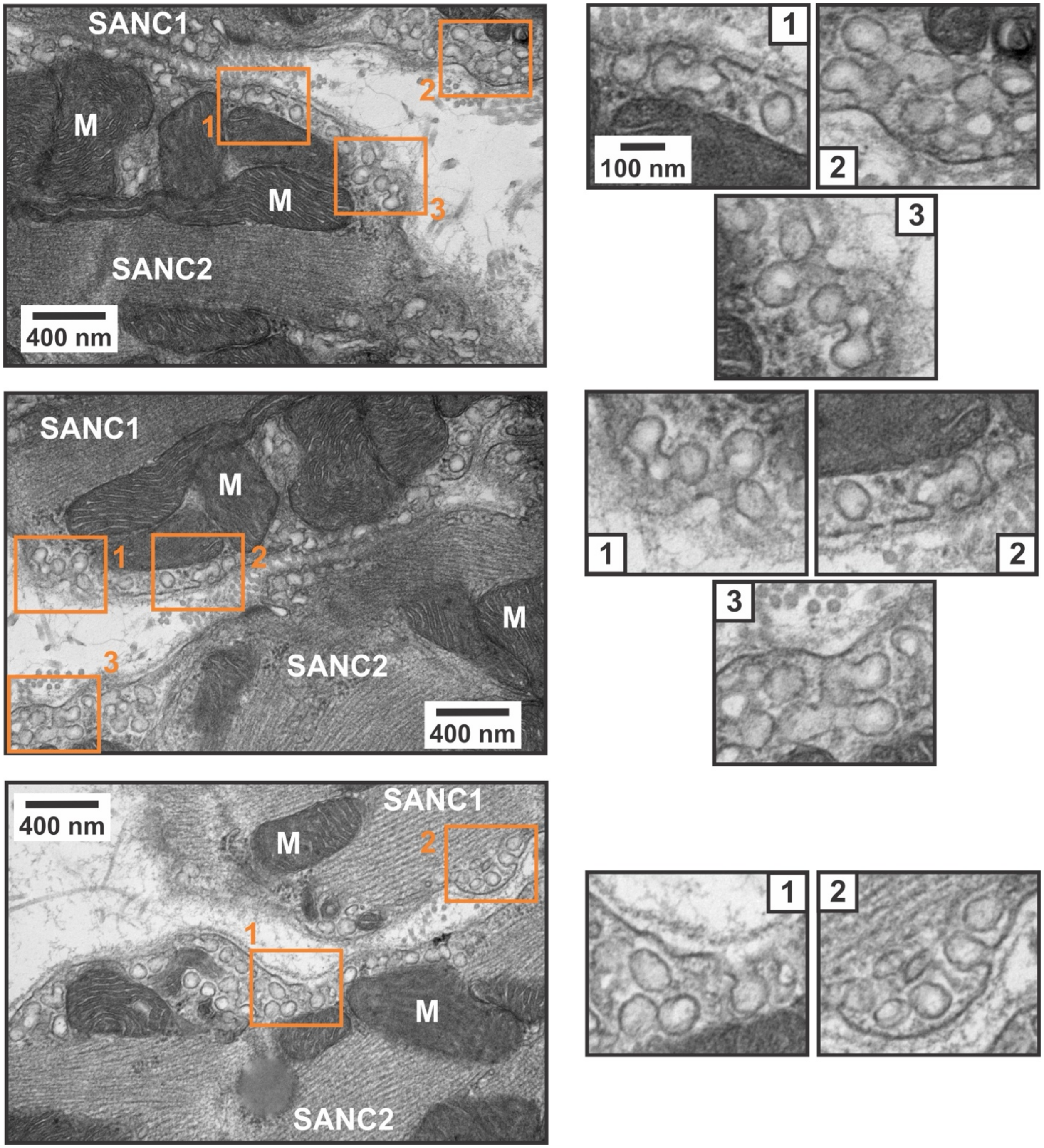


**Supplementary Figure S1.** Representative examples of caveolae structures in sinoatrial nodal (SAN) cells (SANC) visualized by transmission electron microscopy (TEM) imaging from wild type SAN tissue samples.

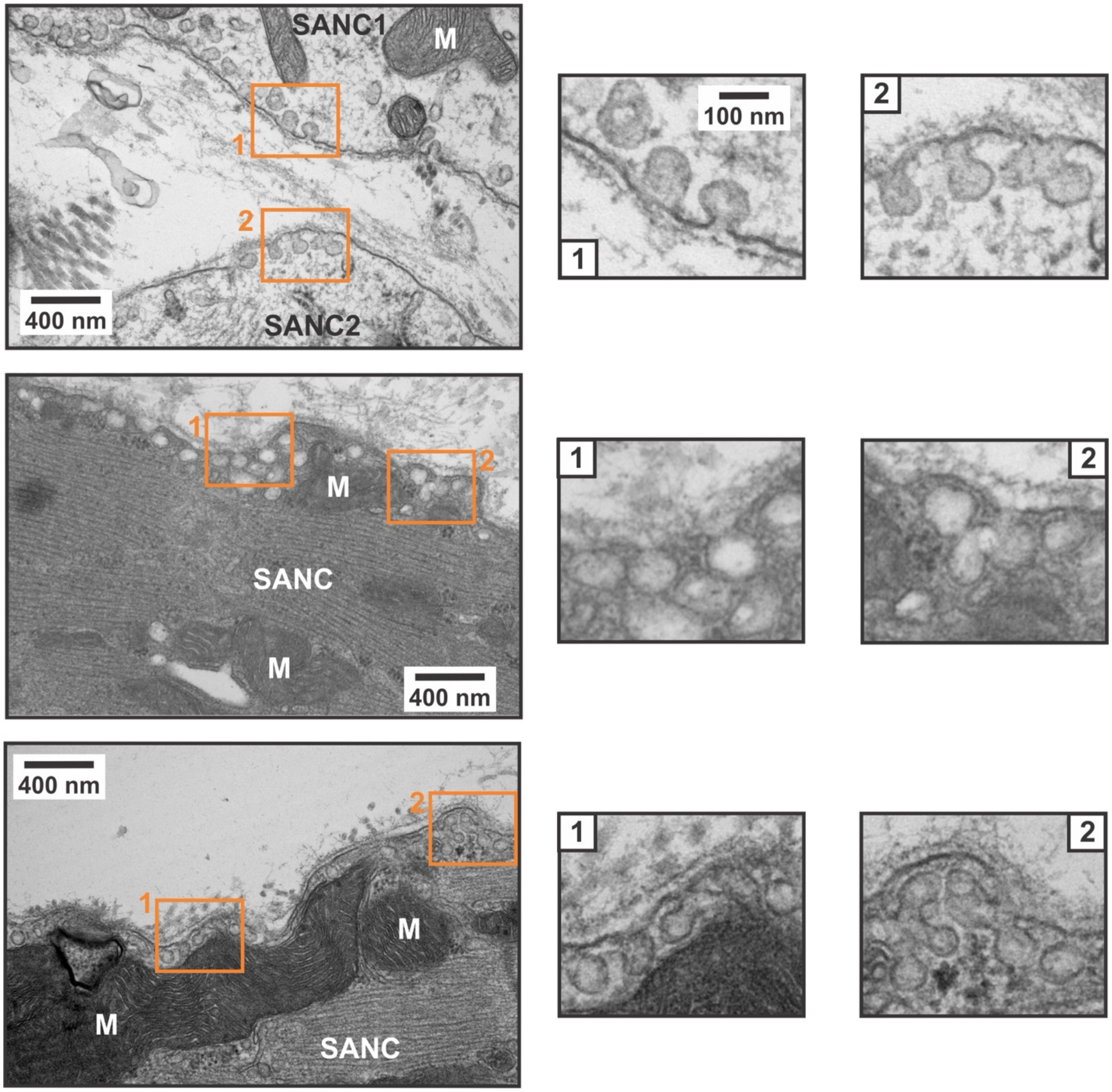


**Supplementary Figure S2.** Additional representative examples of caveolae structures in sinoatrial nodal (SAN) cells (SANC) visualized by transmission electron microscopy (TEM) imaging from wild type SAN tissue samples.

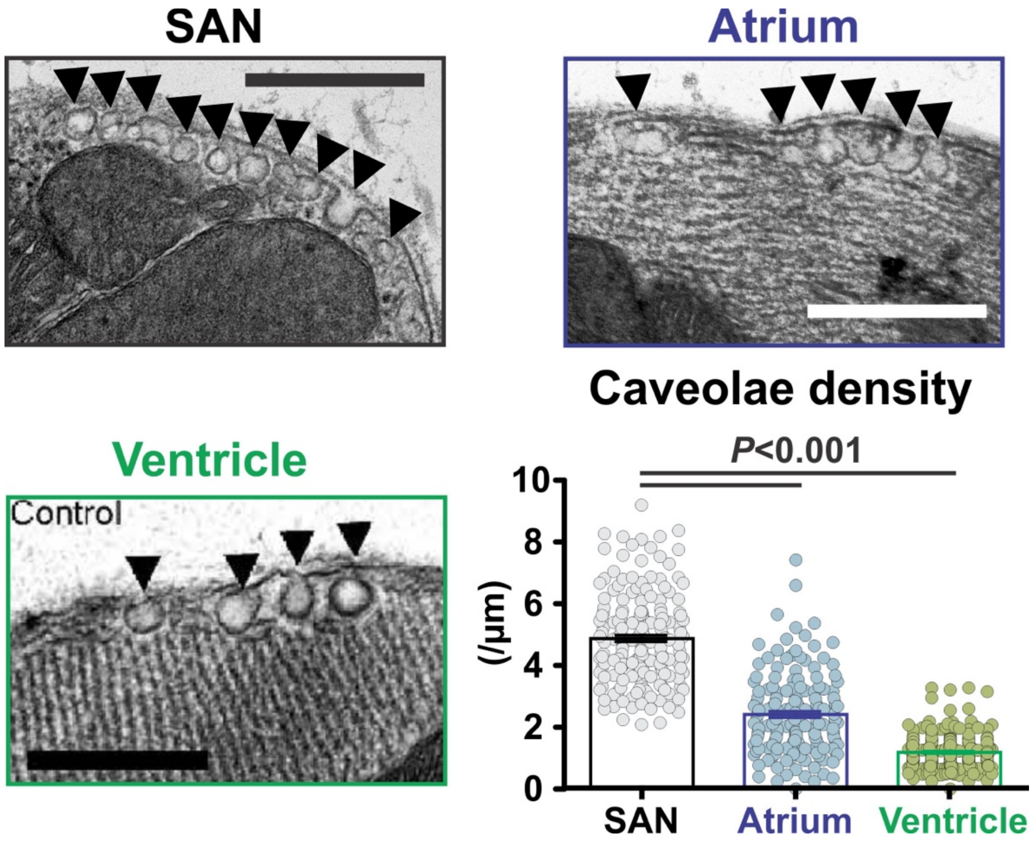


**Supplementary Figure S3.** Caveolae density in control (wild type) mouse SAN, right atrial, and left ventricular myocardium. Representative transmission electron micrographs are shown (scale bar 500 nm). Morphological caveolae are denoted by arrow heads. Caveolae per micron membrane from SAN (n=183 cells from 5 mice), right atrium (n=164 cells from 5 mice), and left ventricle (n = 242 cells from 3 mice). Ventricular TEM image and caveolae density data is reproduced from (50) with permission. P-values were determined by one-way ANOVA with Bonferroni correction.

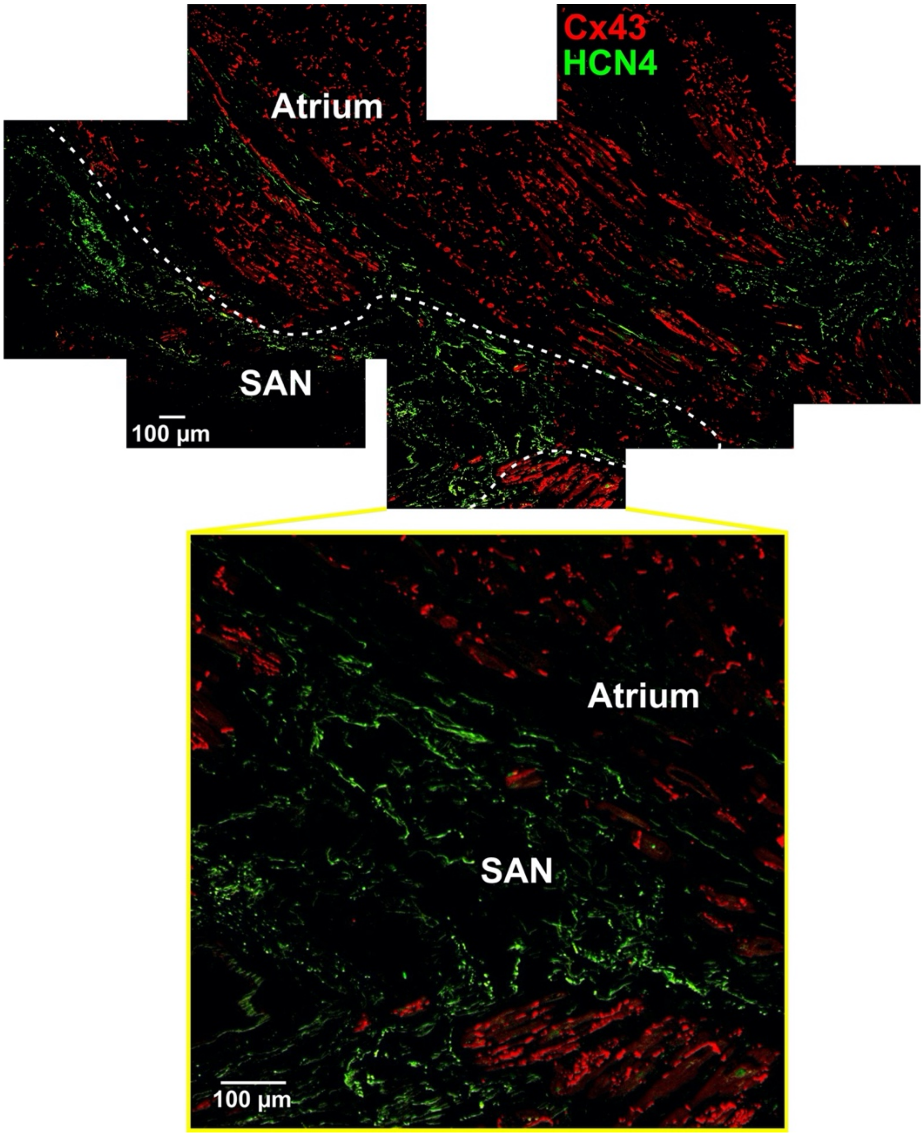


**Supplementary Figure S4.** Mosaic scanning of healthy human SAN tissue slide with fluorescent staining of HCN4 (green) and Cx43 (red). The immunofluorescent staining was used to confirm the SAN area in the human tissue as HCN4 positive and Cx43 negative myocardium.

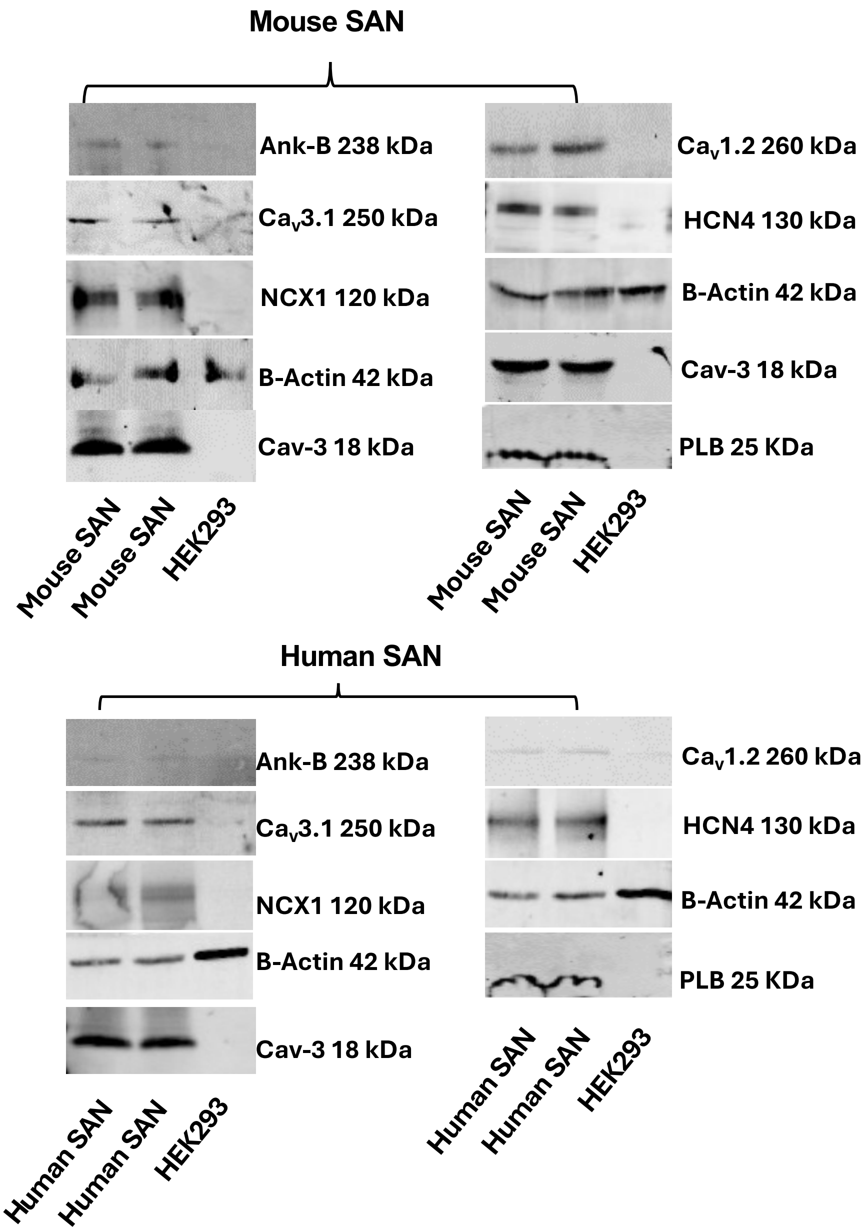


**Supplementary Figure S5.** Western blot antibodies testing in mouse and human sinoatrial node (SAN) tissue lysates for HCN4, NCX1, $Ca_v1.2$ and $Ca_v3.1$ proteins. HCN4 was used to confirm SAN myocardium. HEK293 cells were used as a negative control. Caveolin-3 (Cav-3), ankyrin-B (Ank-B) and phospholamban (PLB) were used as muscle-specific proteins. B-actin was used for loading control.

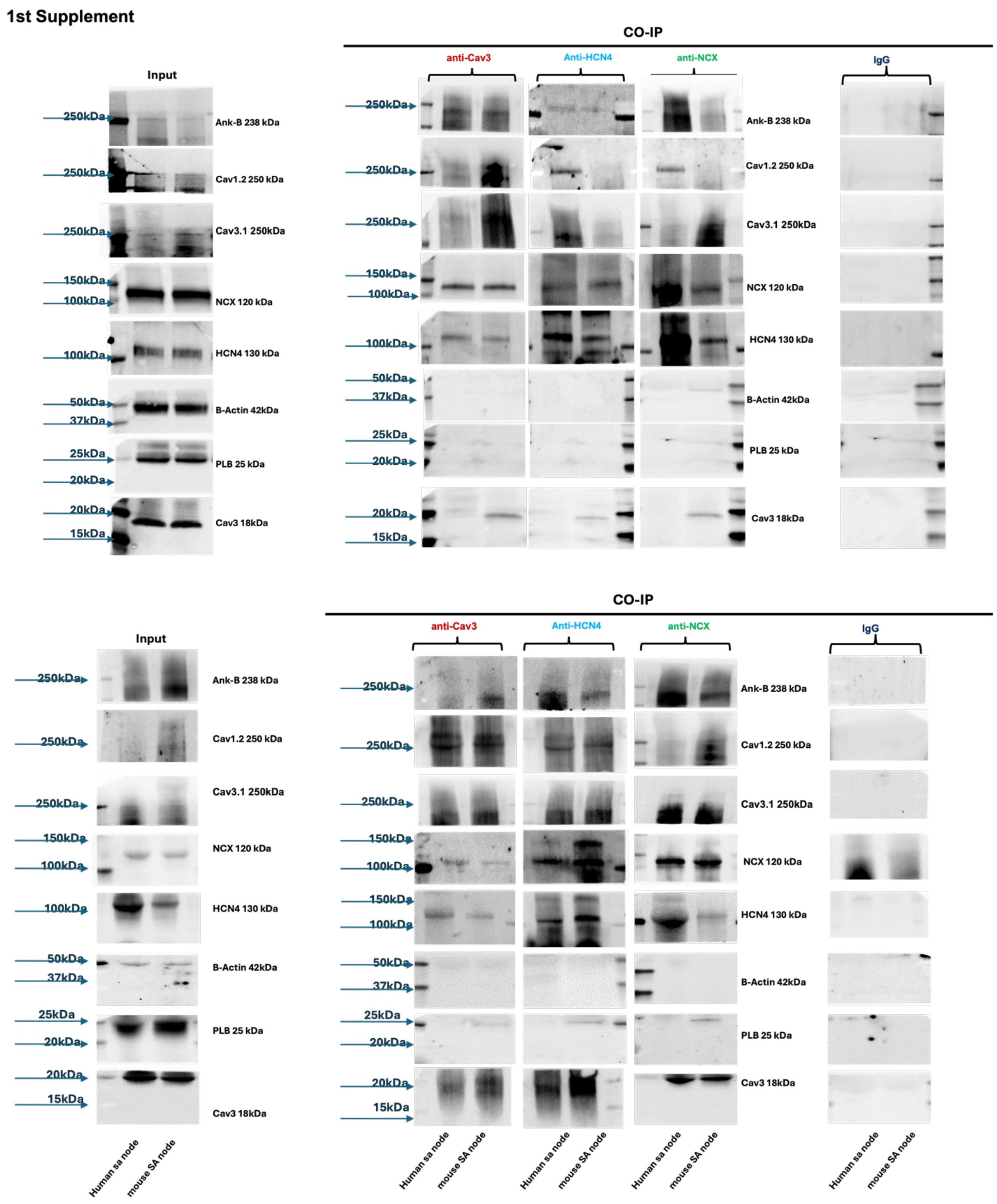


**Supplementary Figure S6.** Co-immunoprecipitation (Co-IP) Western blot experiments performed on mouse and human sinoatrial nodal (SAN) tissues. Top and bottom figures represent, respectively, two biological replicates. The Co-IPs for Fig. 1E and the first supplemental figure above were conducted separately from the same lysate.

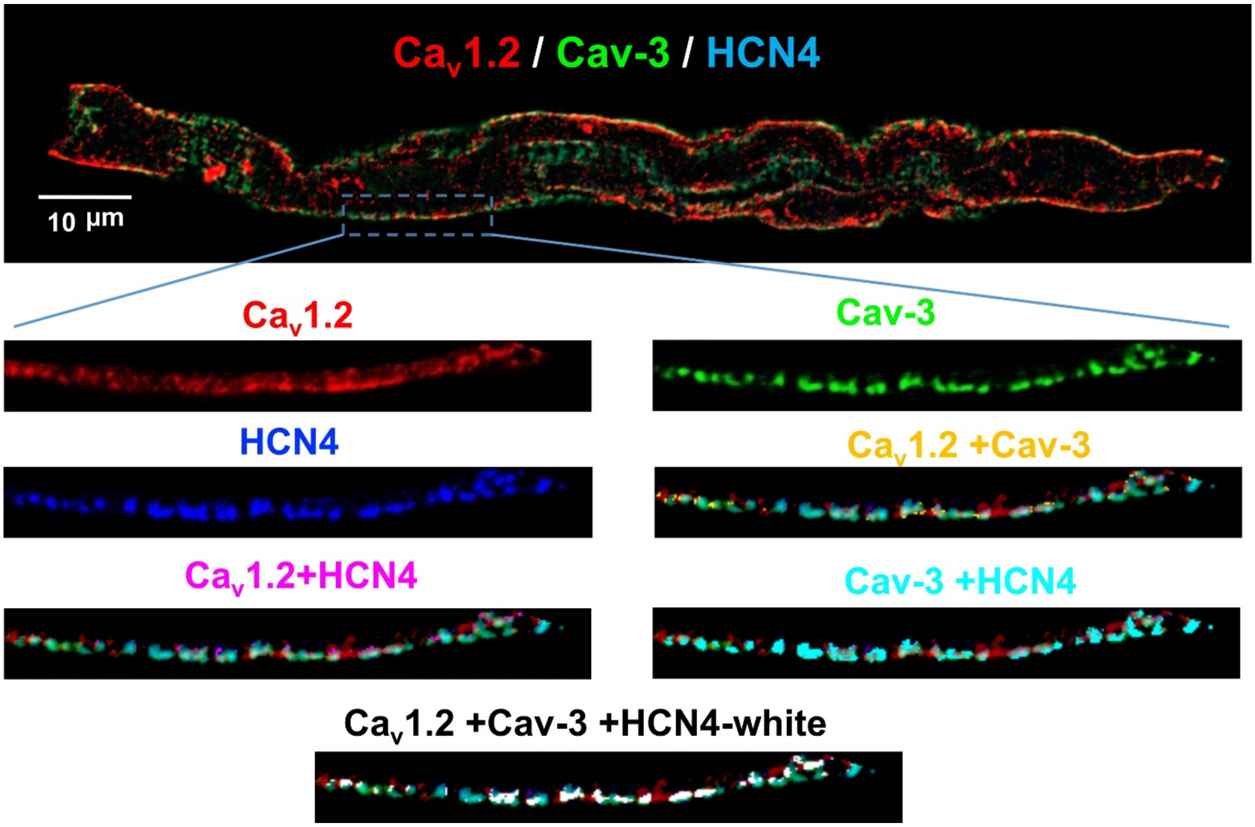


**Supplementary Figure S7.** Triple immunofluorescent staining against Cav-3 (green), L-type $Ca^{2+}$ channel isoform $Ca_V1.2$ (red) and HCN4 channels (blue). Below, single channels, paired and triple merged channels are shown from the selected membrane region of interest enlarged.

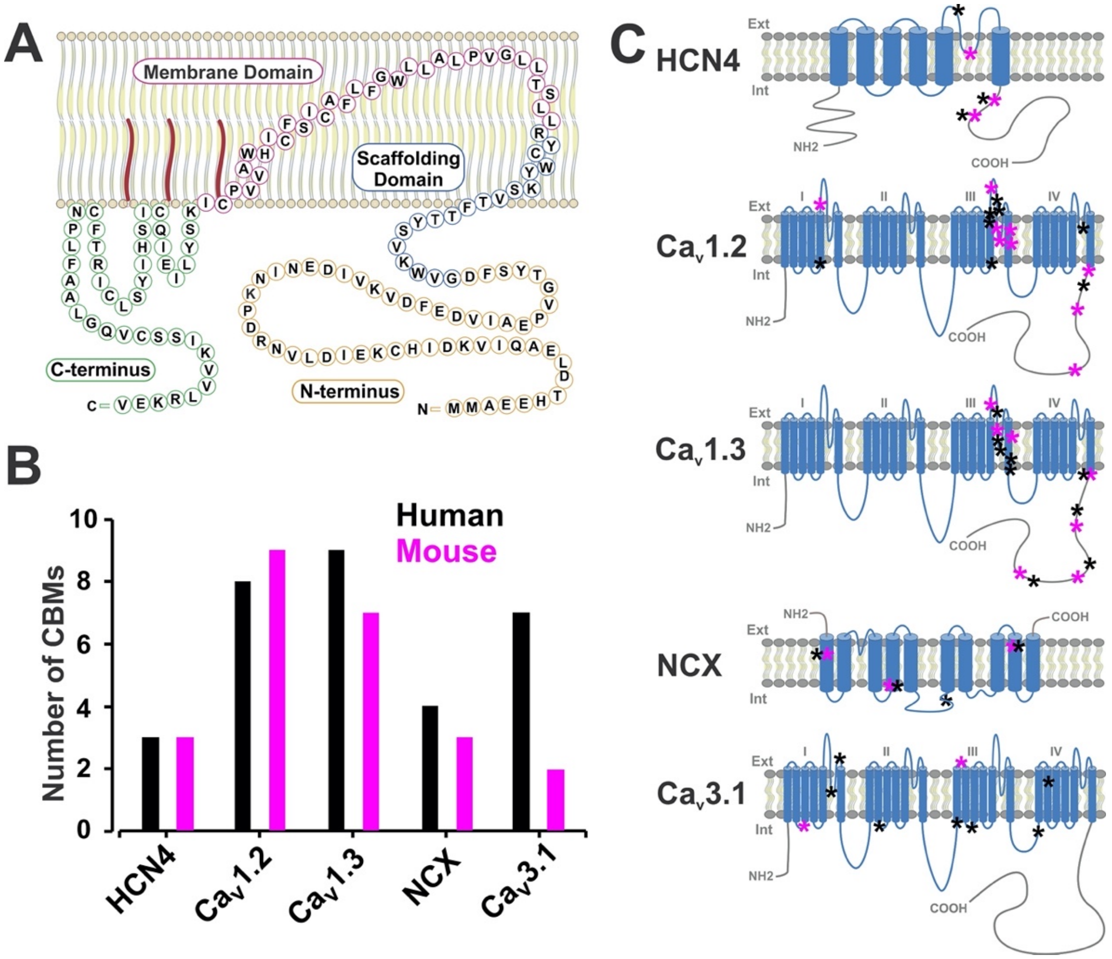


**Supplementary Figure S8.** (A) Structure of caveolin-3 (Cav-3). Cav-3 has four primary domains: $NH_2$-terminal domains (orange); scaffolding domains (blue) that form α-helices and are inserted into the membrane, with a cholesterol recognition/interaction amino acid consensus (CRAC) composed of the eight residues proximal to the membrane domain; helix-turn-helix membrane domains (fuchsia); and COOH-terminal domains (green). Reproduced from (51) with permission. (B) Number of caveolin binding motifs (CBMs) identified in the components of the membrane clock, in both mice and human. CBM analysis of the control proteins revealed one CBM in GAPDH and none in actin cytosolic proteins as well as none in $K_V11.1$ (hERG) $K^+$ sarcolemmal non-caveolar (52) membrane protein. (C) Locations of CBMs shown for the corresponding membrane protein on their structure by asterisks (black shows human CBMs, magenta shows mouse CBMs).

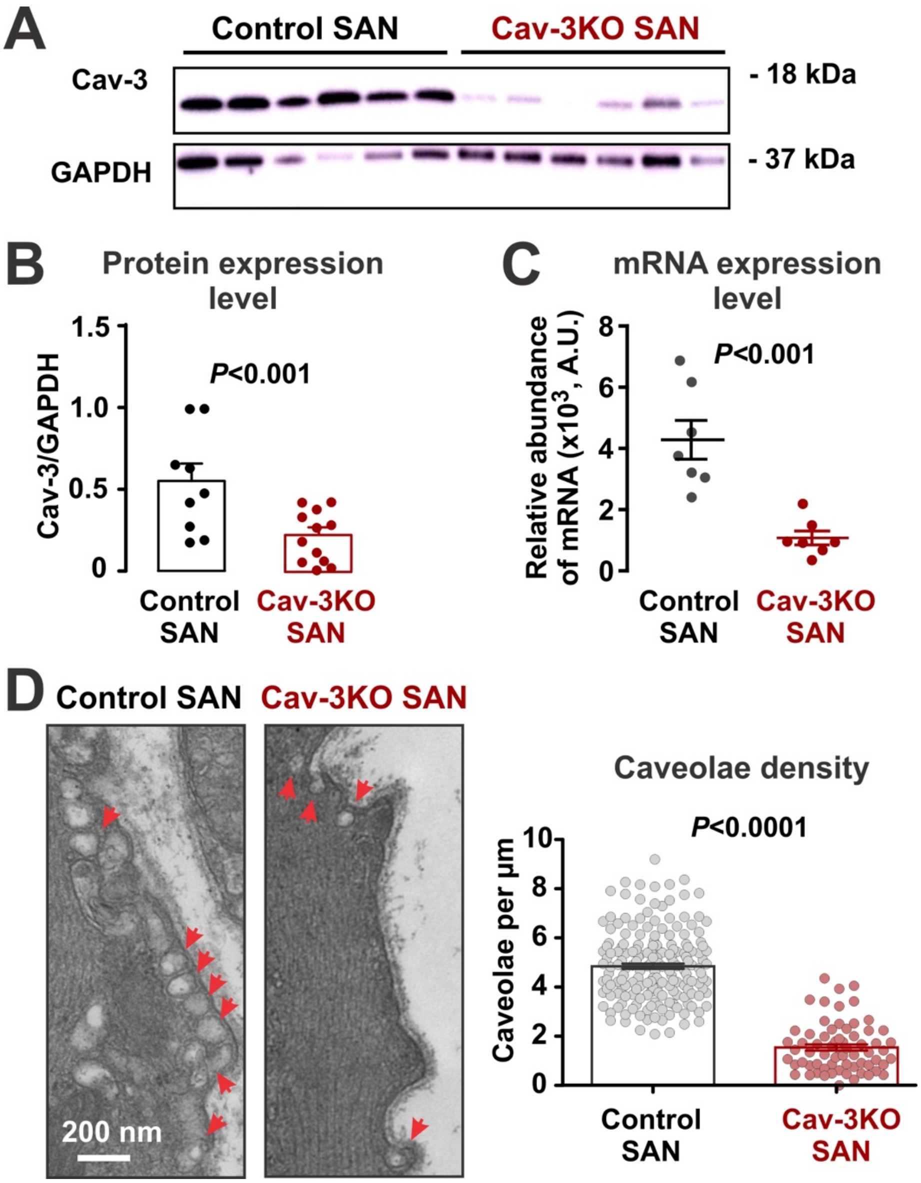


**Supplementary Figure S9.** Downregulation of caveolae structures after cardiac-specific tamoxifen-induced conditional deletion of a muscle-specific caveolar scaffolding protein, caveolin-3 (Cav3) in mice (Cav3-KO). (A) Western blot analysis of Cav-3 protein expression in control (flox-/Cre+ Cav-3KO-littermate controls after tamoxifen treatment) and Cav-3KO SAN tissue samples. (B) GAPDH-normalized Cav-3 protein expression level in control (n=9 mice) and Cav-3KO (n=11 mice) SANs. (C) mRNA expression level of Cav-3 in control (n=7 mice) and Cav-3KO (n=8 mice) SAN tissue samples. Data is reproduced from Figure 6A. (D) Representative transmission electron microscopy (TEM) photograph of control (wild type mice) and Cav3-KO SAN tissues. Subsarcolemmal 50-100 nm in diameter flask-shaped caveolae structures are shown by red arrows. Scale bar: 200 nm. On the right, caveolae densities for control (n=183 cells from 5 mice, data from Supplementary Figure S3) and Cav-3KO (n=66 images from 4 mice) SAN samples are shown. P-values were determined by Student t-test.

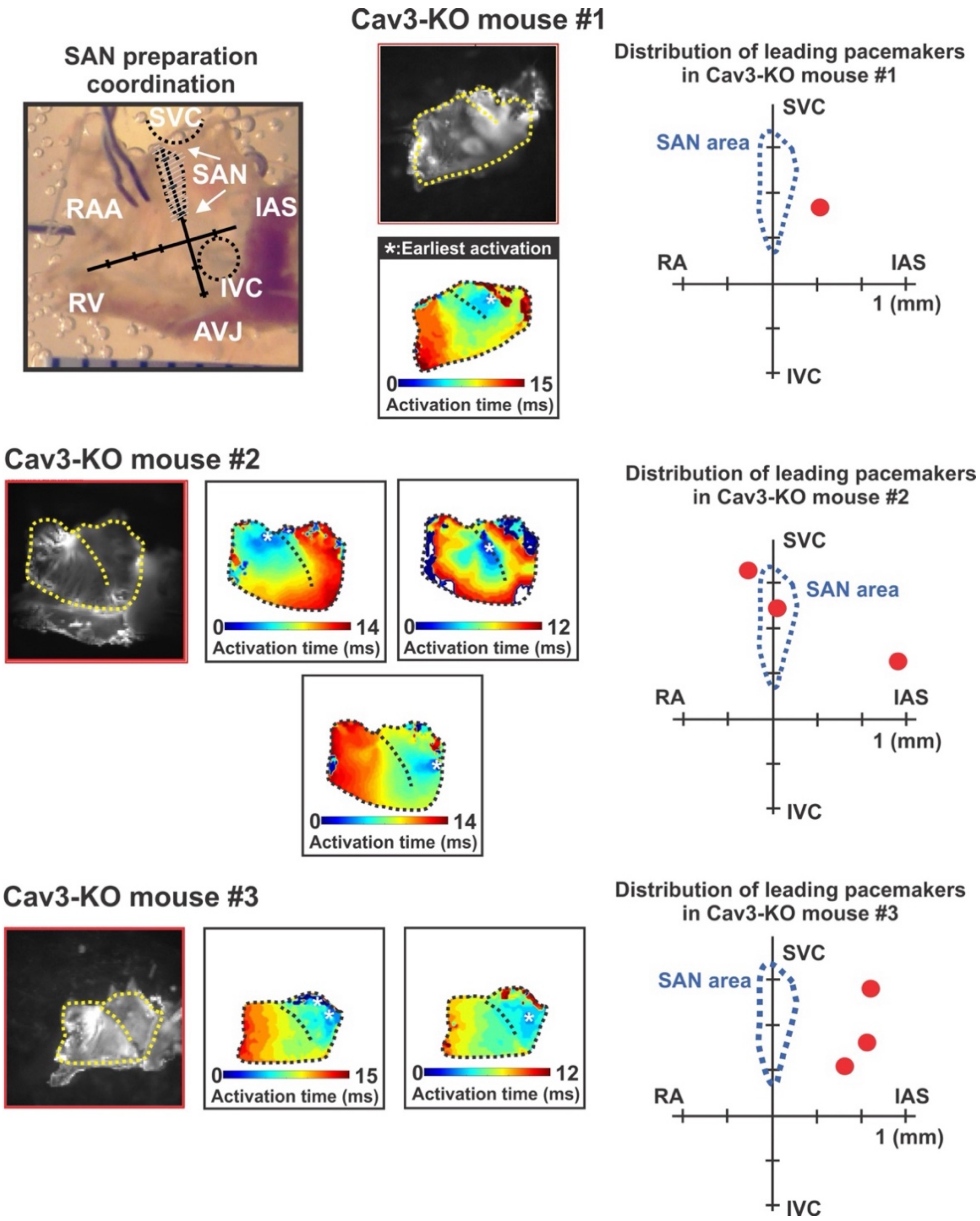


**Supplementary Figure S10. Distribution of the leading pacemaker locations in Cav-3KO mice #1-3.** Data is shown for each individual mouse (#1-3). On the top left panel, photograph of a typical SAN preparation consisting of an isolated mouse right and left atria, SAN, atrio-ventricular junction (AVJ), and the rim of the right (RV) and left (LV) ventricles. The orthogonal axes were used to plot locations of the leading pacemaker site. The axes are plotted so that they cross at the inferior vena cava (IVC); the superior to inferior direction (from the superior vena cava (SVC) to the AVJ through the IVC) is along the ordinate and the lateral to mediate direction (from the right atrial appendage, RAA, to the inter-atrial septum, IAS) is along the abscissa. On the right and below, representative activation maps are plotted for unique locations of leading pacemakers in each mouse. The maps are accompanied by enlarged illustrations of the pacemaker distribution in each mouse. The location of the leading pacemaker is shown by red circles; the SAN region is outlined by a blue dotted line.

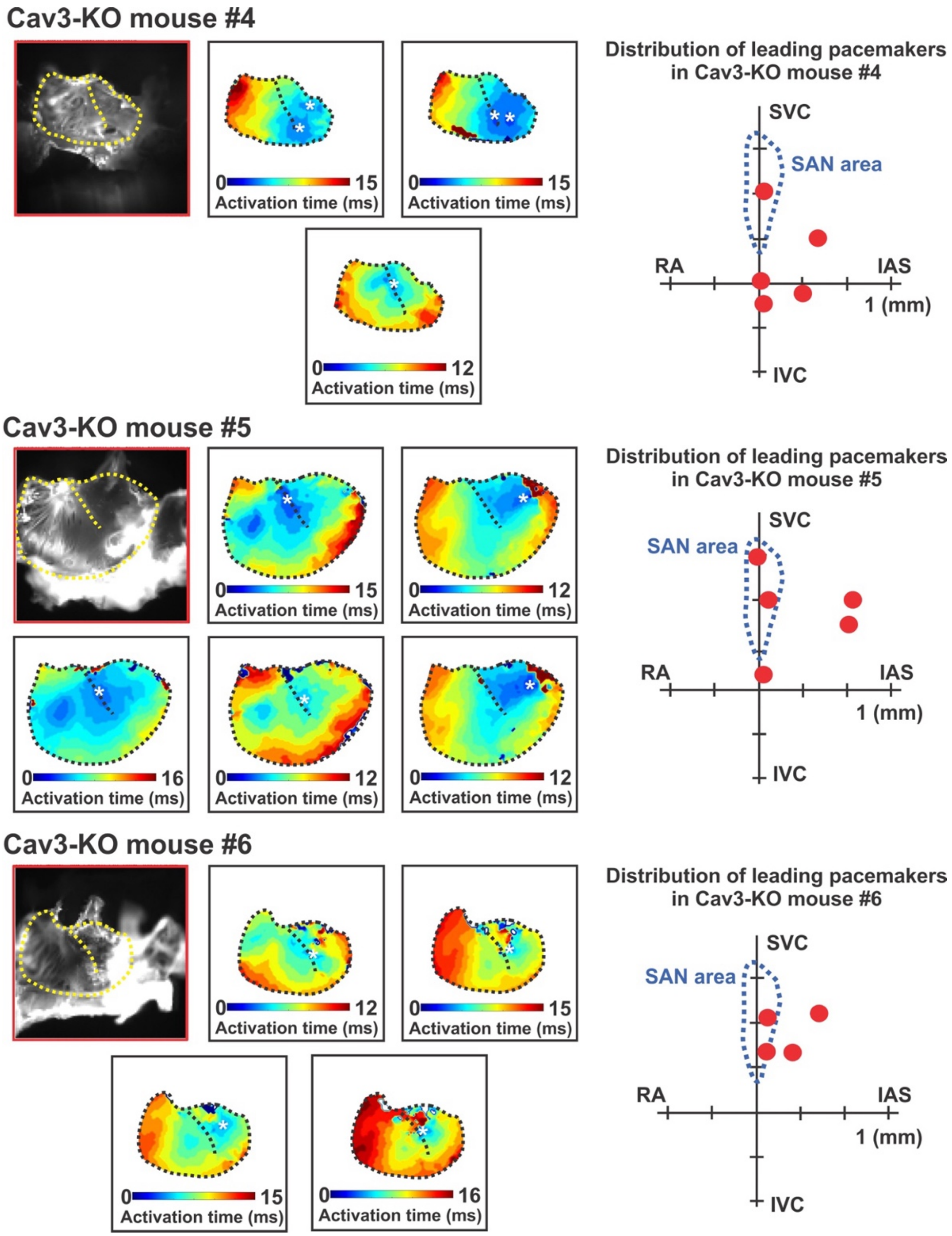


**Supplementary Figure S11. Distribution of the leading pacemaker locations in Cav-3KO mice #4-6.** Data is shown for each individual mouse (#4-6). Abbreviations and details are the same as in Figure S8.

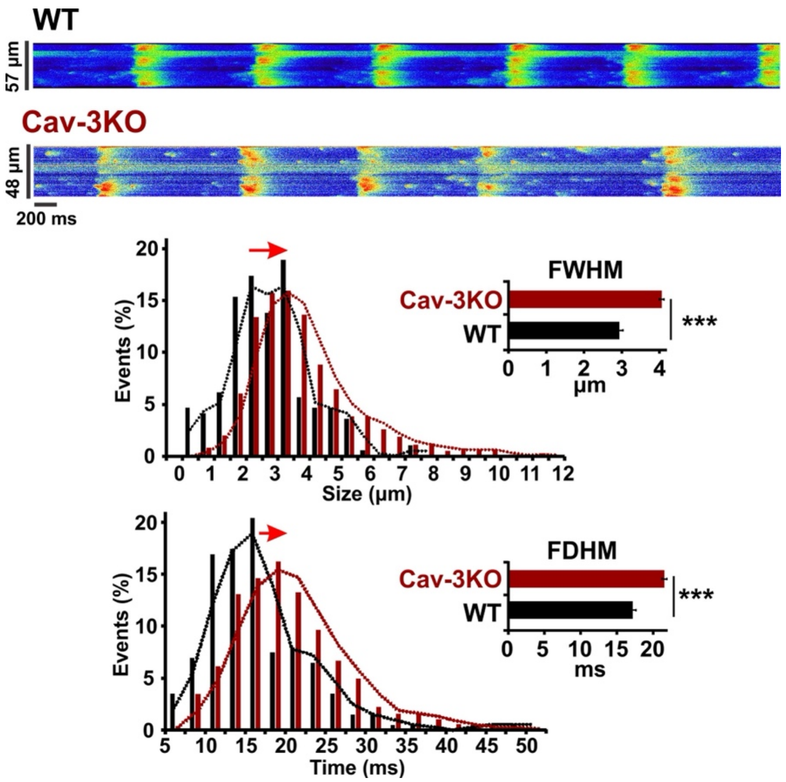


**Supplementary Figure S12.** Increase in spontaneous sarcoplasmic reticulum $Ca^{2+}$ release activity in Cav3-KO SAN myocytes. On the top, representative line scan recordings from spontaneously beating WT and Cav-3KO SAN cells. Below, $Ca^{2+}$ spark size (full width at half maximum, FWHM) and duration (full duration at half maximum, FDHM) are shown. N=13 cells for WT-SAN, and n=27 cells for Cav-3KO SAN. *** - P<0.001 vs. WT by Student t-test.

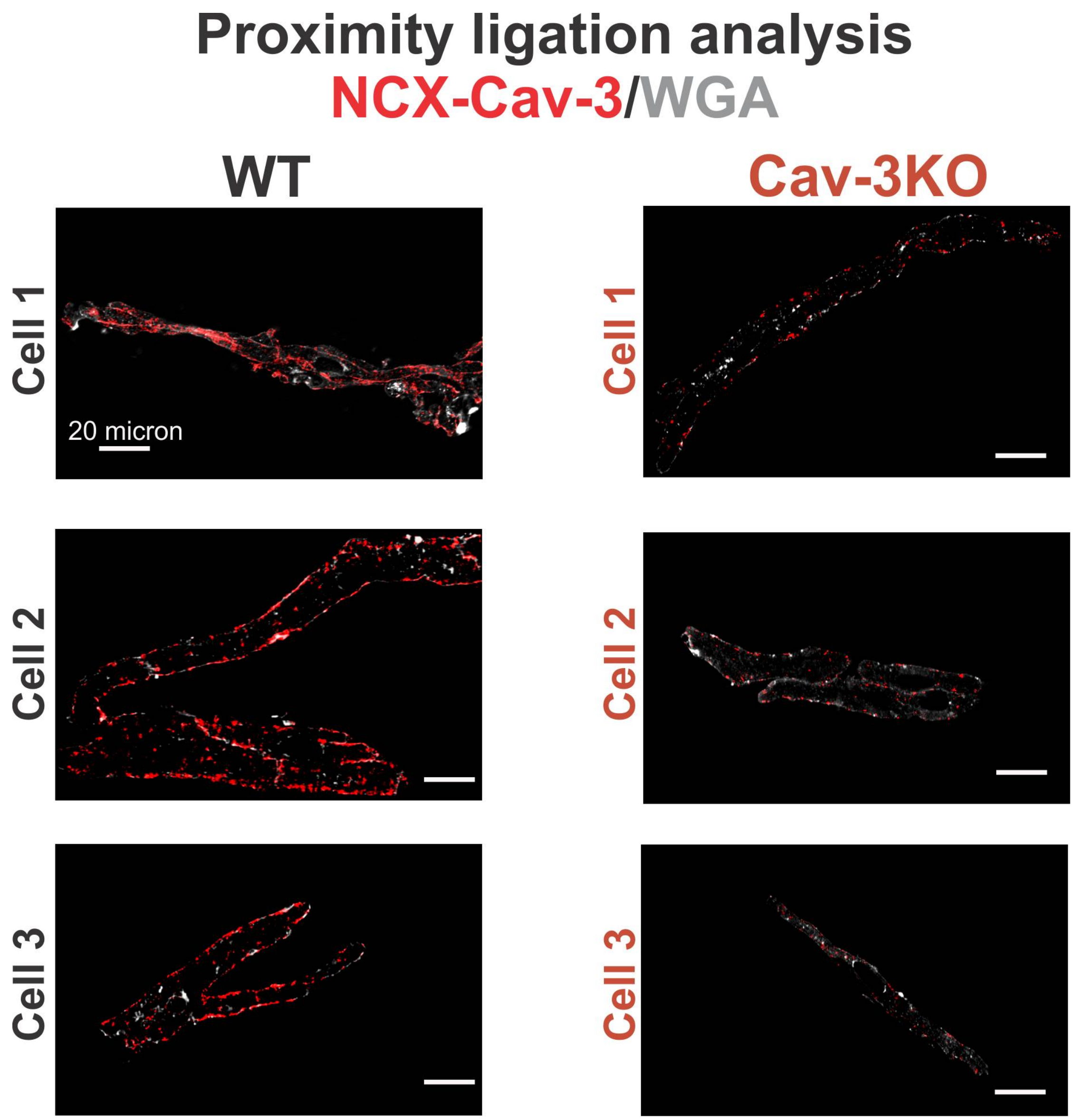


**Supplementary Figure S13.** Representative examples of proximity ligation assay staining for NCX1 and Cav-3 (in red) together with sarcolemmal membrane labeling with glycophilic lectin wheat germ agglutinin (WGA) in WT and Cav-3KO cells.

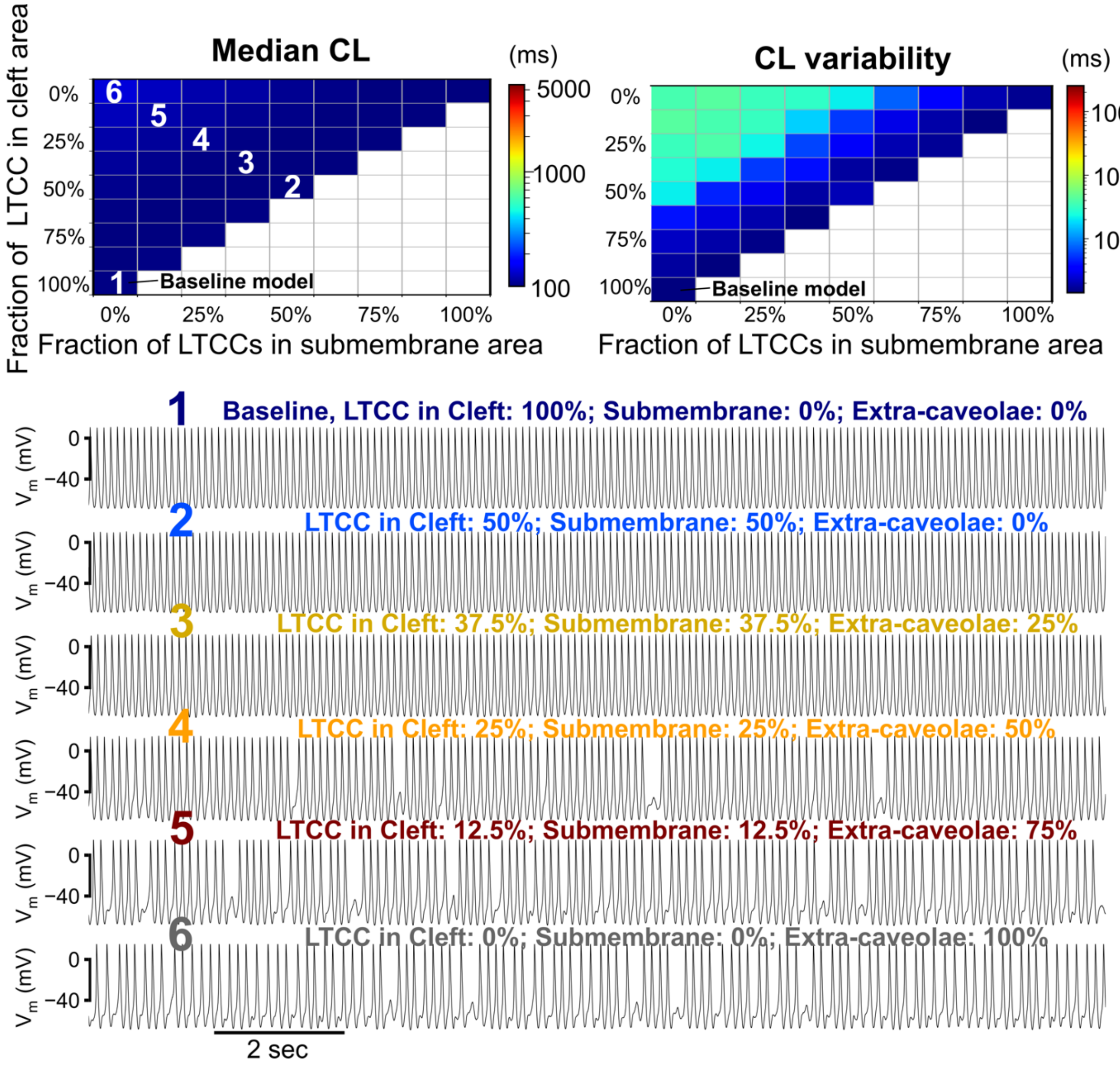


**Supplementary Figure S14. Simulated redistribution of L-type $Ca^{2+}$ channels (LTCC) away from portions of the sarcolemma facing the cleft and submembrane $Ca^{2+}$ compartments, revealing minimal impact on the cycle length (CL) and minor effects on CL variability.** *Top*: Median CL (*left*) and CL variability (*right*) with respect to varying fractions of LTCC in the cleft space (Y-axis) and the submembrane area (X-axis); the remaining fraction of LTCC was redistributed to face the cytosolic $Ca^{2+}$ compartment (extra-caveolae). *Bottom*: Time courses of $V_m$ for the baseline model and those models with various degrees of LTCC redistribution; numbers indicate the LTCC distribution parameters marked in the top left panel.

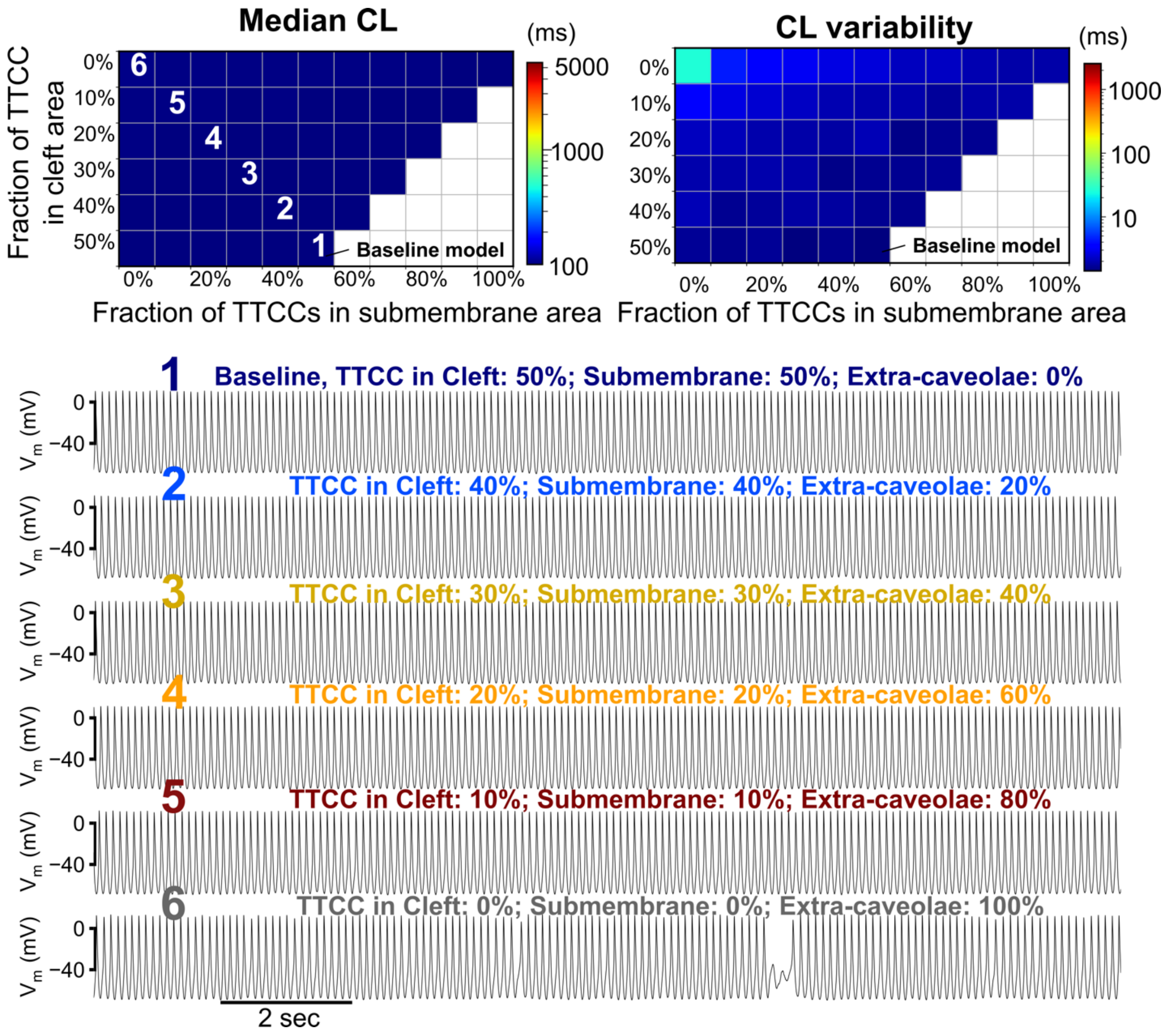


**Supplementary Figure S15. Simulated redistribution of T-type $Ca^{2+}$ channels (TTCC) away from portions of the sarcolemma facing the cleft and submembrane $Ca^{2+}$ compartments, revealing minimal impacts on the cycle length (CL) and on CL variability.** *Top*: Median CL (*left*) and CL variability (*right*) with respect to varying fractions of TTCC in the cleft space (Y-axis) and the submembrane area (X-axis); the remaining fraction of TTCC was redistributed to face the cytosolic $Ca^{2+}$ compartment (extra-caveolae). *Bottom*: Time courses of $V_m$ for the baseline model and those models with various degrees of TTCC redistribution; numbers indicate the LTCC distribution parameters marked in the top left panel.

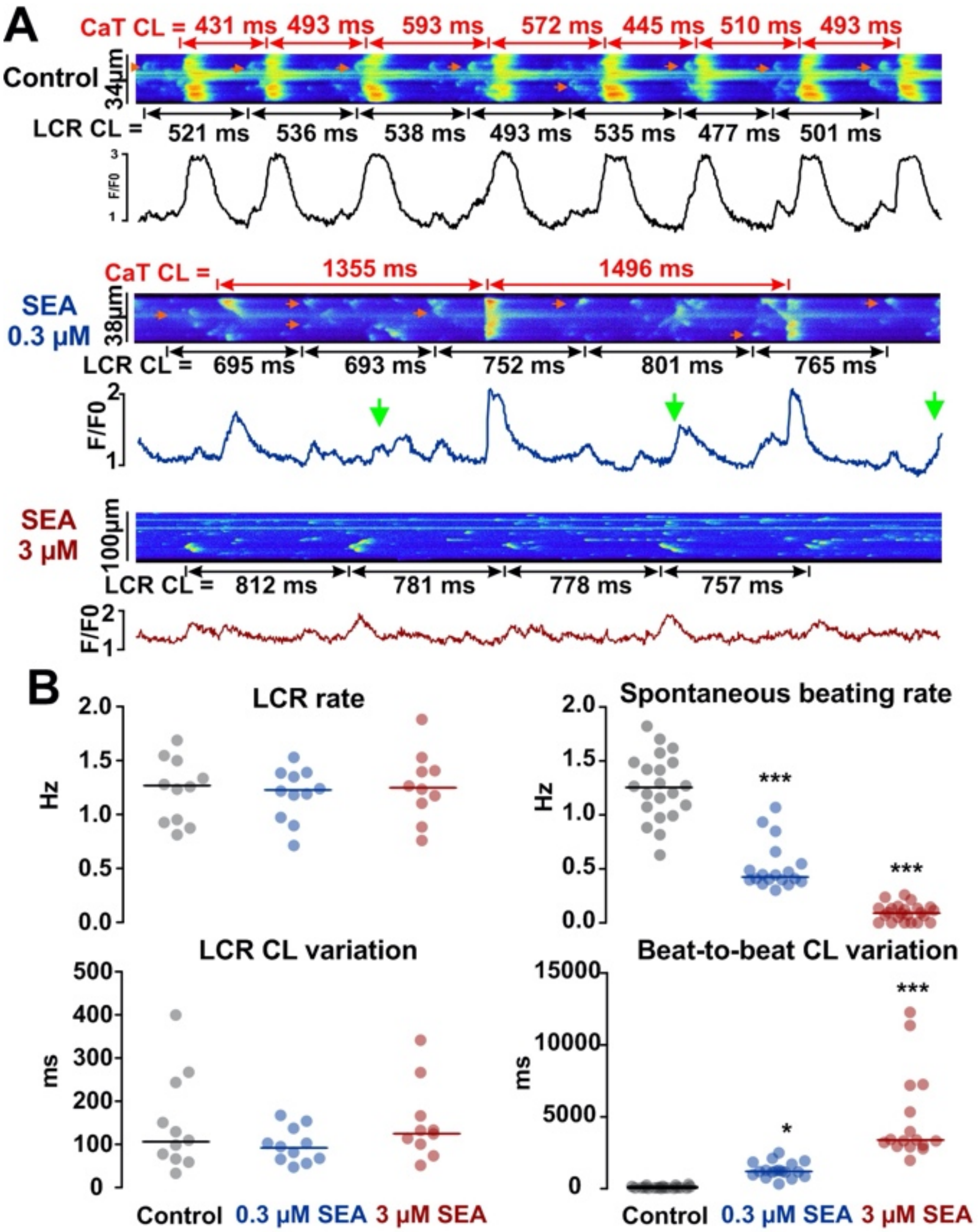


**Supplementary Figure S16. Uncoupling of membrane and $Ca^{2+}$ clocks mediated by NCX.** (A) Partial blocking of NCX in the WT isolated SAN cells using 0.3 µM SEA0400 showed similar phenotype as in Cav-3KO SAN cells, which is characterized by the missing $Ca^{2+}$ transients (CaTs) with regular local $Ca^{2+}$ release events (LCRs) and therefore significantly decreased spontaneous CaT rates as summarized in (B). Further blocking of NCX using 3 µM SEA0400 significantly exaggerated the uncoupling between the membrane and $Ca^{2+}$ clocks and resulted in dramatically slowed spontaneous CaT rate and increased beat-to-beat CaT cycle length variation. Whereas, LCR rates and cycle length during both NCX blocking conditions were not significantly different from control condition. *, *** – $P<0.05$, 0.001 by one-way ANOVA with Bonferroni correction.

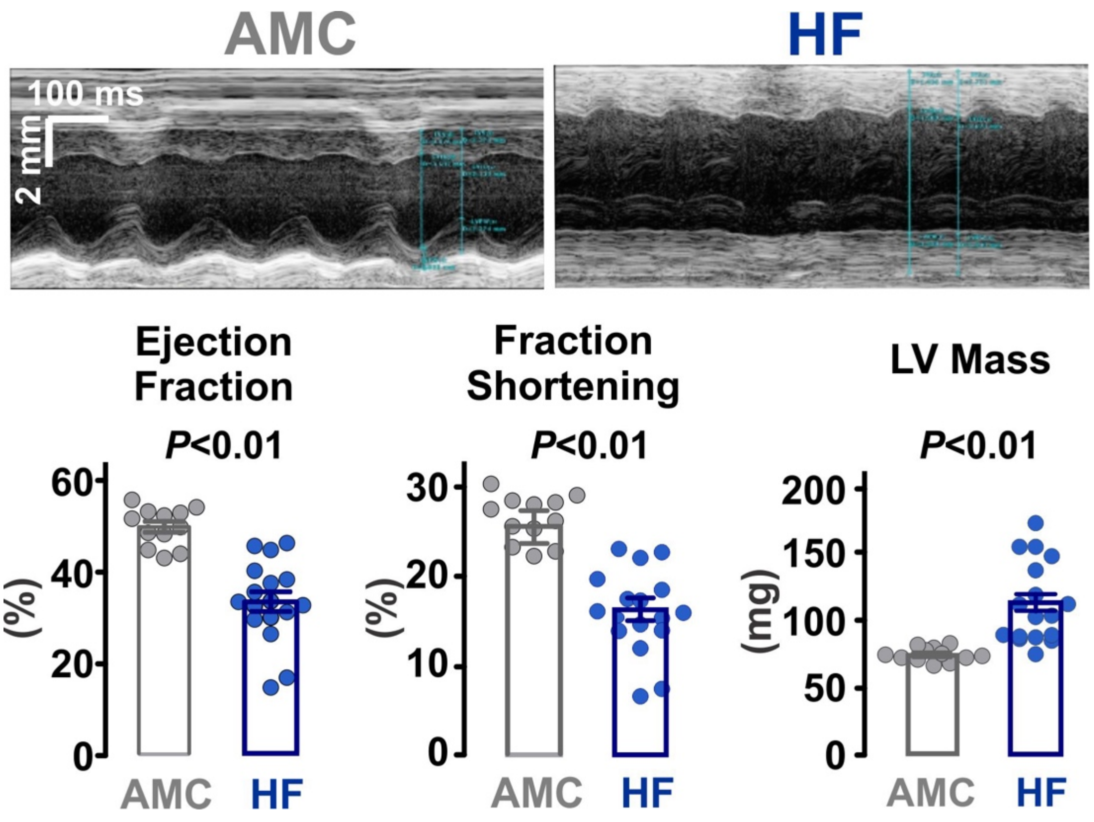


**Supplementary Figure S17.** 8-week post-myocardial infarction heart failure (HF) mice (n=17) showed a reduced ejection fraction and fraction shortening as well as an increase in left ventricular (LV) mass compared to age matched control mice (AMC, n=12). *P*-values were determined by Student t-test.

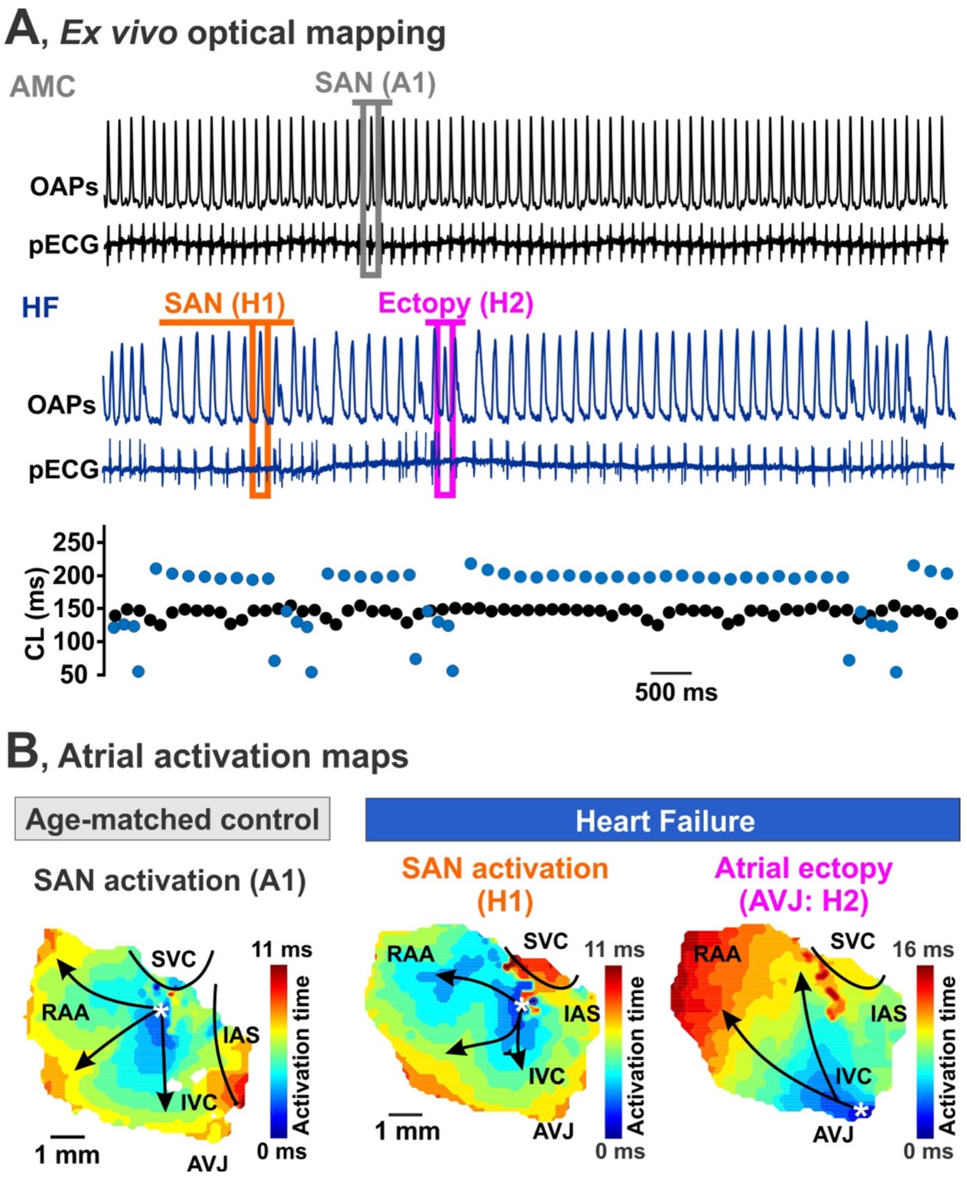


**Supplementary Figure S18.** *Ex vivo* optical mapping of atrial activity from heart failure (HF) and age-matched control (AMC) isolated atrial preparations. **(A)** Representative optical action potentials (OAPs) and pseudo-ECG (pECG) traces (*upper panels*) are shown together with corresponding beat-to-beat CL changes (*bottom panel*) for AMC and HF mice. Boxes indicate specific beats used for activation reconstruction (panel B): the black box (A1) for a typical SAN beat in the AMC mouse, the orange box (SAN H1) for a SAN beat in the HF mouse, and the purple box (ectopy from atrio-ventricular junction, AVJ, H2) for an ectopic beat in the HF mouse. (B) Representative atrial activation maps reconstructed for SAN beats (A1 for AMC and H1 for HF mice) and atrial ectopic beat (H2 for HF mouse). SVC and IVC, superior and inferior vena cava; RAA, right atrial appendage; RV, right ventricle; CT, crista terminalis; IAS, inter-atrial septum; AVJ, atrioventricular junction.

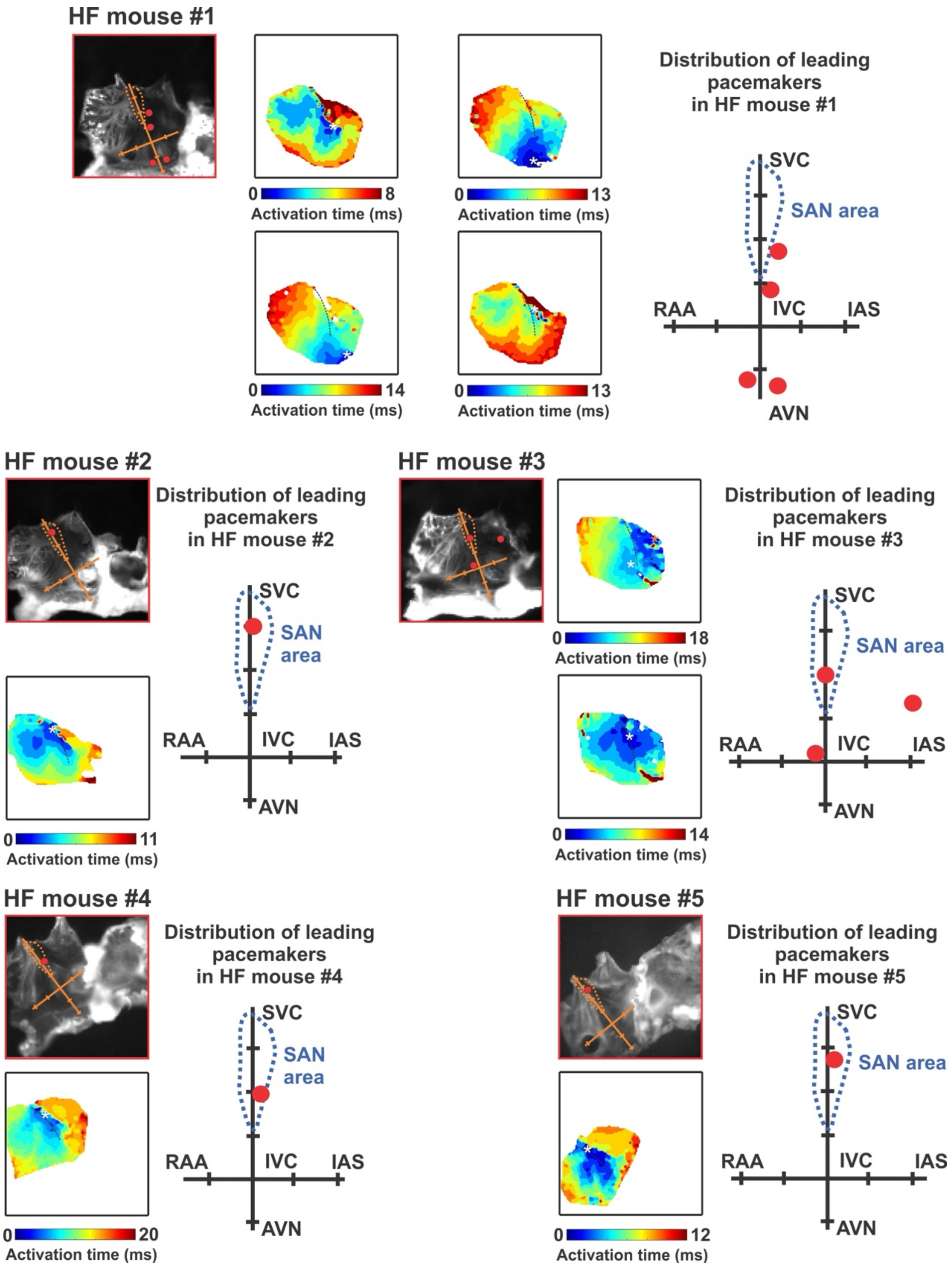


**Supplementary Figure S19. Distribution of the leading pacemaker locations in heart failure (HF) mice #1-5.** Data is shown for each individual mouse (#1-5). Abbreviations and details are the same as in Figure S8.

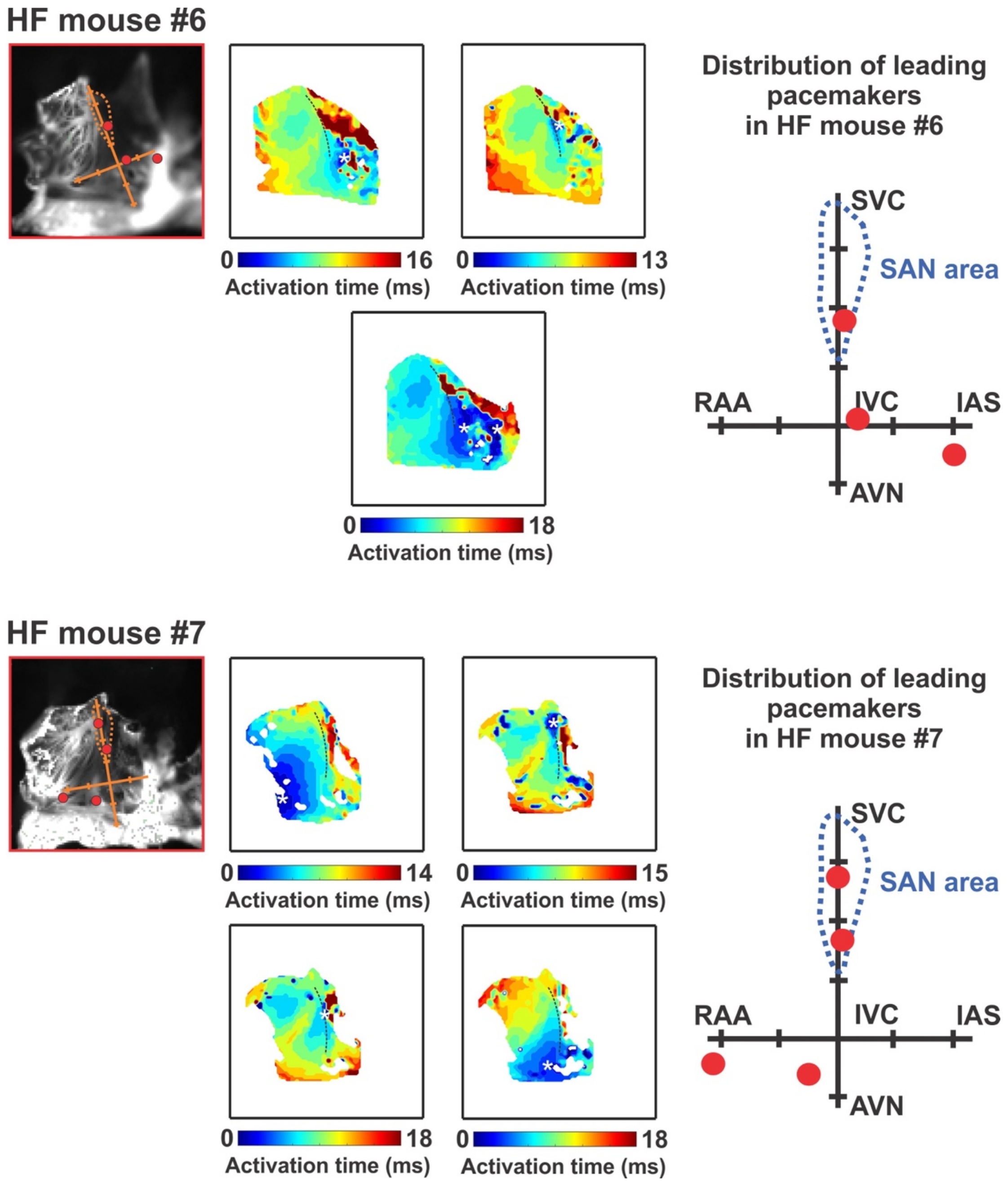


**Supplementary Figure S20. Distribution of the leading pacemaker locations in heart failure (HF) mice #6-7.** Data is shown for each individual mouse (#6-7). Abbreviations and details are the same as in Figure S8.

**Data Supplement Table I. List of used primary antibodies.**

| Target | Company and number | Host species | Dilution | Fixation type |
|---|---|---|---|---|
| Cav-3 | Abcam, ab4930 | Polyclonal rabbit | 1:500 | Methanol/PFA |
| Cav-3 | BD Biosciences, 610421 | Mouse monoclonal | 1:800 | Methanol/PFA |
| $Ca_v1.2$ | Alomone Labs, ACC-003 | Rabbit polyclonal | 1:100 | Methanol |
| $Ca_v1.3$ | Alomone Labs, ACC-005 | Rabbit polyclonal | 1:100 | PFA |
| $Ca_v3.1$ | Neuro Mab, MABN464 | Mouse monoclonal | 1:100 | Methanol |
| NCX | Swant, p 11-13 | Rabbit polyclonal | 1:200 | Methanol |
| HCN4 | Thermofisher scientific, MA3-903 | Rat monoclonal | 1:100 | PFA |
| NKA | Millipore, 05-369 | Mouse monoclonal | 1:100 | PFA |
| RyR2 | Sigma Aldrich, HPA0200228 | Rabbit polyclonal | 1:500 | Methanol |
| RyR | Thermofisher scientific, MA3-916 | Mouse monoclonal | 1:100 | Methanol |
| pSer2808 RyR | Badrilla, A01031AP | Rabbit polyclonal | 1:100 | Methanol |
| pSer2814 RyR | Badrilla, A01030AP | Rabbit polyclonal | 1:100 | Methanol |

**Data Supplement Table II. List of parameters for the 3D SAN cell model.**

| Parameter | Description | Value | Changed from Sato & Bers, 2011 (39) |
|---|---|---|---|
| $v_i$ | Local cytosolic volume | 0.75 $\mu m^3$ | Y |
| $v_s$ | Local submembrane space volume | 0.025 $\mu m^3$ | N |
| $v_p$ | Local cleft space volume | 0.00252 $\mu m^3$ | Y |

| | | | |
|---|---|---|---|
| $v_{JSR}$ | Local junctional SR volume | 0.01 $\mu m^3$ | Y |
| $v_{NSR}$ | Local network SR volume | 0.025 $\mu m^3$ | N |
| $C_m$ | Membrane capacitance | 25 pF | Y |
| $\tau_i^L$ | Longitudinal cytosolic $Ca^{2+}$ diffusion | 2.32 ms | N |
| $\tau_i^T$ | Transverse cytosolic $Ca^{2+}$ diffusion | 2.93 ms | N |
| $\tau_s^L$ | Longitudinal submembrane $Ca^{2+}$ diffusion | 1.42 ms | Y |
| $\tau_s^T$ | Transverse submembrane $Ca^{2+}$ diffusion | 1.42 ms | N |
| $\tau_{tr}$ | Junctional SR refilling time | 5 ms | N |
| $\tau_{ps}$ | Cleft to submembrane $Ca^{2+}$ diffusion | 0.022 ms | N |
| $\tau_{si}$ | Submembrane to cytosolic space $Ca^{2+}$ diffusion | 0.1 ms | N |
| $[Ca]_O$ | Extracelluar $Ca^{2+}$ concentration | 1.8 mM | N |
| $[Na]_O$ | Extracelluar $Na^+$ concentration | 136 mM | N |
| $[Na]_i$ | Intracellular $Na^+$ concentration | 10 mM | N |
| $v_{up}$ | Strength of SR $Ca^{2+}$ uptake | 1.2 μM/ms | Y |
| $g_{leak}$ | SR leak strength | $1.035\times10^{-5}$ $ms^{-1}$ | Y |
| $K_u$ | CSQN-unbound opening rate of RyR | 5.0 $ms^{-1}$ | N |
| $K_b$ | CSQN-bound opening rate of RyR | 0.005 $ms^{-1}$ | N |
| $\tau_u$ | CSQN unbinding timescale | 416.6 ms | Y |
| $\tau_b$ | CSQN binding timescale | 0.5 ms | N |
| $\tau_{c1}$ | CSQN-unbound RyR closing timescale | 2.0 ms | N |
| $\tau_{c2}$ | CSQN-bound RyR closing timescale | 0.3 ms | N |
| $K_{cp}$ | Concentration of $[Ca]_{cleft}$ for 50% of maximum closed-to-open transition rate | 5 μM | Y |
| $Max_{SR}$ | Maximum $[Ca]_{SR}$-dependent rate scaling factor | 15 | Added from Shannon et al., 2004 (49) |
| $Min_{SR}$ | Minimum $[Ca]_{SR}$-dependent rate scaling factor | 1 | |
| $EC_{50SR}$ | Concentration of $[Ca]_{SR}$ for half maximal $[Ca]_{SR}$-dependent scaling | 450 μM | |
| $N_{RyR,peri}$ | Number of RyRs in each peripheral CRU | 100 | N |
| $N_{RyR,central}$ | Number of RyRs in each central CRU | 1 | Added in new model |
| $J_{max}$ | Strength of SR $Ca^{2+}$ release from RyR | 0.2646 $\mu m^3$ $ms^{-1}$ | N |
| $P_{Ca}$ | L-type $Ca^{2+}$ channel permeability | 8.925 μM/(C·ms) | Y |
| $N_L$ | Number of LTCCs per CRU | 8 | Y |
| $k_{NaCa}$ | Strength of exchanger | 330 μM/ms | Y |
| $G_{CaBk}$ | Strength of the background sarcolemmal $Ca^{2+}$ flux | 0.005026 μM/ms | Y |
| $V_{max}$ | Strength of the sarcolemmal Ca pump | 0.0022 μM/ms | N |
| $Q_{CaP}$ | Constant | 9.4 | Y |
| $K_{mCaP}$ | Constant | 0.5 μM | N |
| H | Constant | 1.6 | N |
| $F_{NCX,cleft}$ | Fraction of NCX coupled to the cleft space | 0.5 | Added in the new model |
| $F_{NCX,sm}$ | Fraction of NCX coupled to the submembrane space | 0.5 | |

| | | | |
|---|---|---|---|
| $F_{CaT,cleft}$ | Fraction of $I_{CaT}$ in the cleft space | 0.5 | |
| $F_{CaT,sm}$ | Fraction of $I_{CaT}$ in the submembrane space | 0.5 | |
| $F_{CaP,cleft}$ | Fraction of $I_{CaP}$ in the cleft space | 0.11 | Added from Shannon et al., 2004 (49). |
| $F_{CaP,sm}$ | Fraction of $I_{CaP}$ in the submembrane space | 0.89 | |
| $F_{CaBk,cleft}$ | Fraction of $I_{CaBk}$ in the cleft space | 0.11 | |
| $F_{CaBk,sm}$ | Fraction of $I_{CaBk}$ in the submembrane space | 0.89 | |
| Gst | Sustained inward $Na^+$ current | 0.00228 nS/pF | Updated from Kharche et al., 2011 model. (41) |
| $G_{Na1.1}$ | TTX-sensitive $Na^+$ current | 0.000237 nS/pF | |
| $G_{Na1.5}$ | TTX-resistant $Na^+$ current | 0.000299 nS/pF | |
| $G_{CaT}$ | T-type $Ca^{2+}$ current | 1.0556 nS/pF | |
| $G_{CaL1.2}$ | L-type $Ca^{2+}$ current | 0 | |
| $G_{CaL1.3}$ | L-type $Ca^{2+}$ current | 0.605 nS/pF | |
| $G_f$ | Hyperpolarization-activated (funny) current | 0.233 nS/pF | |
| $G_{K1}$ | Time-independent $K^+$ current | 0.0318 nS/pF | |
| $G_{Kr}$ | Rapid delayed rectifying $K^+$ current | 0.138 nS/pF | |
| $G_{Ks}$ | Slow delayed rectifying $K^+$ current | 0.0116 nS/pF | |
| $G_{to}$ | Transient component of the 4-AP-sensitive $K^+$ current | 0.612 nS/pF | |
| $G_{sus}$ | Sustained component of the 4-AP-sensitive $K^+$ current | 0.0487 nS/pF | |
| $G_{bNa}$ | Background $Na^+$ current | 0.00526 nS/pF | |
| $G_{CaBk}$ | Background $Ca^{2+}$ current | 0.000636 nS/PF | |
| $v_{NKA}$ | $Na^+/K^+$ pump rate | 6.969 pA/pF | |
| $v_{NCX}$ | $Na^+/Ca^{2+}$ exchanger rate | 209.418 pA/pF | |
| $v_{RyR}$ | $Ca^{2+}$ release via ryanodine receptor rate | $130\times10^4$ $ms^{-1}$ | |
| $v_{SERCA}$ | Sarcoplasmic reticulum $Ca^{2+}$ pump rate | 0.04 mM/ms | |
